# Sequence based prediction of cell type specific microRNA binding and mRNA degradation for therapeutic discovery

**DOI:** 10.1101/2025.05.15.654105

**Authors:** Bhargav Kanuparthi, Sara E. Pour, Scott D. Findlay, Omar Wagih, Jahir M. Gutierrez, Rory Gao, Jeff Wintersinger, Junru Lin, Martino Gabra, Emma Bohn, Tammy Lau, Chris Cole, Andrew Jung, Albi Celaj, Fraser Soares, Rachel Gray, Brandon Vaz, Kate Delfosse, Varun Lodaya, Sakshi Bhargava, Diane Ly, Farhan Yusuf, Ken Kron, Greg Hoffman, Shreshth Gandhi, Brendan J. Frey

## Abstract

MicroRNAs and RNA binding proteins are crucial elements of post-transcriptional gene regulation, which governs the fate of mRNA molecules in the cell. However, the landscape of these regulatory interactions, particularly across different mammalian cell types, remains underexplored. We describe REPRESS, a deep learning model that predicts cell-type-specific microRNA binding and mRNA degradation directly from RNA sequence. REPRESS was trained on AGO2-CLIP, miR-eCLIP and Degradome-Seq data profiling millions of microRNA binding and mRNA degradation sites across multiple cell types in human and mouse. It reveals biology that other state-of-the-art methods did not, such as identifying repressive non-canonical miRNA target sites and decoding the regulatory effects of sequence context and miRNA binding site multiplicity. REPRESS outperforms other advanced methods and neural architectures on a comprehensive suite of seven orthogonal tasks, including identifying genetic variants that affect microRNA binding, predicting out-of-distribution data from massively parallel reporter assays, and predicting canonical and non-canonical miRNA mediated repression. To demonstrate the general utility of REPRESS, we show that it provides insights into novel biology and the design of RNA therapeutics. Code is available at : https://github.com/deepgenomics/repress

## 1 Introduction

In eukaryotes, gene expression is regulated at various stages including transcription, RNA processing (eg, splicing and RNA editing), nuclear export, and translation.

Post-transcriptional gene regulation (PTGR) is particularly critical and affects mRNA stability and localization, protein synthesis, and ultimately cell function [1]. PTGR is mediated primarily by interactions between mRNAs and two main classes of regulatory factors: RNA-binding proteins (RBPs) and microRNAs (miRNAs) [2].

miRNAs are short regulatory RNAs that guide the RNA-Induced Silencing Complex (RISC) to target mRNAs via partial sequence complementarity, leading to repression through transcript destabilization and translational inhibition [3–6]. Interactions between miRNAs and target RNAs are either canonical, with perfect base pairing within the miRNA seed region (positions 2-7), or non-canonical, consisting of imperfect seed matches and occasional extended 3’ end pairing [4, 7]. Conversely, RBPs directly interact with both pre-mRNA and mature mRNA to influence PTGR through splicing, RNA editing, transport, localization, degradation, and translation [8, 9]. Over 2,300 miRNAs [10] and 1,500 RBPs [11, 12] have been identified in humans, and collectively engage mRNA via a combination of sequence-specific, structural, and cofactor-dependent interactions [13–15]. The diversity of functions carried out by these molecules reflects the complexity of the post-transcriptional regulatory networks they participate in [16].

Despite considerable progress in elucidating the functions of miRNAs and RBPs, the full complexity of their interactions and the associated regulatory outcomes across different cellular environments remains only partially understood. Existing machine learning models for predicting miRNA-mediated target repression either rely on handcrafted features [13] or are trained on a small number of miRNAs and are limited to very short input sequences [14]. Thus, there may be additional sequence determinants of miRNA targeting that are not fully captured by existing models. Other deep learning approaches have relied on artificial sequences as negative sets for model training, restricting performance due to their limited ability to learn the endogenous biology governing miRNA targeting [17–20]. A substantial limitation of most existing approaches is that they do not take cell type into account, despite the fact that tissue and cell type specific miRNA expression and targeting are relevant to both development and disease [21–23].

Lastly, efforts to incorporate (non polyadenylation-related) site-specific endonucleolytic cleavage data have been primarily focused on identifying miRNA targets rather than predicting general mRNA degradation [24–26], or have been more descriptive in nature [6, 27–31]. Regarding data used to train models, new technologies have made transcriptome-wide identification of miRNA-specific target sites possible [32–34]. However, these methods have been applied to only a limited number of cell types that often do not include primary cell lines most relevant to disease modeling and the development of therapeutics. Furthermore, efforts to profile molecular consequences of miRNA and RBP binding, such as site-specific cleavage and degradation across mammalian transcriptomes, have been limited [27, 35].

To address these gaps in our understanding and modeling of PTGR, we generated miRNA-Seq, miRNA enhanced crosslinking and immunoprecipitation (miR-eCLIP; profiling miRNA-specific miRNA binding), and Degradome-Seq (profiling mRNA cleavage sites) data from a broad range of cell lines and tissues in human and mouse to construct a cell type specific and transcriptome-wide map of post-transcriptional gene regulation. We leverage the generated miR-eCLIP and Degradome-Seq datasets along with AGO2-CLIP data curated from the literature to develop REPRESS, a deep learning model that separately predicts cell type specific miRNA targeting and site-specific mRNA degradation for any input sequence at single base resolution. We demonstrate that REPRESS enables new discoveries by learning causal biology, outperforms existing methods on a wide variety of orthogonal tasks unrelated to its training data and involving novel sequences, and supports the design of RNA therapeutics across multiple modalities.

## 2 Results

### 2.1 Transcriptome-wide regulatory maps of miRNA binding and mRNA degradation

To establish a transcriptome-wide map of miRNA targeting, we first curated 18 existing AGO2-CLIP binding site datasets from the literature. While abundant, AGO2-CLIP data does not directly identify the specific miRNA(s) binding at a particular target. However, publicly available datasets from miRNA specific binding assays such as miR-eCLIP [34] were not very abundant, especially for cell lines and tissues of therapeutic interest. Thus, we curated three publicly available datasets, purchased access to an additional two datasets, and generated novel miR-eCLIP data from four human cell lines and two mouse tissues, for a total of 11 miRNA-specific CLIP datasets. To establish a transcriptome-wide map of site-specific mRNA degradation, we conducted Degradome-Seq, which identifies uncapped 5’ transcript fragment ends resulting from endonucleolytic cleavage, in six human cell lines and four mouse tissues, as existing Degradome-Seq datasets have primarily been generated using non-mammalian tissues [36, 37]. Collectively, these efforts resulted in the high quality compendium of datasets needed to train REPRESS, a deep learning model that separately predicts cell type specific miRNA targeting and mRNA degradation for any input sequence to enable biological insights (Fig. 1a,b).

**Fig. 1:**
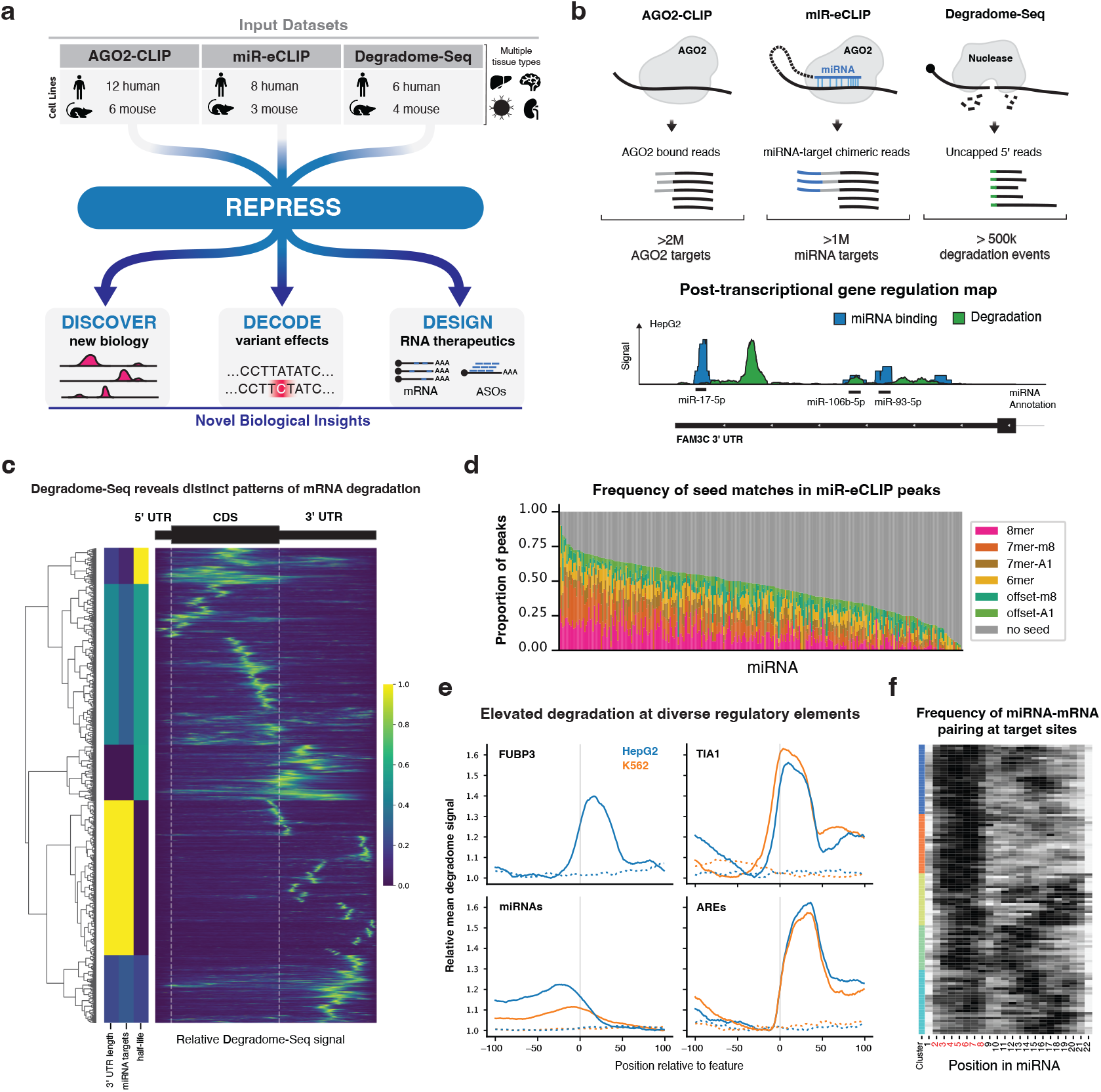
**(a):** REPRESS framework for modeling post-transcriptional gene regulation. **(b):** Summary of datasets used to train REPRESS. Top: Schematic illustration of AGO2-CLIP, miR-eCLIP, and Degradome-Seq assays used to generate REPRESS’s training dataset. Bottom: Example of a post-transcriptional gene regulation map of miRNA binding and mRNA degradation at the *FAM3C* 3’ UTR in HepG2 cells. **(c):** Relative degradome read coverage along protein-coding transcripts. Each row represents a transcript, each column corresponds to a segment of the 5’ UTR, CDS, or 3’ UTR. Values are scaled to the maximum read coverage within each transcript so that all transcripts can be visualized together (Methods). Colored vertical bars on the left show the normalized average 3’ UTR length, number of 3’ UTR miRNA targets, and half-life of transcripts for each of the five major clusters obtained by hierarchical clustering of the data, depicted by the dendrogram. **(d):** The proportion of A549 miR-eCLIP peaks containing miRNA-specific seed matches, shown by individual miRNAs. Columns are ranked by the proportion of seed matches (including offset-m8 and offset-A1). **(e):** Metaplots of average relative degradome read coverage at RBP eCLIP peak loci for FUBP3 (n = 1,021 for HepG2, no data for K562) and TIA1 (n = 479 for HepG2, n = 1,021 for K562), targets of conserved miRNAs (n = 11,585 for HepG2, n = 26,798 for K562), and UUAUUUAUU ARE motifs (n = 692 for HepG2, n = 643 for K562). Dashed lines indicate degradome read counts around control sites selected randomly from the same 3’ UTRs. Gray vertical bars represent the estimated location of the miRNA target / RBP binding site. To enable inter-cell line and target-control comparisons, degradome values for each dataset are scaled such that the minimum degradome value in the +/- 100 base window is 1.0. **(f):** LinearCoPartition analysis of the rate of miRNA-target base pairing along the entire mature miRNA. Each row represents the average number of times the base was predicted to be paired to an mRNA across all targets of a single miRNA. K-means clustering was done with Kmeans++ initialization.

Peak calling on our miR-eCLIP datasets revealed precise miRNA-specific binding at more than one million target sites across 24,000 protein-coding genes across the human and mouse transcriptomes. A high proportion (average of 44% across cell lines and tissues) of these miRNA peaks contained a seed match of the targeting miRNA identified (Fig. 1d) similar to, or exceeding, rates obtained by cross-linking ligation and sequencing of hybrids (CLASH) and miR-eCLIP [32, 34]. Furthermore, RNA co-fold analysis using LinearCoPartition [38] revealed clusters of miRNAs with distinct miRNA-target base pairing frequencies within and beyond the miRNA seed region, highlighting the molecular detail captured by miR-eCLIP (Fig. 1f, Supp Fig. 1a). Together, these results demonstrate the ability of miR-eCLIP to identify precise miRNA-specific target loci. Overall, we detected a diverse set of miRNA targets across cell lines and tissues; miRNA-seq revealed highly tissue-specific expression profiles across miRNAs (Supp Fig. 1b), and we observed a corresponding strong positive correlation between miRNA expression levels and the number of miR-eCLIP peaks identified for it in each cell line (average *R* = 0.76, *p <* 2.2 × 10^−16^; Supp Fig. 2a). While most miRNAs demonstrated preferences for targeting near the beginning and, to a lesser extent, the end of 3’ UTRs, several cell types had small clusters of miRNAs with distinct targeting patterns, including increased targeting toward the middle of 3’ UTRs (Supp Fig. 1c).

While CLIP-based approaches offer powerful detection of targets of miRNAs (and RBPs), they do not in isolation foretell the functional consequences of any particular binding event. To assess one functional aspect of PTGR, we leveraged Degradome-Seq to sequence uncapped 5’ transcript ends and identify site-specific RNA degradation events resulting from either miRNA or RBP binding, in a mechanism-agnostic fashion. This resulted in over 500,000 unique Degradome-Seq peaks. Degradation levels at individual loci demonstrated some cell type specificity (average inter-sample Pearson *R* = 0.53; *P* = 4.8 × 10^−10^; Supp Fig. 3a). Next, we performed hierarchical clustering on the relative Degradome-Seq signal at positions along protein-coding mRNAs, and consistently identified distinct clusters of transcripts, including those with: i) shorter 3’ UTRs with few miRNA targets, longer half-lives, and degradation predominantly in the CDS; and ii) longer 3’ UTRs with many miRNA targets, shorter half-lives, and highly localized 3’ UTR degradation (Fig. 1c, Supp Fig. 3c).

To further explore potential associations between different classes of RNA regulatory elements and sites of degradation captured by Degradome-Seq, we leveraged ENCODE eCLIP data for RBP binding [39] and AU-Rich Element (ARE) motifs, in addition to our miR-eCLIP peaks. Notably, we detected overall elevated degradation at each of these classes of regulatory elements, highlighting the utility of Degradome-Seq data to uncover diverse functional post-transcriptional regulatory elements (Fig. 1e, Supp Fig. 3b). Among all RBPs with available ENCODE eCLIP binding data, we detected elevated Degradome-Seq read coverage most prominently for FUBP3, PUM2, TIA1, and UPF1. This finding is consistent with UPF1 recently being identified as having the most enriched binding (among all RBPs analyzed) near capped 5’ ends in human 3’ UTRs [40], while also suggesting that other RBPs such as FUBP3, PUM2, and TIA1 may have underappreciated roles in regulating mRNA decay through endonucleolytic cleavage. Consistent with observations of occasional miRNA-directed target cleavage [35, 41], we also detected a more modest increase in Degradome-Seq read coverage around miRNA targets overall, and site-specific degradation for individually validated instances of miRNA directed endonucleolytic mRNA cleavage [35] (Supp Fig. 3d). Lastly, we detected robust elevation in Degradome-Seq read coverage around AREs, suggesting these elements may direct degradation by endonucleases. Intriguingly, these findings also offer insight into potential mechanistic differences between mRNA cleavage events associated with miRNA or RBP binding, as inferred cleavage sites occurred most frequently at miRNA targets, and downstream of AREs and RBP binding sites. In summary, our results suggest site- and tissue-specific mRNA cleavage is an integral part of PTGR mediated by diverse molecular interactions. Together, these miRNA binding and mRNA degradation datasets provide valuable biological insights and facilitate the training of a robust deep learning model.

### 2.2 REPRESS: A single-base resolution, cell type specific model of miRNA binding and mRNA degradation

REPRESS makes use of a novel 133M parameter neural network that takes an input nucleotide sequence and separately predicts miRNA binding (Supp Table 1) and mRNA degradation (Supp Table 2) (degradome-seq read coverage) across 29 and 10 cell lines/ tissue types, respectively, at single base-pair resolution (Fig. 2a, Supp Fig. 4). REPRESS uses a novel neural architecture inspired by ConvNeXt [42], including residual blocks with gradually increasing receptive fields that capture long range-dependencies by incorporating 12.5 kb of context around the query sequence, which is sufficient to model information across the entire mature mRNA length for 99.2% of all protein-coding mRNAs in the human and mouse genomes.

**Fig. 2:**
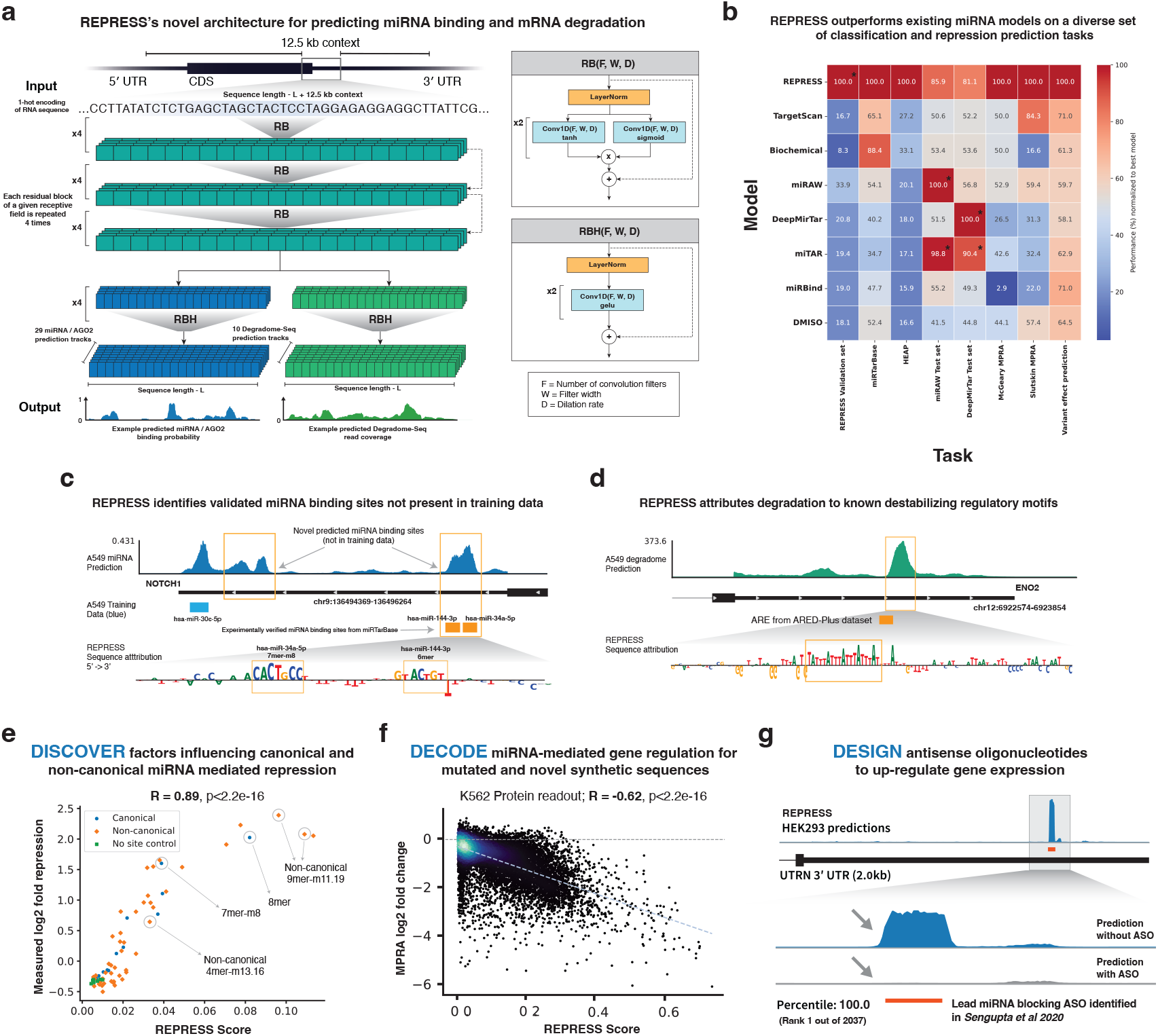
**(a):** REPRESS’s ConvNeXt inspired architecture and prediction pipeline. REPRESS takes RNA sequence as input along with bases from a 12.5 kb context window around the query sequence. The one-hot encoded sequence is then passed into a dilated convolutional neural network architecture with residual blocks and skip connections. Information from selected base layers is pooled and passed to specific output layers for generating the miRNA and degradome predictions respectively. **(b):** Heatmap comparing the performance of REPRESS to other popular miRNA binding models over a comprehensive suite of miRNA related tasks including predicting miRNA binding sites from miRTarBase, HEAP, miRAW and DeepMirTar test set; predicting miRNA mediated repression of synthetic sequences from the McGeary and Slutskin MPRA and predicting the consequences of miRNA binding altering variants. * indicates non-orthogonal test data, i.e., that test data came from the same experimental setup as the training data. **(c):** REPRESS’s miRNA prediction on the 3’ UTR of *NOTCH1*. The figure illustrates one miRNA binding site of hsa-miR-30c-5p in REPRESS’s training dataset, but the model predicts four additional peaks not present in the training dataset. Two of these corresponded to the binding sites of hsa-miR-34a-5p and hsa-miR-144-3p which are validated binding sites from miRTarBase. The sequence attribution methods on the miRNA prediction revealed the identity of the seed sequence of the miRNA bound resulting in the predicted peaks. **(d):** REPRESS’s degradome predictions on the 3’ UTR of *ENO2*. The strongest predicted peak corresponded to an ARE site identified in the ARED-Plus dataset. Sequence attribution methods confirmed that the repeating overlapping ATTTA motif is driving the prediction of the strongest degradation peak. **(e):** REPRESS learns underlying biology governing canonical and non-canonical miRNA mediated repression for 68 engineered target sites in a fixed sequence context [52] with a Pearson R=0.89. REPRESS score is defined as the average REPRESS miRNA prediction over the entire query sequence. **(f):** REPRESS decodes the effect of miRNA binding altering variants and accurately predicts miRNA mediated repression for mutated and novel out-of-distribution sequences from the Slutskin MPRA [51] (n=12,545) with a Pearson R=-0.62. REPRESS score is defined as the average K562 miRNA prediction across the entire MPRA construct. **(g):** REPRESS can efficiently design up-regulating ASOs ranking a validated miRNA target site blocking ASO from Sengupta et al. [73], 1 out of the 2,037 possible ASOs that could target the 3’ UTR of the *UTRN* gene.

To evaluate REPRESS’s performance on miRNA target detection, we applied it to a validation set (4-fold cross validation) containing 25% of transcripts from the human and mouse transcriptome. REPRESS achieves an average 4-fold cross validation Area under Receiver Operating Characteristic (AUROC) curve of 0.88 (Supp Fig. 5a) and an average Spearman correlation of 0.60 when predicting degradation events (Supp Fig. 5b). To further test REPRESS’s performance on orthogonally generated sets of experimentally validated miRNA targets we leveraged data from miRTarBase [43] and DianaTarBase [44] (Methods). REPRESS accurately predicted cell-type specific miRNA targets with an average AUROC curve of 0.78 including those that were not present in the training dataset (Supp Fig. 5c).

REPRESS’s ConvNeXt-inspired dilated CNN architecture exhibits strong scaling laws and outperforms other state of the art architectures that we trained on the same data. REPRESS’s accuracy, measured by performance on the validation set of both miRNA binding and degradome tasks, scales smoothly and strongly with the number of parameters in the model (Supp Fig. 5f). This is consistent with the scaling laws observed in deep learning models used for natural language processing [45] and biological sequence modeling [46]. When the model capacity is kept fixed at a large size, we also found that models jointly trained on both miRNA and Degradome-Seq data performed better than models trained on either individual dataset. We evaluated the generalization performance of REPRESS’s ConvNeXt-inspired architecture against other widely used deep learning architectures that we trained on the same data, including a vanilla CNN, Transformers [47], Mamba (a state-space model) [48], and a vanilla dilated CNN (without the ConvNeXt inspired residual block), all utilizing the same 12.5 kb context sequence. REPRESS’s ConvNeXt-inspired architecture demonstrated superior generalization, outperforming the Mamba model, the next-highest performing architecture, by 10%, and the vanilla dilated CNN by 17% (Supp Fig. 5g).

To evaluate and benchmark REPRESS’s predictive capabilities against existing models of miRNA targeting, we compiled a comprehensive suite of diverse tasks related to the classification and functional impact of miRNA binding sites. We evaluated performance on all tasks for the following models: TargetScan [13], Biochemical model [14], miRAW [49], DeepMirTar [17], miTAR [18], miRBind [19] and DMISO [20]. Tasks included: 1. Identifying experimentally validated miRNA binding sites from i) REPRESS’s test set, ii) the miRAW test set [49], iii) the DeepMirTar test set [17], iv) miRTarBase, and v) HEAP binding data [50]; 2. Predicting miRNA-mediated repression of synthetic sequences from two Massively Parallel Reporter Assays (MPRAs) designed to interrogate miRNA targeting [51, 52]; and 3. Predicting the impact of miRNA binding-altering variants curated from the literature. REPRESS substantially outperformed the other models on all tasks, with the exception of tasks where the evaluation data was from the same distribution used to train the corresponding model, as indicated by asterisks (*) (Fig. 2b, Supp Fig. 6). Specifically, while miRAW performed best on its own test set, it performed poorly on the DeepMirTar test set, and vice versa, suggesting overfitting to their respective datasets. For both the miRAW and DeepMirTar tasks, REPRESS was still the best performing independently trained model, demonstrating its ability to generalize across diverse miRNA datasets.

For any arbitrary sequence of interest, REPRESS separately predicts the likelihood of miRNA binding and Degradome-seq read coverage. In addition to these predictions, REPRESS can be used to query all possible single nucleotide variants along a sequence to predict precisely which nucleotides are important for miRNA binding or mRNA degradation. As an example of miRNA binding prediction, REPRESS accurately predicted a hsa-miR-30c-5p binding site in the 3’ UTR of human *NOTCH1* present in the training data, but also two additional upstream binding sites (for hsamiR-34a-5p and hsa-miR-144-3p) not present in the training data but independently experimentally validated [53], with REPRESS attributing this binding to seed matches for the corresponding miRNAs (Fig 2c). As an example of degradome read coverage prediction, the strongest REPRESS degradome peak in the ENO2 3’ UTR overlapped an AU-rich element (ARE) present in the ARED-Plus database [54], with REPRESS accurately attributing this signal to an (ATTT)_4_ repeat characteristic of AREs (Fig 2d).

More generally, REPRESS can be deployed as a powerful tool for biological discovery, decoding the impact of genetic variants, and designing genetic medicines. Highlights of these capabilities include demonstrated learning of diverse canonical and non-canonical [7] modes of binding used by endogenous miRNAs and accurate quantitative prediction of subsequent repression for a subset of sequences screened from the McGeary et al. MPRA [52] (Fig. 2e), strong correlations between predicted miRNA binding and observed repression for mutated derivatives of miRNA target sites, even at the level of protein (Fig. 2f), and extremely efficient design of validated ASOs to sterically block miRNA binding sites and increase expression of therapeutic targets (Fig. 2g).

### 2.3 REPRESS enables discovery of post-transcriptional regulatory biology

We next wanted to explore how REPRESS could be leveraged to discover both validated and novel aspects of endogenous biology. We first utilized REPRESS to generate sequence attributions for strongly predicted target sites of highly expressed miRNAs to assess to what extent each target site position/base contributed to miRNA binding. These sequence attributions were computed by running REPRESS on in silico mutated sequences (called in silico mutagenesis, ISM) and were represented by positional scores along the mature miRNA sequence for interpretability. As expected, REPRESS generally attributed miRNA binding as being most prominently driven by the seed region spanning positions 2-7 (Fig. 3a). Strikingly, REPRESS identified that hsa-miR-148a-3p had a shifted attribution pattern that spanned positions 3-8, instead of 2-7, corresponding to an offset-6mer site type. While it is well established that offset-6mers are generally less potent than their other canonical site type counterparts [14, 55], REPRESS offered a hypothesis that offset-6mer targets are of increased functional relevance for the binding of hsa-miR-148a-3p specifically. To test this hypothesis and explore the validity of REPRESS’s prediction, we analyzed both the underlying miR-eCLIP data and orthogonal cross-species conservation data. We found that relative to typical miRNAs (e.g. hsa-let-7a-5p), hsa-miR-148a-3p engaged offset 6mer target sites more frequently, and these targets were conserved across vertebrates at a significantly higher rate than 6mer targets (at a rate similar to that observed for 8mer targets) (Fig. 3a, Supp Fig. 7b).

**Fig. 3:**
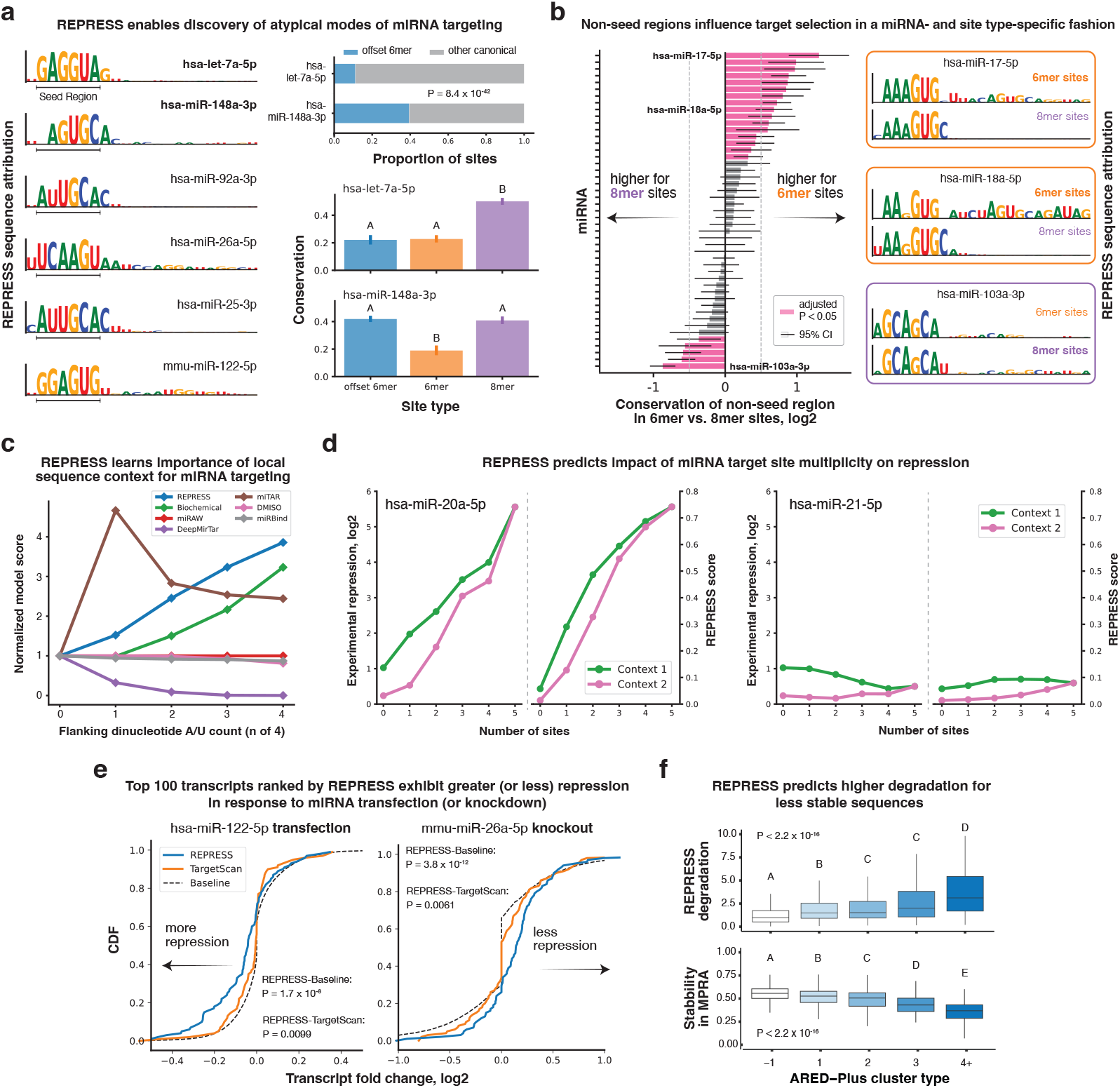
**(a):** Left: Sequence attribution base heights show the relative impact on miRNA binding that REPRESS assigned to each position (represented in terms of the corresponding miRNA position/base). Right, top: Proportion of miR-eCLIP peaks containing offset-6mers for hsa-let-7a-5p and hsa-miR-148a-3p. P value is the result of a two proportion Z-test comparing the proportion of offset 6mer sites between hsa-let-7a-5p and hsa-miR-148a-3p. “Other canonical” includes 6mer, 7mer-A1, 7mer-m8, and 8mer sites. Bottom: Conservation is the proportion of seed bases with PhyloP 100-way *>* 3. Different letters indicate statistically significant groupings of site types (*P <* 0.05) after pairwise two proportion Z tests and Benjamini-Hochberg adjustment for multiple hypothesis testing. **(b):** Left: miRNAs are ranked by the odds ratio for non-seed region conservation rates in 6mer targets relative to 8mer targets. P values and confidence intervals (CI) are the result of Benjamini-Hochberg adjusted two-sided Fisher Exact Tests. Right: REPRESS sequence attributions for 6mer and 8mer targets of highly expressed miRNAs with substantial differences in non-seed region conservation between 6mer and 8mer targets. Remaining miRNAs are shown in Supp Fig. 7c. **(c):** REPRESS learns the importance of the local AU content of the dinucleotides flanking 8mer targets. For each model in the line plot, the mean score across all sites is normalized to sites with no A or U bases in either of the dinucleotides flanking positions 1-8. **(d):** REPRESS predicts the impact of miRNA target site multiplicity on repression. Experimental repression (log_2_ fold-change) values and corresponding REPRESS predictions (average score across sequence) are shown for 3’ UTRs designed to have a variable number of miRNA targets embedded in two different sequence contexts. Examples of two miRNAs for which increasing the number of target sites differentially impacts observed repression are shown. **(e):** CDF plots showing the effect of miRNA modulation on transcript fold-change (log_2_) in two experiments. Left: hsa-miR-122-5p transfection. Right: mmu-miR-26a-5p knockout. In both panels, the top 100 ranked transcripts by REPRESS (blue line) and TargetScan (orange line) are compared to the baseline of all transcripts analyzed (dashed black line, n = 19,195 for hsa-miR-122 experiment, n = 20,810 for mmu-miR-26a experiment). P values are the result of two-sided KS tests between REPRESS and TargetScan sets for each experiment. **(f):** Boxplots for relative REPRESS degradation predictions (normalized to median value for ARED-Plus cluster type = −1 are shown above boxplots for experimentally determined relative stability values for sequences stratified by ARED-Plus cluster type. The clusters are based on the number of overlapping AUUUA pantamers a sequence contains, with a higher number generally corresponding to more degradation and decreased stability. Horizontal lines indicate (from bottom to top) the first quartile, second quartile (median), and third quartile. Whiskers extend 1.5 × IQR below Q1 and 1.5 × IQR above Q3, where “IQR” = interquartile range. “ARED”: AU-Rich Element Database.

REPRESS also predicted an influence of positions outside of the seed region for some miRNAs. This prompted us to generate REPRESS sequence attributions for different site types for each miRNA. When comparing generally weaker canonical 6mer sites to generally stronger canonical 8mer sites, REPRESS attributed increased importance to non-seed region positions for 6mer targets for many miRNAs. One possible interpretation of this finding is that the weaker seed region base pairing offered by 6mers is often augmented by additional extended miRNA-target interactions, whereas stronger 8mers are less dependent on such interactions for achieving higher affinity binding [56]. To obtain independent evidence for this hypothesis, we compared rates of conservation across vertebrates for non-seed regions of both 6mer and 8mer target sites. Nearly half (21 of 47 = 45%) of miRNAs eligible for analysis had significantly higher rates of conservation at non-seed positions in 6mer targets relative to 8mer targets (mean odds ratio = 1.64), despite the vast majority of miRNAs having lower rates of conservation within the seed region for 6mer targets relative to 8mer targets (Fig. 3b). Strikingly, for all of the highly expressed miRNAs with substantially higher non-seed region conservation for 6mer targets, REPRESS also attributed increased importance to non-seed positions for 6mer targets. In contrast, for the miRNA with the highest non-seed region conservation for 8mer targets (hsa-miR-103a-3p), REPRESS correspondingly attributed increased importance of non-seed positions for 8mer targets (Fig. 3b, Supp Fig. 7c).

In addition to these novel insights into miRNA biology, we also confirmed that REPRESS has learned well established general rules of miRNA targeting: The ranking of site types by mean REPRESS score matches the relative strength of each site type, 8mer *>* 7mer-m8 *>* 7mer-A1 *>* 6mer [55] (Supp Fig. 7e), and there is a strong positive linear relationship between the AU content of the dinucleotides flanking the 8mer seed region and mean REPRESS score (Fig. 3c, Supp Fig. 7d). The latter result is consistent with independent observations of high AU contexts permitting stronger binding which has been attributed to secondary structure-dependent target accessibility [14, 57]. Notably, among all of the other models tested, only the biochemical model [14] (mostly) demonstrated the expected linear relationship between AU content and target strength (Fig. 3c).

The biology of miRNA-mediated regulation is complex and nearby binding sites can act additively, synergistically, or competitively [51, 57, 58]. To determine whether REPRESS’s large receptive field enables it to account for sequence context and binding site multiplicity, we examined data for a series of engineered sequences containing a variable number of target sites for one of five different miRNAs in two different background sequence contexts [51]. As the number of miRNA targets increased from zero to five, REPRESS predictions tracked experimental repression for all five miRNAs in both sequence contexts, and was more accurate than all other models tested (Supp Fig. 8). For hsa-miR-20a-5p and hsa-miR-21-5p, REPRESS correctly predicted that one background sequence had higher repression than the other (*P* = 4.8 × 10^−10^ and 9.7 × 10^−35^, respectively) and also predicted that for hsa-miR-20a-5p, repression strongly increased with multiplicity (*P* = 4.2 × 10^−21^), whereas for hsa-miR-21-5p, it did not (Fig. 3d). Notably, no other method that we tested correctly predicted the observed context-dependent repression or the difference in the effect of multiplicity on repression for the two different miRNAs (Supp Fig. 8), likely because other models are trained on shorter sequences (less than 60 nt) or model binding sites individually.

Given that REPRESS was trained using unperturbed wild type cells and tissues, we next examined whether it could predict miRNA mediated repression in the context of miRNA overexpression and knockout/knockdown experiments [59–62]. For each experiment, we calculated the average miRNA binding predicted by REPRESS or TargetScan, and selected the top 100 targets for each model. We then compared the distribution of log2 fold-changes of the corresponding transcript sets relative to all transcripts analyzed as a baseline. For the representative hsa-miR-122-5p transfection, REPRESS successfully identified transcripts that were significantly more repressed after transfection relative to all transcripts (KS statistic = 0.3, *P* = 1.7 × 10^−8^) and those highly ranked by TargetScan (KS statistic = 0.23, *P* = 0.009). Similarly, for the representative mmu-miR-26a-5p knockout, REPRESS identified transcripts that were significantly less repressed after knockout relative to all transcripts (KS statistic = 0.36, *P* = 3.8 × 10^−12^) and those highly ranked by TargetScan (KS statistic = 0.24, *P* = 0.0061) (Fig. 3e). Notably, for experiments involving transfection of miRNAs with very sparse binding in our miR-eCLIP datasets, transcripts identified by REPRESS were not significantly more repressed after transfection compared to both baseline and TargetScan (Supp Fig. 9).

We also found that REPRESS’s degradome predictions were inversely correlated with RNA stability measurements from an MPRA of ARE-containing 3’ UTRs [63]. Specifically, UTRs with higher ARED-plus cluster numbers [54] and lower stability in the MPRA had corresponding higher REPRESS degradation predictions (Fig. 3f). One notable example of the predictive capabilities of REPRESS’s degradome model was identifying an experimentally validated stabilizing ARE in the mouse *Bcl2* gene whose deletion caused reduction in the measured RNA stability [64] (Supp Fig. 7g). Collectively, these results demonstrate that in learning fundamental aspects of miRNA targeting and mRNA degradation, REPRESS can aid in the discovery of new biology.

### 2.4 REPRESS decodes the impact of genetic variants

Accurate prediction of the impact of variant alleles on gene expression, stability, and disease phenotypes is a foundational goal in the application of deep learning to genomics. Such predictions allow for an improved understanding of the regulatory mechanisms mediating variant effects, and can support fine-mapping efforts to identify causal variants. To evaluate REPRESS’s capacity to determine the effect of sequence mutations on miRNA binding sites, we benchmarked its ability to discriminate a novel curated set of miRNA binding-altering variants against a negative set of background variants from gnomAD [65]. REPRESS successfully identified individual variants known to disrupt (rs1876439052, [66]) (Fig. 4a) and create (rs1063320, [67]) (Fig. 4b) miRNA binding sites, and accurately distinguished the miRNA binding-altering variants from the background variants. REPRESS outperformed existing miRNA binding models with an AUROC of 0.73 (Fig. 4c) while the next best model TargetScan [13] attained an AUROC of 0.59 (Supp Fig. 10b). For “pathogenic” and “likely pathogenic” (P/LP) variants from ClinVar [68, 69], REPRESS predicted higher scores for P/LP variants expected to modulate miRNA targeting relative to other P/LP variants and putative benign variants (Fig. 4d).

**Fig. 4:**
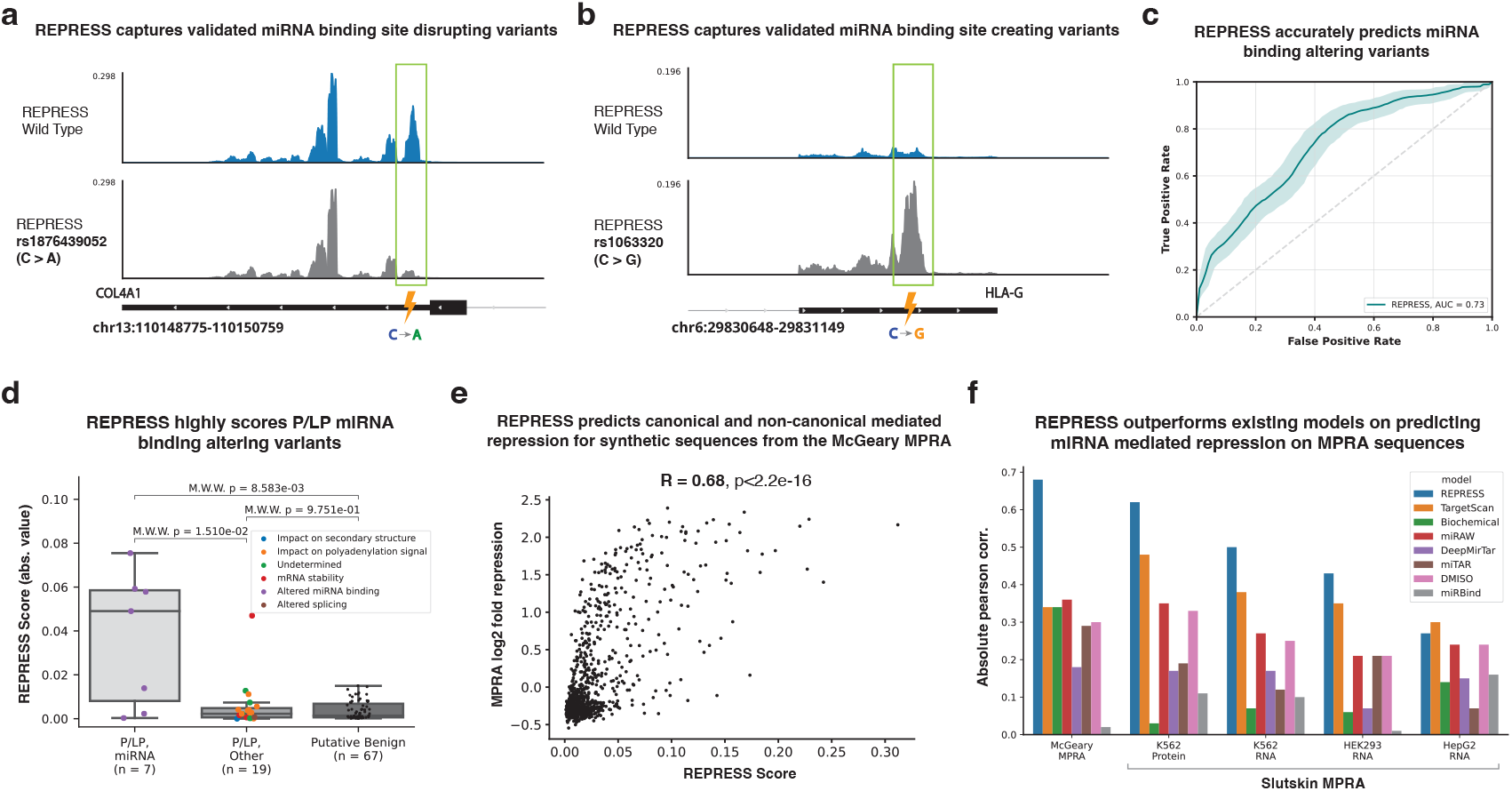
**(a), (b):** REPRESS’s wild type and mutant predictions on verified miRNA binding site altering variants. The (a) variant rs1876439052 disrupts the binding of hsa-miR-29b-3p and the (b) variant rs1063320 creates a hsa-miR-148a-3p binding site. **(c):** REPRESS’s ROC curve for predicting verified miRNA altering variants (n=100, Supp Table 3) from background variants (n=174) with high allele frequencies in gnomAD. Shaded area represents a 95% confidence interval resulting from bootstrapping. The diagonal dotted line represents the performance of a random classifier. **(d):** REPRESS variant scores on miRNA mediated P/LP variants, other P/LP variants and putative benign variants. REPRESS score for a variant is defined as the absolute difference between the mean REPRESS miRNA prediction over the wild type and mutant sequence. Horizontal lines indicate (from bottom to top) the first quartile, second quartile (median), and third quartile. Whiskers extend 1.5 × IQR below Q1 and 1.5 × IQR above Q3, where “IQR” = interquartile range. P values are the result of Mann-Whitney Wilcoxon (“M.W.W”) tests. **(e):** Scatter plot illustrating the relationship between REPRESS scores and log_2_ fold repression from the McGeary MPRA (n=952). Each point represents a synthetic sequence, with REPRESS scores on the x-axis and MPRA log_2_ fold repression on the y-axis. REPRESS accurately predicts miRNA mediated repression for synthetic sequences from the McGeary MPRA with Pearson R=0.68. REPRESS score is defined as the average miRNA binding prediction over the entire MPRA sequence. **(f):** Performance comparison of different miRNA models on predicting miRNA mediated repression from MPRA datasets. REPRESS significantly outperforms other baseline miRNA models for synthetic sequences from the McGeary and Slutskin MPRAs.

We further investigated REPRESS’s ability to identify the underlying causal mechanisms behind disease pathologies and its utility in supporting therapeutic target discovery. The rs712 variant associated with increased risk of non-small cell lung cancer (NSCLC) was experimentally shown to impact expression by disrupting a repressive let-7 miRNA binding site in the 3’ UTR of the oncogenic *KRAS* gene [70–72]. REPRESS successfully made a high-scoring prediction that this variant would decrease miRNA binding (Supp Fig. 10d). We also evaluated REPRESS predictions against a set of 24,574 3’ UTR variants of uncertain significance (VUSs) and found that rs886049197 was also predicted to disrupt let-7 binding, suggesting a similar mechanism through which KRAS upregulation may be conferred (Methods) (Supp Fig. 10d). The other methods that we tested had low precision and scored these variants more weakly than they scored known benign variants, and they also did not exhibit statistically significant enrichment when comparing P/LP to benign variants (Supp Fig. 10g,h).

To evaluate more broadly whether REPRESS’s predictions can generalize to novel sequences and variants, we used MPRA datasets from McGeary et al. [52] and Slutskin et al. [51] to evaluate REPRESS’s ability to decode the impact of variants on miRNA mediated repression. The McGeary MPRA consists of 952 synthetically designed constructs of 120 bp containing either a canonical or non-canonical binding site of let-7a. The non-canonical sites used in the MPRA library included imperfect seed matches with varying degrees of 3’ supplementary binding. REPRESS accurately predicted the fold repression for the synthetic sequences from the MPRA (R=0.68) (Fig. 4e) and outperformed other miRNA binding models by a large margin (Fig. 4f).

The Slutskin MPRA dataset measured changes in RNA and protein levels across multiple cell lines for 12,545 regulatory sequences that modulate miRNA binding by varying target complementarity, target multiplicity, and other sequence features. REPRESS successfully recovered the experimentally measured variant effects as shown by a log-linear relationship between predicted miRNA binding and observed fold-change (Fig. 2f, Supp Fig. 10a), outperforming other competing models (Fig. 4f). It is notable that the sequences evaluated from both MPRAs fall outside of the distribution of wild-type sequences that REPRESS was trained on, but even so, REPRESS was able to accurately generalize its predictions to these datasets in a “zero-shot” manner.

### 2.5 REPRESS facilitates the efficient design of RNA therapeutics

By pinpointing post-transcriptional regulatory elements, REPRESS can be used to inform the design of RNA therapeutics that target these elements to modulate gene expression. For instance, the activity of antisense oligonucleotides (ASOs) can be simulated with REPRESS to identify those likely to increase gene expression upon blocking of a repressive regulatory element (Fig. 5a) (see Methods). Two instances exemplify how REPRESS can predict ASO effects: In one study of the *UTRN* gene, the target region bound by the ASO conferring the highest increase in expression [73] corresponds to a strong miRNA binding site predicted by REPRESS (Fig. 2g). Additionally, when ASO binding was simulated by masking the sequence in this region, REPRESS correctly predicted the loss of the miRNA target. Similarly, in a separate study focused on the *CHD9* gene [74], REPRESS predicted that the lead ASO blocks a strong miRNA binding event without affecting other miRNA events in the same 3’ UTR (Fig. 5b). In both of these examples, the lead ASO was ranked by REPRESS as rank 1 and rank 2 out of all possible 2,037 and 2,606 ASOs, respectively.

**Fig. 5:**
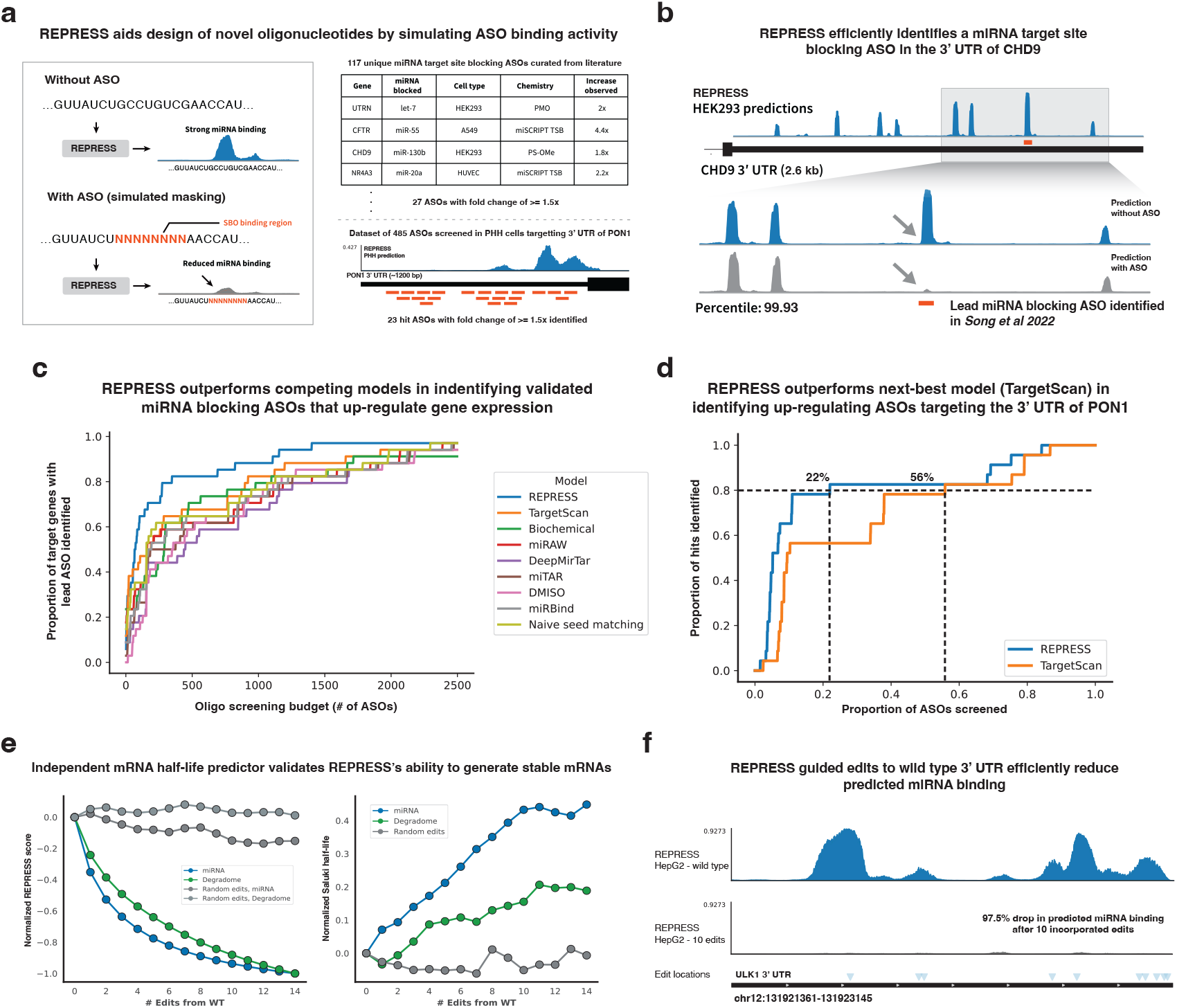
**(a):** RREPRESS aids the design of novel antisense oligonucleotides that sterically block miRNA binding sites. The binding of the ASO is simulated by replacing the wild-type target sequence of the ASO with N’s and the ASO’s activity is ranked by the predicted drop in miRNA binding. To evaluate the performance of REPRESS, we curated 27 miRNA-blocking ASOs that up-regulate their target gene expression by 1.5x and internally screened 485 ASOs targeting the 3’ UTR of *PON1*. **(b):** REPRESS’s miRNA predictions over the 3’ UTR of *CHD9*. The validated upregulating ASO perfectly corresponds to the strongest predicted REPRESS peak and is ranked at the 99.93th percentile (rank 2) out of all possible ASOs (2,606) targeting the 3’ UTR of *CHD9*. **(c):**Fraction of literature-curated ASOs captured by each model as a function of the oligo screening budget (x-axis, representing the number of ASOs screened). For each screening budget, the y-axis indicates the proportion 27 validated hits for which the lead ASO is identified using the different predictors to rank the ASOs. **(d):** REPRESS outperforms the next-best model TargetScan in identifying hit (≥ 1.5x up-regulation of *PON1*) ASOs from the *PON1* dataset. The plot illustrates the fraction of *PON1* ASO hits captured after screening the top x% of ASOs nominated by REPRESS or TargetScan predictions. REPRESS requires ~ 2.5 × fewer ASOs to identify 80% of the hits identified during screening. **(e):** Successive sequence edits nominated by REPRESS have higher stability as predicted by Saluki, which is trained on orthogonal mRNA half-life data. Edits were made within the native 3’ UTRs for 116 genes, with miRNA-nominated edits resulting in a 44.6% average stability increase and degradome-nominated edits yielding a 20.6% average stability increase. **(f):** REPRESS’s miRNA prediction scores drop by 97.5% after incorporating 10 edits nominated by the miRNA predictions for *ULK1*.

To benchmark ASO predictions, we curated a total of 27 ASO hits from 22 studies reported to increase RNA or protein levels of their gene targets by at least 1.5-fold by blocking miRNA binding sites (Fig. 5a, Supp Table 4). For each possible per gene oligo screening budget, we determined the fraction of the 27 ASO hits that would have been found if we had screened every gene using only the budgeted number of ASOs with the highest model predicted scores (Fig. 5c). REPRESS ranked the ASO hits highest among all methods tested, with 18 of the 27 ASO hits scoring in the top 10% of their distributions. Relative to TargetScan, REPRESS required more than four times fewer ASOs on average. None of the existing miRNA models evaluated showed a significant improvement over a naive seed matching approach.

In a prospective study of *PON1*, which is involved in cholesterol metabolism and implicated in diseases including familial hypercholesterolemia [75, 76], we designed 485 ASOs targeting the 3’ UTR and tested them in primary human hepatocytes (PHHs), measuring the corresponding increase in *PON1* expression using an AlphaLISA assay (PerkinElmer) [77, 78] (Fig. 5a and Methods). Of the 485 ASOs, we found 23 hits that increased the expression of *PON1* by more than a therapeutically relevant threshold of 1.5 fold. We used REPRESS to rank all possible ASOs that could target the 3’ UTR of *PON1* and found that REPRESS identified 80% of the hits among its top 22% of predicted ASOs, while for the next-best model TargetScan this hit rate of 80% required 56% of predicted ASOs (~ 2.5 × more) (Fig. 5d). This highlights REPRESS’s ability to prioritize regulatory regions in the 3’ UTR and efficiently identify ASOs that can upregulate expression of their target gene.

Finally, we turned to another therapeutic modality, that of synthetic mRNA and gene therapies. REPRESS’s sequence attribution methods can be used to identify bases that drive miRNA binding or mRNA degradation, enabling the iterative design of sequences predicted to be more stable. As a proof-of-concept study, we selected 116 natural transcripts (Supp Table 5) and progressively introduced single nucleotide changes to their 3’ UTRs using ISM, resulting in a substantial decrease in predicted miRNA binding and degradation scores (Fig. 5e). To validate the impact of these edits on mRNA stability, we scored each sequence variant using the Saluki model [79], a predictor trained using orthogonal half-life data from transcriptional inhibitor and pulse-labeling based protocols to predict mRNA stability from sequence. Half-lives predicted by Saluki increased by an average of 45% after incorporating all 14 edits nominated by REPRESS’s miRNA predictions, and by 21% after incorporating all 14 edits suggested by its degradome predictions (Fig. 5e). An illustrative example of this is shown in Fig. 5f, which shows 10 edits that REPRESS predicted would decrease miRNA binding in the *ULK1* 3’ UTR by 97.5%.

## 3 Discussion

The ability of REPRESS to accurately predict repression mediated by RBPs and miRNAs deepens our understanding of post-transcriptional gene regulation and can profoundly improve RNA therapeutic development. In learning many of the biological principles of post-transcriptional gene repression, REPRESS enables biological discoveries such as atypical modes of miRNA binding, decodes the impacts of competing target sites and genetic variants on miRNA mediated repression, and accelerates both ASO and mRNA sequence optimization important for the design and development of RNA therapeutics.

Developing REPRESS necessitated the generation of a large scale dataset of miRNA binding and mRNA degradation across 29 cell types and tissues in both human and mouse, and the creation of a novel ConvNeXt-inspired neural architecture. REPRESS’s neural architecture has a capacity that far exceeds existing methods (133 million parameters vs tens of thousands), and exhibits a strong scaling law of performance gains with model size. Interestingly, the REPRESS-specific architecture also outperformed alternative implementations utilizing alternative architectures including transformers.

REPRESS can overcome deficiencies in the underlying training data. For example, we demonstrated that REPRESS can correctly identify validated miRNA binding sites not identified in the specific miR-eCLIP datasets used for training. This increased sensitivity is particularly important when modeling repressive phenomena that may push targets of strong interactions below an assay’s detection threshold. The ability of REPRESS to overcome the limitations of assays and ‘denoise the training data’ is advantageous. By jointly modeling miRNA binding and mRNA degradation, REPRESS achieves synergies during training and when making predictions. We found that training on the joint dataset yielded better test performance on the individual miRNA binding and mRNA degradation tasks, compared to training on task-specific data. Notably, we found that this performance increase became larger when we increased the model size. Although REPRESS cannot generalize to cell lines not present in the training dataset, we propose a prototype of a “miRNA-specific” variation of REPRESS and provide preliminary evidence showing that it can predict miRNA targets in unseen cell lines and tissue types based solely on miRNA expression, at least in part alleviating the need to generate additional training data (Supp Fig. 11).

In support of the development of RNA based therapeutics across multiple modalities, REPRESS enables in silico experimentation at scales that are often not experimentally tractable. For instance, REPRESS can nominate genetic variants or therapeutic molecules (e.g. ASOs) that dampen repression and increase overall gene expression for therapeutic targets exhibiting a loss-of-function phenotype. These prioritized variants and/or therapeutic molecules can then be subsequently screened using experimental assays, such as gene editing or screening a prioritized set of ASOs in disease relevant models. REPRESS also enables the design of novel UTR sequences optimized to maximize tissue-specific expression for both mRNA and DNA encoded payloads including gene replacement therapies.

There are several future directions to take this work. Although REPRESS was able to accurately predict miRNA overexpression and knockdown data, the model could be augmented by training on these datasets. Training on additional miR-eCLIP, RBP-eCLIP and structural accessibility data, possibly from diverse cell lines, could increase REPRESS’s ability to better model and predict the effects of miRNAs and RBPs on repression and degradation. Although REPRESS predicts miRNA-mediated repression in a “zero-shot” manner, it could be further fine-tuned on MPRA or mRNA half-life datasets to enhance the accuracy of its repression predictions. A current limitation of REPRESS is that it only learns miRNA binding for the abundantly expressed miRNAs in the specific cell lines used for training. One possible direction is to train a separate miRNA encoder network based on the miRNA sequences and use it to condition the predictions of REPRESS. Such an encoder could generalize and predict the binding of novel miRNAs not present in the training dataset.

REPRESS is a state-of-the-art model for predicting miRNA binding and RNA degradation from sequence, with single-nucleotide resolution, and is a step toward a unified view of repressive processes affecting PTGR. In summary, REPRESS constitutes a major step toward a unified framework to interrogate miRNA binding and mRNA degradation, and provides a platform for the community to better understand the determinants of cell-type-specific RNA regulation and to support therapeutic development.

## 4 Methods

### 4.1 Experimental Methodology

#### 4.1.1 Cell Culture

HepG2 cells were cultured in Dulbecco’s Modified Eagle Medium (DMEM), high glucose, pyruvate [Gibco: 11995065] supplemented with 10% FBS [Gibco: 16140071], 1% penicillin/streptomycin [Gibco: 15140122] and grown at 37°C with 5% CO_2_. K562 were cultured in supplemented Iscove’s Modified Dulbecco’s Medium (IMDM) [Gibco: 12440053] while A549 cells were cultured in DMEM (Gibco: 11965092) supplemented with 10% FBS. Primary Human Hepatocytes (PHHs) were acquired from BioIVT for one male donor and one female donor. They were thawed in Cryopreserved Hepatocyte Recovery Medium (CHRM) [Gibco; CM7000], plated in InVitroGRO CP Hepatocyte Medium (BioIVT: Z99029) supplemented with ROCK inhibitor-Y-27632 [Tocris Small Molecules: 1254/1] for 24 hours, the media was then changed to Cellartis Power Primary HEP Medium (Takara Bioscience: Y20020) and the cells were maintained in culture until day 7.

TET-ON NGN2 transgenic iPSCs were grown on plates coated with Vitronectin XF^™^ [Stemcell: 07180] and grown in mTeSR^™^ Plus media [Stemcell: 5825] supplemented with Rock Inhibitor. For neural induction, TET-ON NGN2 transgenic iPSCs were cultured on matrigel (VWR: 354234) -coated flasks for 3 days in vitro (DIV) in the presence of 2*µ*g/mL doxycycline (Millipore Sigma: D9891) and N2 supplement (Gibco: 17502048). Following neural induction, neuronal cultures were maintained up to 14 DIV in neuronal maintenance medium consisting of NeuroBasal Plus (Gibco: A3582901), B27 Plus (Gibco: A3582801), GlutaMAX^™^ (Gibco: 35050061), MEM Non-Essential Amino Acids (Gibco: 11140050), supplemented with BDNF 10ng/ml, NT-3 10ng/ml and Laminin 1 *µ*g/mL (Millipore Sigma: CC095).

#### 4.1.2 miR-eCLIP

miR-eCLIP data for HEK293T and 8 weeks old C57/BL6J mouse liver tissue were curated from [34]. Huh-7 data was curated from [80]. A549 and K562 data were directly purchased from Eclipsebio (San Diego, USA). miR-eCLIP data for all additional cell lines and tissues were generated by Eclipsebio, following a protocol developed by Van Nostrand et al. (2022) to identify miRNA-Ago2 chimeric pairs. All cell lines were cultured and processed by the authors, whereas whole tissue samples were prepared directly by Eclipsebio. The miR-eCLIP protocol was conducted on duplicate samples of HepG2, iPSC derived neurons, and primary human hepatocytes; and on triplicate samples of Yecuris human liver cells (sourced from Yecuris Corporation), post natal 2-day old C57/BL6 mouse cortex tissue (P2), and 8-weeks old C57BL/6J mouse cortex (sourced from the Jackson Laboratory). Each replicate of HEPG2 consisted of 20 million cells, all other cell samples consisted of 40 million cells. The whole tissue samples were prepared directly by Eclipsebio. For all in-house cell preparations, cells plated in 10-cm dishes were covered with DPBS, no calcium, no magnesium [Gibco: 14190144] and UV-irradiated with 400 mJoules/cm2 using a 254-nM bulb. Samples were then scraped and collected, flash frozen, and packaged for sample transfer. Additional details about the miR-eCLIP protocol are mentioned in miR-eCLIP publication [34].

#### 4.1.3 Degradome-Seq

Triplicate total RNA samples from 3 million cells of A549, K562, HepG2, iPSC-derived neurons, day0 primary human hepatocytes, day7 primary human hepatocytes and Yecuris liver were lysed in RLT buffer [Qiagen: 1015750] supplemented with 1% *β*-mercaptoethanol [Gibco: 21985023] using the RNeasy [Qiagen: 74104] kit. Triplicate total RNA samples of 20 mg of tissue derived from Mouse liver 8-weeks old, Mouse cortex 8-weeks old, Mouse CNS e18 day 4 and Mouse CNS e18 day 11 were homogenized and also extracted using the RNeasy kit. Briefly, the degradome libraries were generated as follows: Poly-T magnetic beads were used to purify polyadenylated RNAs, followed by RNA ligase-mediated adapter ligation to the uncapped 5’ ends of 3’ cleavage products. The libraries were size selected using AMPureXP beads and the cDNA was PCR amplified yielding a library of 200-400 bp fragments. Libraries were sent to LC Sciences (Houston, USA) for 50-bp single-end sequencing on the Illumina Hiseq2500 [81].

#### 4.1.4 Small RNA-Seq

Total RNA extracted by RNeasy [Qiagen: 74104] was collected from 1 million cells in triplicate for A549, K562, HepG2, iPSC-derived neurons, day0 primary human hepatocytes and day7 primary human hepatocytes. Total RNA was also processed from 40 cortices of post-natal 2-day old C57/BL6 mouse tissue (P2) sourced from the Jackson laboratory, and from 5 Yecuris liver samples obtained from Yecuris Corporation. Further processing and sequencing was conducted by The Centre for Applied Genomics (TCAG), Sick Kids Hospital (Toronto, Canada). Briefly, the small-RNA libraries were prepared with the Nextera XT DNA Library Preparation Kit (Illumina) and sequenced by the Illumina HiSeq 2500 platform generating at read lengths of 125 bp and depth of 60M reads per sample.

### 4.2 Data generation and processing

We conducted a thorough curation of AGO2-CLIP datasets from the literature in order to create a comprehensive training dataset. Datasets were discovered by searching the Gene Expression Omnibus (GEO) Database using keywords including “AGO2-CLIP” and “PAR-CLIP”, “HITS-CLIP”, “eCLIP” in conjunction with “miRNA”. We only included studies where AGO2-CLIP was performed in untreated cells, ensuring the data reflected miRNA binding in the cell’s unperturbed, wild-type state. Raw fastq files from each study were downloaded and processed separately. Only studies that yielded over 10,000 non-overlapping AGO2-CLIP peaks across all replicates after trimming, genome alignment, and peak calling were kept. Studies that did not yield sufficient peaks due to low sequencing depths or low genome alignment rates were not included in the final dataset to ensure only high quality data is used for model training. This yielded a total of 18 AGO2-CLIP datasets across human and mouse cell lines curated from the literature.

AGO2-CLIP data curated from the literature was processed in a protocol specific manner. For PAR-CLIP fastp [82] was used to trim the reads with the following arguments : `--adapter sequence XXX -w 16 --trim poly x --cut tail 30`. Once trimmed, the reads were aligned to the genome using the bowtie aligner [83] with the following parameters : `-v 2 -m 1 --best --strata` followed by peak calling using PARalyzer [84].

For HITS-CLIP and eCLIP data, fastp was also used for trimming reads with the same parameters while bowtie2 [85] was used to align the reads to the genome with following arguments : `-D 20 -R 3 -N 0 -L 20 -i S,1,0.50 -k 4 -p 32`. After genome alignment, the reads were de-duplicated using picard and peak calling was done using CLIPper [86].

All miR-eCLIP data was generated and processed by EclipseBio. UMIs were first extracted using umi-tools [87] and the adapters were trimmed using Cutadapt [88]. The chimeric reads were then reverse-mapped to a database of mature miRNA sequences from miRBase [89] using bowtie to get the miRNA identity and then aligned to the genome using the STAR aligner [90]. Peaks were then called on the chimeric reads using CLIPper.

All Degradome-Seq data was processed using the same pipeline. The reads were first trimmed using the fastp trimmer with the following arguments : `--adapter sequence XXX --length required 20`. After trimming, the reads were then aligned to the genome using HISAT2 [91], a splice aware aligner. Peak calling was not performed on the degradome data and the model is trained to predict the Degradome-Seq read coverage as a function of the input RNA sequence.

All miRNA-Seq data was processed using the following pipeline. The small RNA reads were first trimmed using cutadapt with -m 15. Once trimmed, the reads were aligned to the genome using the STAR aligner [90] with the following parameters : --alignEndsType EndToEnd --outFilterMismatchNmax 1 --outFilterMultimapScoreRange 0 --quantMode TranscriptomeSAM GeneCounts --outReadsUnmapped Fastx --outSAMtype BAM SortedByCoordinate –outFilterMultimapNmax 10 --outSAMunmapped Within –outFilterScoreMinOverLread --outFilterMatchNminOverLread 0 --outFilterMatchNmin 16 --alignSJDBoverhangMin 1000 --alignIntronMax 1 --outWigType wiggle --outWigStrand Stranded --outWigNorm RPM. The featureCounts package [92] was then used to count the number of reads mapping to different pre-miRNAs and mature miRNAs in the genome with the following parameters `-M -O -s 1 --minOverlap 3`. After this the counts were transformed to CPM values, the final CPM for a given miRNA was taken as the average across all the replicates.

### 4.3 Model Architecture

For a RNA sequence of length *L*, we first transform it into a one-hot-encoded sequence (A = [1,0,0,0], C = [0,1,0,0], G = [0,0,1,0], T = [0,0,0,1]) to obtain an input matrix of size *L* × 4. N = [0,0,0,0] is used for the part that extends outside the transcript. This matrix is used as the input to the REPRESS model. The output of the model is a matrix of size *L* × *K*, where *K* denotes the total number of cell lines. The *k*-th row of the matrix represents the predicted miRNA binding probability or predicted degradome read coverage of each base pair for the *k*-th cell line.

The REPRESS architecture consists of base layers and output layers, the core components of which are residual blocks composed of dilated convolution layers. All convolution layers have the same number of filters, denoted by *N*, except for the last one in the output layers, which is determined by the number of cell types to predict. Padding is used in convolution layers so that the output sequence has the same length as the input sequence after convolution layers.

The input is first processed with an initial convolution layer and then goes through 16 residual blocks (RB) with gradually increasing convolutional dilation rate (receptive field), the dilation rate for each of the 16 residual blocks are - 1, 1, 1, 1, 4, 4, 4, 4, 10, 10, 10, 10, 25, 25, 25, 25. In each residual block, layer normalization is applied first, the output of which is passed through two separate convolution layers with sigmoid and tanh activation functions. Then the two outputs will go through an element-wise multiplication operation. The convolution and element-wise multiplication is repeated twice. Finally, the input is added to the output as a skip connection. All the residual blocks have the same number of filters, and each set of four residual blocks has filter widths of 11, 11, 21 and 41. The output of the initial convolution layer, and the 4th, 8th, 12th and 16th of the residual blocks are then passed through a convolution layer separately. These five outputs are accumulated, concatenated and passed into the output layers of the model.

The output layers consist of four components based on the data species and data type: human-miRNA, human-degradome, mouse-miRNA, and mouse-degradome. Each component has an identical structure, made up of an initial convolution layer and four residual blocks, except for the final output convolution layer. In each residual block (RB-H) of the output module, a layer normalization layer is applied first, followed by two convolution layers with GELU activation. Then the input to the residual block is added to the output as a skip connection. The four residual blocks have the same number of filters as the ones in the base layers, but have filter widths of 11, 11, 21 and 41, with dilations of 1, 4, 10, and 25, respectively. Before the final output convolution layer, the previous output is cropped at both ends by half the context length (receptive field of the model) so that the final output has the same length as the initial input to the model without the added context sequence. For the last convolution layer, the number of filters is based on the number of cell lines in for the corresponding output, and the activation functions used are sigmoid for miRNA binding prediction and softplus for degradation prediction.

REPRESS’s novel residual block (RB and RB-H) is inspired from changes to the residual block proposed in the ConvNeXt publication [42] which improved the performance on CNNs on ImageNet helping it achieve state-of-the-art performance on par with transformers. The changes involved replacing batch normalization layers with layer normalization and reducing the total number of normalization layers to only 1 per residual block instead of 2. Also, the GELU activation was used in the residual blocks (RB-H) of the output layers instead of the commonly used ReLU activation function. These changes alone boosted the validation performance of REPRESS on predicting miRNA binding and mRNA degradation by 17% and 10% respectively (Supp Fig. 5g).

### 4.4 Model training and evaluation

REPRESS was trained using annotations from ncbi refseq.v109 for the human cell lines and ncbi refseq.m38.v106 for the mouse cell lines from GenomeKit. Only protein-coding transcripts were used to train the model. For each gene, the most principal transcript (according to APPRIS [93]) was used. For genes with no APPRIS annotations, the first transcript in GenomeKit’s gene.transcripts table was used. Each transcript from both the human and mouse transcriptomes were divided into non-overlapping windows of 3000 nucleotides across its mature mRNA sequence. A batch of 10 windows was randomly sampled, and 6.25 kb of transcript context sequence around the sampled window was appended to each side, resulting in a total input sequence length of 15.5 kb used to train the model. REPRESS’s miRNA predictions were trained on all exonic CLIP peaks, and no filtering based on miRNA identity, conservation, or site type was performed.

For a given sequence, the true labels corresponding to the miRNA outputs are a binary matrix of shape (3000, 29) corresponding to the length of the sequence and the total number of human and mouse cell lines/tissues with miRNA binding data and predictions. The binary matrix has a true label of 1 for every location corresponding to a CLIP identified peak and 0 everywhere else. Similarly, the true labels for the degradome outputs are a real valued matrix of shape (3000, 10) corresponding to the degradome-seq read coverages for the given input sequence and the total number of human and mouse cell lines/tissues with Degradome-Seq data/predictions. The degradome-seq labels are processed with squashed transformation (*x*^0.375^) to reduce the skewness.

REPRESS’s miRNA binding outputs are in the range (0, 1) while its degradation outputs are in the range (0, inf). We use binary cross-entropy loss ℒ_1_ for the miRNA binding output and Poisson loss ℒ_2_ for the degradation output. The total loss is the weighted sum of the two:

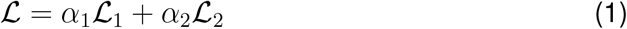

When a sequence from a human transcript is sampled, only gradients from the corresponding human miRNA and degradome output heads are used to update the network, and vice-versa when a sequence from a mouse transcript is sampled. The final model is an ensemble of models trained on the four splits with different sets of chromosomes for training and validation. The four splits use chromosomes (1, 2), (6, 7, 8, 9), (10, 11, 12, 13), and (14, 15, 16, 17, 18) for validation (4-fold cross validation) respectively and the remaining for training. The splits were chosen as described to ensure that ~ 75% of identified miRNA CLIP peaks were used for training and ~ 25% for validation.

For evaluation, we first process predictions and labels of the validation intervals. For each cell line, we have a pre-calculated window length based on median peak width for the CLIP data, and use a fixed length of 50 base pairs for the degradation data. First, we apply average pooling for both predictions and labels using the window length as the pool size. Then, we divide the predictions and labels into multiple chunks with the defined window length. For the miRNA predictions, we assign the value 1 to the averaged label window value ≥ 0.5 and assign 0 otherwise. Then, for each cell line, we calculate area under the curve (AUC) and area under the precision-recall curve (AUPRC) across all intervals for miRNA binding predictions, and Spearman for degradation predictions. We report the mean value of each metric across all cell lines as the final metrics.

The code was implemented in TensorFlow 2.11.0. We initialized the model weights using the Glorot Uniform distribution. For model training, we used the Adam optimizer with default settings for hyperparameters *β*_1_ = 0.9, *β*_2_ = 0.999, *ϵ* = 1 × 10^−8^. We trained with a batch size of 5 on a single NVIDIA RTX A6000 GPU. We tuned the learning rate and the loss weights based on the evaluation metrics on the validation set, and also conducted early stopping in 60 training epochs based on these metrics. The optimal learning rate we found is 0.00005 for the 133M parameter model, with *α*_1_ = 1 and *α*_2_ = 0.1. ReduceLROnPlateau is used for learning rate scheduling, monitoring on validation loss with factor=0.5, patience=5, and cooldown=1.

### 4.5 Sequence attribution interpretation of REPRESS

#### 4.5.1 In-silico mutagenesis

Given a genomic interval starting and ending at positions *p*_start_ and *p*_end_, respectively, and a reference genome sequence *x* of length *L*, the attribution scores *s* are computed as follows: For each position *i* such that *p*_start_ ≤ *i* ≤ *p*_end_, and for each possible nucleotide *u* in the set of nucleotides {*A, C, G, T*}, we generate a variant of *x* by substituting the nucleotide at position *i* with *u*, denoted as *x*_*i,u*_. We then compute the ISM scores *s* for each position *i* and nucleotide *u* using REPRESS, ℳ as given by the equation:

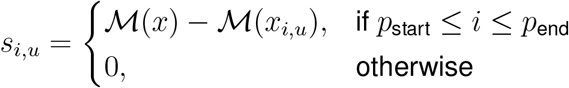

To visualize the effects across the interval, the ISM scores are averaged over all possible nucleotide substitutions, yielding the visualization scores 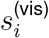:

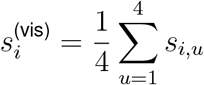

The visualization scores 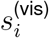 represent the average impact of mutating each position within the interval to any of the four nucleotides.

### 4.5.2 Smoothgrad

Let *X* ∈ ℝ^*L*×*A*^ denote the input tensor, where *L* is the sequence length and *A* represents any of {*A, C, G, T*}). The SmoothGrad saliency map *S* is computed by: 1. Generating Noisy Samples: We create *N* noisy samples of *X* by adding noise drawn from a normal distribution 𝒩 (*µ, σ*^2^):

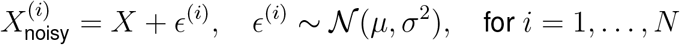

2. Computing the Saliency Map: We calculate the gradients of REPRESS’s output with respect to 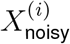, optionally focused on a particular class index *c* when a specific output cell-type head:

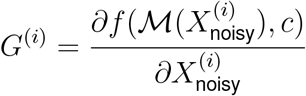

where *f* is the function applied to the saliency map and ℳ denotes the REPRESS score. 3. Averaging the Gradients: The final SmoothGrad saliency map *S* is obtained by averaging the model gradients over all noisy samples:

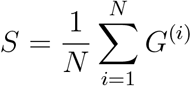

### 4.6 miRNA frequency of complementarity map

We ran LinearCoPartition [38] with the central 100nt of each CLIP peak and the sequence of the corresponding mature miRNA using default settings to get the MFE folded structure and accessibility values for each miRNA-mRNA pair, where the miRNA was between 21-23 nt. After folding, we rejected any structure pair where the miRNA showed intramolecular folding to itself, as this is unlikely to occur while loaded into the RISC complex, and any pair that was predicted to have fewer than four paired positions, as this may represent noise. miRNAs with fewer than 20 instances in each cell-type were also dropped. For each miRNA in each cell type, we counted the average number of times that a position in the miRNA was predicted to be paired and clustered the miRNAs using scikit-learn with n clusters=5, using a k-means++ center initialization. Each cluster was further organized using hierarchical clustering with method=“ward”, to be then plotted using seaborn.clustermap.

### 4.7 Transcriptome wide degradation map

We used hierarchical clustering to construct a heatmap of the Degradome-Seq profiles along all human protein-coding genes. For each gene, the most principal transcript (according to APPRIS [93]) from RefSeq (v109) was used. For genes with no APPRIS annotations, the first transcript in GenomeKit’s gene.transcripts table was used. Degradome-Seq read coverage for each replicate was first normalized to the maximum value anywhere along the transcript, then averaged across replicates, and finally re-normalized to the maximum normalized average value. Replicates with no read coverage for a given transcript, and transcripts with no read coverage in any replicates, were excluded from analysis. To enable comparison of position-specific Degradome-Seq profiles across transcripts of varying lengths, each transcript was split into 205 segments: 15 segments for the 5’ UTR; 100 segments for the CDS, and 90 segments for the 3’ UTR. The number of UTR segments was based on the average length of these regions divided by the average CDS length. These values were then multiplied by 100 (the arbitrarily selected number of CDS segments) and rounded to the nearest five. Transcripts with a 5’ UTR less than 15 bases, CDS less than 100 bases, or 3’ UTR less than 90 bases were excluded from analysis. Degradome-Seq read counts were then averaged across all positions within a segment for clustering using seaborn.clustermap(method = ‘ward’). miRNA target counts were obtained from cell line/tissue-matched miR-eCLIP peaks. Half-life data (for available cell lines) based on 4sU data [94, 95] was obtained from Table S1 from [79].

### 4.8 Degradome and RNA element overlap analysis

To profile Degradome-Seq signal at various potential regulatory sites, we analyzed RBP binding sites for all RBPs with ENCODE eCLIP data [39], miR-eCLIP peaks, and the AU-rich element (ARE) motif UUAUUUAUU. For each of HepG2 and K562 cells, we assigned each peak/site to the most highly expressed overlapping Gencode (v29) transcript based on ENCODE RNA-Seq data. Transcripts were required to have at least one position with at least five Degradome-Seq reads to be included for analysis. Sites in introns, within 100 bases of the transcriptional start site, or within 500 bases of the poly(A) site were excluded from analysis. Degradome-Seq read counts from 100 bases upstream to 100 bases downstream of each site were averaged across replicates. To enable comparison across many transcripts with different baseline levels of expression/degradation, read counts were normalized to the read count at position −100. For each site, a control site was randomly selected from the same region (5’ UTR, CDS, or 3’ UTR) of the same transcript and analyzed in an identical fashion. While RBP binding sites were analyzed at the level of individual RBPs, all miRNAs were analyzed collectively due to data sparsity.

### 4.9 Validation set analysis

REPRESS was benchmarked against TargetScan v8, Biochemical model, miRAW, DeepMirTar, miTAR, miRBind and DMISO for predicting miRNA binding on its validation set. To benchmark TargetScan, for each transcript in the validation set we run TargetScan on the entire transcript and obtain context++ scores for every identified miRNA binding site for the top 50 miRNAs expressed in that cell line. TargetScan scalar values for each binding site were parsed into prediction tracks, where the interval spanned by an identified binding site is assigned the context++ score as the predicted score. Overlapping context++ scores (caused by different miRNAs predicted to bind to the same region) were averaged position wise. We do not provide 3’UTR MSA alignment input to TargetScan due to the difficulty in locating MSA alignments for all ~ 4*k* transcripts in the validation set. Instead, we perform a lookup to precomputed Pct [55] values for each 3’ UTR-binding site, obtained from the TargetScan website. AIRs (affected isoform ratio for a binding site) are also obtained from a lookup table. The precomputed AIRs data was obtained from the “3P-seq tag info” file from the TargetScan website. If no value was found then the value is set to 1. AUC’s are calculated on a window level classification task similar to REPRESS. Similarly, the top 50 expressed miRNAs were used to calculate the transcript wide miRNA binding using the biochemical model.

Unlike REPRESS, miRNA binding models miRAW, DeepMirTar, miTAR, miRBind and DMISO predict miRNA binding as a function of both the target sequence *and* the corresponding miRNA. miRAW models target sequences of size 40 nt; DeepMirTar and miTAR model target sequences of size 53nt; miRBind and DMISO models target sequences of length 50 and 60 respectively. In order to benchmark these models on the REPRESS validation set, we queried the top 30 miRNAs in each cell line against each of the transcripts in the validation set. For each miRNA-transcript pair we slide a window corresponding to the target sequence size of the model across the entire transcript with a stride of 1. Each window-miRNA pair is now scored with the model as this score is used to generate the prediction track across the entire transcript. We further tested querying only those windows that contain a seed region of the corresponding miRNA and assigned a 0 score to any window that did not have a seed region of the corresponding queried miRNA, this consistently yielded better performance for all models.

### 4.10 Predicting experimentally validated miRNA binding sites

For this task, miRNA binding sites from miRTarBase and DianaTarBase were cross-referenced to generate the positive set of cell type specific experimentally validated miRNA binding sites. DianaTarBase contains information about miRNA-target interactions and the corresponding cell type but does not tell you the exact interval of miRNA binding, while miRTarBase contains information about exact binding interval and the corresponding miRNA without cell type information. Hence, binding sites that were present in both these datasets were used to generate a cell type specific set of experimentally validated miRNA binding sites. For the negative set, we took intervals corresponding to 8-mer seed region matches of miRNAs that are not expressed in those corresponding cell lines (≤ 3 CPM). Hence, using both the positive and negative sets we were able to benchmark REPRESS against this task by computing the corresponding AUCs and AUPRCs. For each the baseline models TargetScan, biochemical, miRAW, DeepMirTar, miTAR, miRBind and DMISO were directly queried on each target sequence miRNA pair for both the positive and negative sets. The predicted output scores were used to compute the AUCs and AUPRCs.

### 4.11 Predicting HEAP identified miRNA binding sites

We analyzed miRNA binding sites identified in the HEAP dataset (GSE139344) to predict miRNA-target interactions in exonic regions. Specifically, we focused on peaks detected in the P13 mouse cortex samples (GSE139344 P12 cortex peaks.csv), as this was a cell type that was available in REPRESS (Mouse cortex p2). To ensure relevance, our analysis was restricted to miRNA binding sites located in exonic regions, removing intergenic and non-coding RNA features. Genomic sequences extracted using mm10 (ncbi refseq.m38.v106) for the corresponding intervals. REPRESS predictions were the average Mouse cortex miRNA binding prediction across the entire binding interval identified by HEAP.

We benchmarked against TargetScan, Biochemical, miRAW, DMISO, miRBind DeepMirTar and miTAR incorporating flanking sequence to meet the length requirements of each model’s input format (e.g., 40-60 nt for miRAW, DeepMiRTar, DMISO). Each model was applied to each given sequence and corresponding binding miRNA identified from HEAP. Performance of each model was assessed by stratifying the predictions based on bins of log_2_FC values using predefined thresholds (*<*2, 2-3, 3-4, 4-5, 5-6, 6-7, 7+) to evaluate model performance in identifying repressive vs non-repressive sites across the range of log_2_FC reported. We report ROC and for each model across the range of log_2_FC thresholds (Supp Fig. 6f).

In addition to repressive vs non-repressive peak identification analysis, we also performed an analysis for identifying binding sites identified from HEAP vs non-binding sites. Similar to the validation set analysis, we predicted transcriptome wide binding for Mouse cortex, binned the predictions into windows of 71, and computed the AUPRC in predicting HEAP identified sites from background sites (Supp Fig. 6e).

### 4.12 REPRESS performance comparison heatmap

All the performance values presented in the heatmap are normalized as a percentage of the highest performing model for that particular task. The metrics used for each of the following tasks are as follows - The validation set analysis of REPRESS used the average AUPRC across the 29 human and mouse cell lines; unnormalized values of the model performances can be found in Supp Fig. 6a. The miRTarBase prediction analysis used the average AUROC across the 8 cell lines for which data was cross-referenced from miRTarBase and DianaTarBase; unnormalized AUPRC values for each of the models can be found in Supp Fig. 6d. The HEAP analysis used AUPRC values in from predicting HEAP identified binding sites from non-binding sites; unnormalized values for this analysis can be found in Supp Fig. 6e. Both the miRAW and DeepMirTar test sets report AUPRC values for all the miRNA models; unnormalized values can be found in Supp Fig. 6b,c. The McGeary MPRA dataset uses the Pearson correlation for predicted scores and repression for the 952 sequences from the let-7a transfection MPRA. The Slutskin MPRA uses the average Pearson correlation for predicted scores and repression across 4 values - K562 protein readout, K562 RNA readout, HEK 293 RNA readout, HepG2 RNA readout; unnormalized values for all models can be found in Fig. 4f. The variant effect prediction task uses the AUPRC values of predicting validated miRNA binding altering variants from background variants; the unnormalized values for all models are shown in Supp Fig. 10b.

### 4.13 REPRESS performance comparison heatmap

For each of the top 10 miRNAs expressed in each of the cell lines in our dataset, we took the top 500 predicted miRNA binding sites with a seed region match and computed the ISM sequence attribution scores for each of the binding sites. The sequence attribution scores for each miRNA’s corresponding binding sites were averaged across all of its top 500 sites and the average score at each position was used to determine the impact of complementarity at that particular position of the miRNA.

### 4.14 Atypical miRNA binding analysis

REPRESS sequence attributions were performed on the highest scoring 100 miR-eCLIP targets according to REPRESS for each of the top 20 most highly expressed miRNAs in each cell line / tissue, as described in Methods. Sequence attribution scores were aligned, averaged across all targets, scaled to the largest value, and presented in terms of the corresponding mature miRNA sequence (i.e. the reverse complement of a perfectly paired target). Average attributions *<* 0 were plotted as 0 for clarity. For each miRNA with at least 250 unique seed-containing miR-eCLIP targets across all human cell lines profiled with miR-eCLIP, the proportion of canonical targets (including offset 6mer, 6mer, 7mer-A1, 7mer-m8, and 8mer site types) with an offset 6mer site type was determined. A two proportion Z-test was used to compare the proportion of offset 6mer sites between hsa-let-7a-5p and hsa-miR-148a-3p. Conservation was calculated as the proportion of target bases opposite miRNA positions 2-7 with a Phylo-P 100-way score *>* 3. For hsa-let-7a-5p and hsa-miR-148a-3p, statistically significant groupings of site types were identified using two proportion Z tests and Benjamini-Hochberg (BH) adjustment for multiple hypothesis testing.

### 4.15 Comparison of 6mer targets and 8mer targets of miRNAs

All human miRNAs with at least 50 unique 6mer targets and 50 unique 8mer targets in our miR-eCLIP dataset were eligible for this analysis. For each target, we determined the number of conserved bases in the non-seed region–corresponding to bases −13 through −2, where base 0 is the most 5’ base of the 6mer seed match. Bases with PhyloP 100-way values *>* 3 were considered to be conserved. For each miRNA, the odds ratio for the odds of a base being conserved versus non-conserved in 6mer targets versus 8mer targets (and the associated BH-adjusted P values and confidence intervals) were calculated using the statsmodels Python package. Sequence attributions were generated separately for 6mer and 8mer targets of all highly expressed (top 20 in any cell line) miRNAs with absolute log2 odds ratios *>* 0.5 (n = 6), for comparison of sequence attributions and conservation rates in the non-seed regions of 6mer and 8mer targets.

### 4.16 miRNA binding multiplicity analysis

The Slutskin et al. paper included 557 MPRA constructs specifically to analyze the effect of miRNA binding site multiplicity for 5 different miRNAs - hsa-miR-320a-3p, hsa-miR-21-5p, hsa-miR-92a-3p, hsa-miR-19b-3p and hsa-miR-20a-5p. Each MPRA construct was designed to have between zero and five miRNA binding sites for each the corresponding miRNAs The locations of the five possible binding sites were fixed across all sequences and in each construct, each of these locations each site contained either a miRNA binding site or one of two control context sequences: ATGATTATGTCCGGTTATGTA or AAGTTTTCACTCCAGCTAACA.

For each of the five miRNAs, all sequences with the same number of miRNA target sites and the same context sequence were used to calculate average values for: i) observed experimental repression, and miRNA binding predictions using ii) REPRESS, iii) TargetScan, iv) Biochemical model, iv) miRAW, v) DeepMirTar, vi) miTAR, vii) DMISO, and viii) miRBind. Biochemical model fold-change predictions were multiplied by −1 to represent repression. All model predictions excluding REPRESS were scaled to the maximum value predicted by each model across the entire dataset. P values for the impact of context and multiplicity on REPRESS predictions were calculated for all sequences with *<* 5 miRNA targets using formula.api.ols() with “score ~ context + multiplicity” and stats.anova lm() with “type=2” from the statsmodels Python package.

### 4.17 miRNA transfection and knockout datasets

We systematically processed datasets from GSE127211, GSE197363, GSE97060 and GSE123311 to extract log2FC information pertaining to each miRNA-mRNA pair. For each transcript, we first calculated REPRESS miRNA binding prediction across the entire 3’ UTR, then using the sum of REPRESS predictions across all identified miRNA seed sequences we combined these predictions into a single composite score corresponding to the transcript. With this composite score, we ranked the transcripts and selected the top 100 (N=100) with the highest predictive scores to construct the eCDFs.

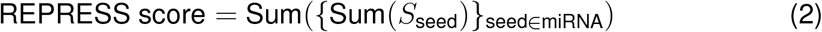

where *S*_seed_ represents the set of scores for each identified seed sequence of the miRNA being tested. To obtain the TargetScan TargetScan was run on every transcript-miRNA pair. The repression score for the transcript is the sum of TargetScan context++ scores for all identified binding sites in the 3’UTR. Again we ranked the transcripts and selected the top 100 to construct the eCDFs.

### 4.18 Positional preference of miRNA targeting analysis

For this analysis, we first intersected miR-eCLIP peaks with human 3’ UTRs from principal transcripts (according to APPRIS [93]) of protein coding genes in v109 of NCBI refseq. For genes with no APPRIS annotations, the first transcript in GenomeKit’s gene.transcripts table was used. Peaks only from miRNAs annotated with conservation *>* −1 in miR Family Info.txt and with *>* 500 peaks in the cell line of interest were considered for this analysis. Peaks passing these filters were then assigned a bin between 1 and 100 based on the relative position of their seed match (or middle of the peak for peaks with no seed matches) along their 3’ UTR, where the first base downstream of the stop codon corresponds to bin 1 and the last base before the poly(A) tail corresponds to bin 100. For each miRNA, the number of peaks in each bin was divided by the total number of peaks. Min-max normalization was applied to these values to enable comparison across miRNAs. These miRNA-specific values were clustered and plotted with seaborn.clustermap().

### 4.19 Predicting validated miRNA binding altering variants

We curated the literature to create a dataset of 100 experimentally validated miRNA binding sites altering (disrupting or creating) variants. This dataset was used as the positive set in order to benchmark REPRESS’s performance on predicting miRNA binding altering variants. For the negative set, common variants from gnomAD with an allele frequency of ≥ 1% in the same 3’ UTRs as the variants from the positive set were taken. The assumption here being, common variants are likely benign and do not disrupt any major underlying cellular processes. This dataset was then used to benchmark the performance of REPRESS on predicting miRNA altering variants.

In order to compute the score of a variant, we computed the mean prediction across the miRNA binding interval for the wild type and mutant prediction and took the absolute difference between these two values. We also averaged this difference across all of REPRESS’s human cell line predictions to generate a scalar value corresponding to the impact score of that variant. For all baseline models, we considered a union of the top 50 miRNAs expressed in all the cell lines and calculated the mean miRNA binding prediction across for the top 50 miRNAs for the wild type and mutant sequences. The absolute difference between the two mean predictions were used to score the variants.

### 4.20 Application of REPRESS to pathogenic and likely pathogenic (P/LP) Clinvar variants that are curated to impact miRNA binding

The dataset consisting of pathogenic or likely pathogenic (P/LP) 3’ UTR variants and putative benign variants was obtained from [96]. For each variant, one prediction was made for the wildtype sequence and one was made for the mutant sequence as follows: The region in the genome affected by the variant was expanded 20bp on either side. This interval was given to REPRESS for inference and the human cell line track predictions corresponding to miRNA binding across the entire interval were extracted. For each cell line head, the average across the prediction interval was taken to get a single value per cell line. The mean of the resulting cell line scalar values was taken to get a single mean scalar value for all the human cell lines. The variant effect was computed as the difference between the wildtype and mutant prediction (mut - wt). The dataset was stratified into P/LP variants that were curated to impact miRNA binding (n=7), P/LP variants curated to operate through another mechanism or have an unknown mechanism (n=19), and putative benign variants (n = 67). We plot the absolute value of the variant effect and perform a Mann-Whitney Wilcoxon 2-sided test between the variant effect scores for each group.

### 4.21 Applying REPRESS to VUS 3’ UTR variants in Clinvar

Variants from ClinVar (last accessed Apr. 30, 2023) that were classified as Variants of Uncertain Significance (VUS) and in 3’ UTR of the transcripts that they were reported in were extracted. These variants were scored using REPRESS in the same manner as the section: “Applying REPRESS to pathogenic and likely pathogenic (P/LP) Clinvar variants that are curated to impact miRNA binding”. From the dataset of variants that impact miRNA binding and common variants (Fig 4b), we split the miRNA binding altering variants into those that create miRNA binding sites and those that disrupt miRNA binding sites. In separate classification tasks for these two sets of variants, we derived thresholds for the 5% FPR and annotated this on a distribution histogram of VUS variant scores from REPRESS.

### 4.22 Applying REPRESS to VUS 3’ UTR variants in Clinvar

The Slutskin MPRA dataset included experimental measurements of repression in four cell lines: K562, HEK293, HepG2, and MCF7, covering 12,545 synthetically designed 176 nt sequences. Since REPRESS was trained on CLIP data from each of these specific cell lines, predictions were made directly for the corresponding cell lines and correlated with the measured fold change. REPRESS was queried using the 176 nt synthetic constructs, and the average prediction across the length of the construct was used to calculate the REPRESS score for each sequence.

The McGeary MPRA library consisted of 952 distinct 120 nt 3’ UTR fragments designed to include both canonical and non-canonical targets of let-7a with varying levels of 3’ complementarity. Let-7a was transfected into DMEM cells and used to measure the corresponding repression on each of the MPRA constructs. Since let-7a was transfected into the cells, we picked the cell line corresponding to the highest endogenous expression of let-7a to query from REPRESS, which was Yecuris liver. REPRESS predictions were averaged across the entire 120 nt sequence to generate the REPRESS score which were then correlated with measured values of repression from the MPRA.

Unlike REPRESS, which makes a direct sequence-to-binding prediction and does not require specification of any miRNA(s), existing models require an individual miRNA to be specified for each query. Therefore, we queried existing models using the top 100 expressed miRNAs in each of the cell lines from the Slutskin MPRA, averaging predictions across miRNAs to generate scalar values representing predicted repression for each MPRA construct. For the McGeary MPRA, each model was queried with let-7a specifically. The MPRA constructs were divided into overlapping windows, each with a stride of 1 and a length equal to the maximum input length tolerated by each model.

Predictions were then averaged across all windows and miRNAs to produce a final scalar value representing predicted repression for each model. For miRAW, DeepMirTar, miTAR, miRBind and DMISO we further tested querying only those windows that contain a seed region of the corresponding miRNA and and assigned a 0 score to any window that did not have a seed region of the corresponding queried miRNA, this consistently yielded better performance for the Slutskin MPRA for all models.

### 4.23 Curated miRNA blocking ASOs Dataset and ASO prediction

117 unique ASOs targeting miRNA binding sites were curated from the literature. Information like assay used to measure fold change, ASO chemistry, target gene, in vitro / in vivo and corresponding cell type used to measure ASO efficacy was also tracked. 27 out of the 117 curated ASOs increased the expression of their target gene by at least 1.5X. In order to measure the efficacy of REPRESS in identifying efficacious miRNA blocking ASOs we measured the reduction in number of ASOs we would have to screen in order to identify the hits when screening ASOs in decreasing order of their predicted score. We simulate the effect of an ASO with REPRESS by masking out the interval corresponding to regions complementary to the ASO and zeroing out the one-hot encodings of that interval. We then take the difference between the mean prediction with and without masking to measure the impact of that ASO in that corresponding cell line. If the corresponding cell line the ASO was tested in was not available we took the average difference across all cell lines.

For all baseline models, a similar approach as the validation set analysis was used to generate track-like predictions from the model using the top 30 expressed miRNAs in that cell line. The average prediction over the ASO interval is taken as the score for that ASO in that particular cell line. For miRAW, DeepMirTar, miTAR, miRBind and DMISO we further tested querying only those windows that contain a seed region of the corresponding miRNA and and assigned a 0 score to any window that did not have a seed region of the corresponding queried miRNA, this consistently yielded better performance for all the models.

### 4.24 PON1 screening SBO Dataset

We screened 485 PS-MOE ASOs designed to modulate miRNA interactions within the 3’ UTR of the *PON1* gene (RefSeq transcript NM 000446). These were screened using an AlphaLISA assay [PerkinElmer: AL389C] to ascertain *PON1* protein expression within primary human hepatocyte (PHH) cells. 18,000 PHH cells per well were reverse transfected with ASO using RNAiMAX [Thermofisher: 13778-150] in a 384-well format in InVitroGRO CP Hepatocyte Medium (BioIVT: Z99029) supplemented with ROCK inhibitor-Y-27632 [Tocris Small Molecules: 1254/1] for 24 hours, the media was then changed to Cellartis Power Primary HEP Medium (Takara Bioscience: Y20020) until day 7. The cells were then lysed in Alphalisa lysis buffer [PerkinElmer: AL003F] supplemented with 1X HALT protease buffer [Thermofisher: 78439] and samples were analyzed using the *PON1* (human) AlphaLISA Detection Kit [PerkinElmer: AL389C].

To compute fold changes, ASO-treated wells were normalized to the mean of three wells treated with a non-targeting ASO control, after both sets of measurements were fit to a standard curve. ASO fold changes were then reported as the mean fold change across three replicates. ASO hits were defined as ASOs that achieved a mean fold change of at least 1.5x. At all five lengths ranging from 18 nt to 22 nt, we scored using REPRESS and TargetScan the 1268 possible intervals falling within the *PON1* 3’ UTR of the given length, yielding 6,340 scores for each model. REPRESS scores were computed as the mean difference between wildtype and N-masked sequence intervals, when used to predict drop in miRNA binding in PHH for this transcript. TargetScan scores were computed as the mean score across the 50 most highly expressed miRNAs in PHH, with the score defined as the difference between the wildtype and N-masked sequence. For the 23 ASO hits exceeding the 1.5x expression threshold, we then determined where their corresponding interval scores fell within the distribution of all 6,340 interval scores for each model.

### 4.25 REPRESS for synthetic mRNA design

We performed ISM on 116 (Supp Table 5) protein-coding genes with high endogenous miRNA binding predictions whose 3’ UTRs were between 1500 and 2000 nt for the human cell lines. The squashing transformation in the degradome tracks was inverted using *x*^1.0*/*0.375^, where x is the model score. Cell lines corresponding to miRNA and degradome were averaged independently, to produce the average ISM result across all cell lines and tissues for degradome and miRNA, respectively. Using the ISM matrix, we then selected the position with the minimum most value (i.e. the mutation that would most reduce the model prediction); importantly, wild-type positions were masked so as to ensure a mutational event at each iteration and prevent getting stuck into local minima. At each iteration, for 14 cycles, the variant resulting in the minimum-most score of the model was selected and used as the input for the next cycle. We also utilized Saluki as our oracle for assessing sequence stability and half-life; for each iteration and mutation, we scored the sequence using Saluki with its default parameters and model weights from the original publication to evaluate improvements in stability. To analyze and visualize the changes in both Saluki and REPRESS scores relative to the wild-type (wt), unedited sequence, we first applied minmax scaling to the scores of each gene across all edits to standardize them. We then subtracted the score of the wt sequence from each edited sequence’s score.

## 6 Supplementary Figures

**Supp Fig. 1:**
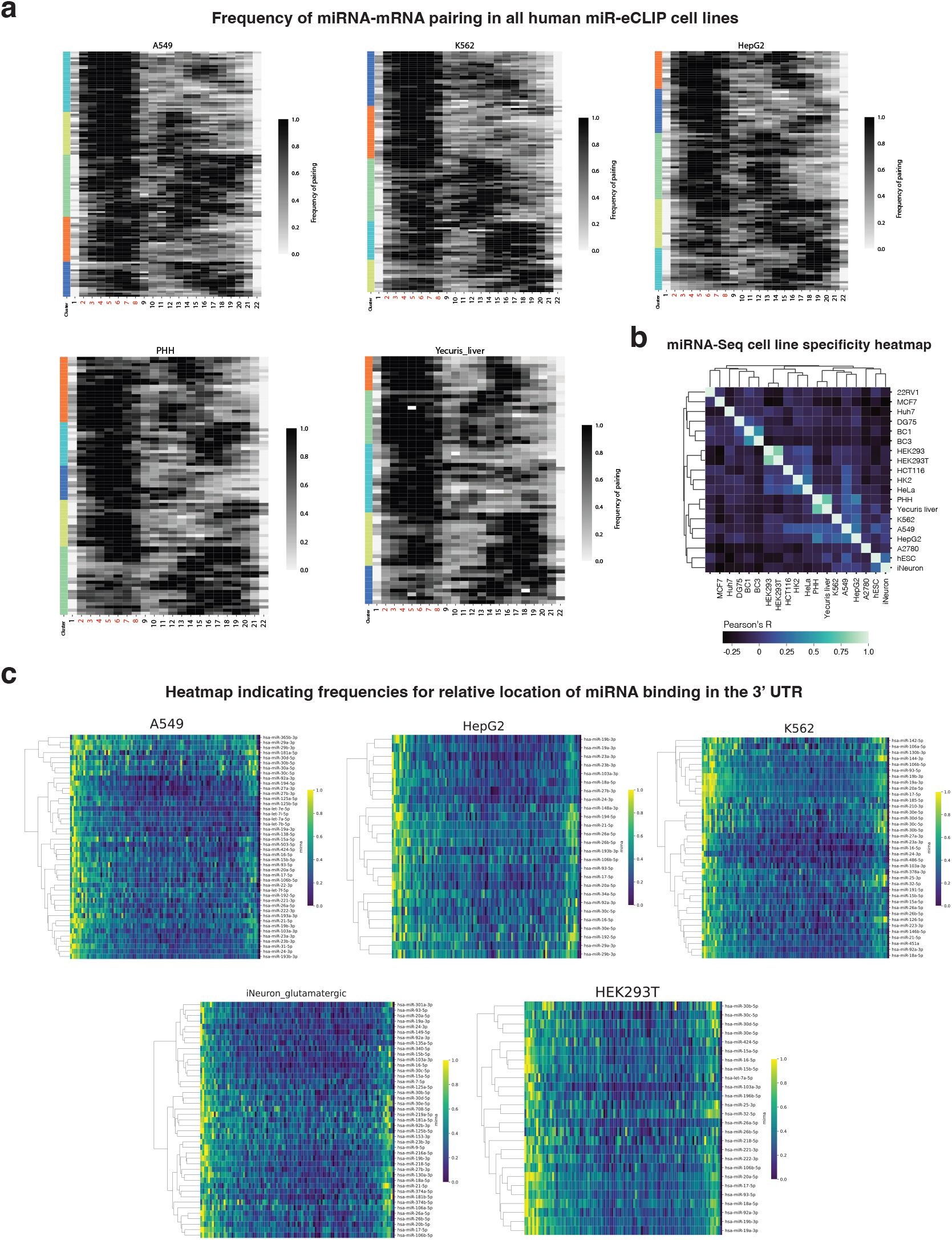
**(a):** LinearCoPartition analysis of the rate of miRNA-target base pairing along the entire mature miRNA for all the other human miR-eCLIP cell lines. Each row represents the average number of times the base was predicted to be paired to an mRNA across all targets of a single miRNA. K-means clustering was performed with Kmeans++ initialization. **(b):** Pairwise Pearson correlations of miRNA expression levels (top 50 miRNAs) across cell lines and tissues. **(c):** Heatmap indicating the frequency of miRNA binding across their relative 3’ UTR target locations for each miRNA across all the human miR-eCLIP cell lines. Preferences in binding near either end of the 3’ UTR and depletion in the middle = consistently seen across miRNAs and cell types.

**Supp Fig. 2:**
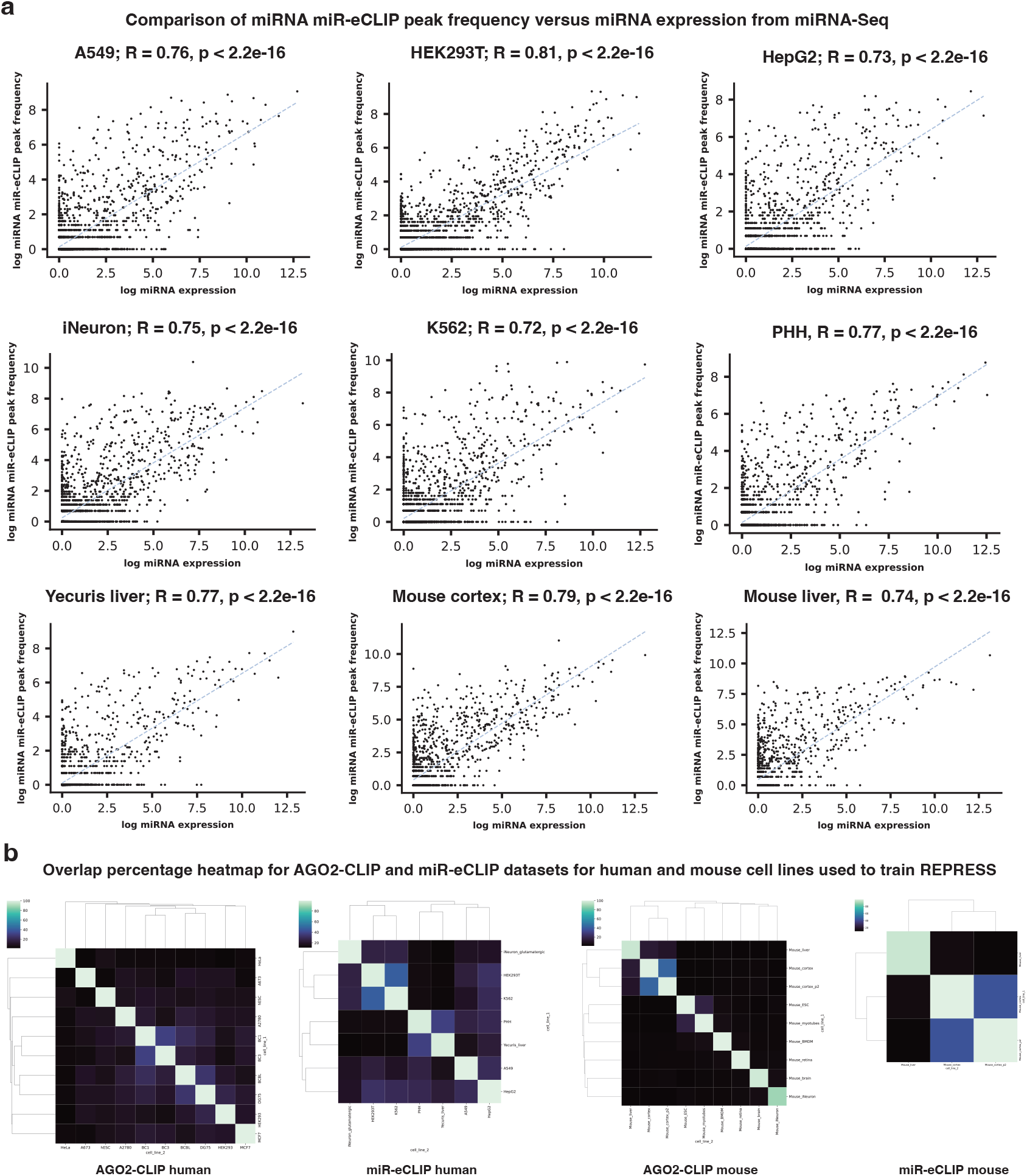
**(a):** Correlation between miRNA expression (CPM) from miRNA-Seq and number of miRNA binding events identified in the miR-eCLIP data. Overall a high correlation is seen across all cell lines (R*>*0.72) for miRNA expression and number of binding events. **(b):** Overlap analysis for the different CLIP identified peaks for the human and mouse AGO2-CLIP and miR-eCLIP cell lines. The average overlap between peaks from across all pairs of miR-eCLIP cell lines was found to be 30%, while the average overlap across all AGO2-CLIP datasets was found to be 14%, with the highest overlap being 72% between two related B-cell cell lines DG75 and BCBL.

**Supp Fig. 3:**
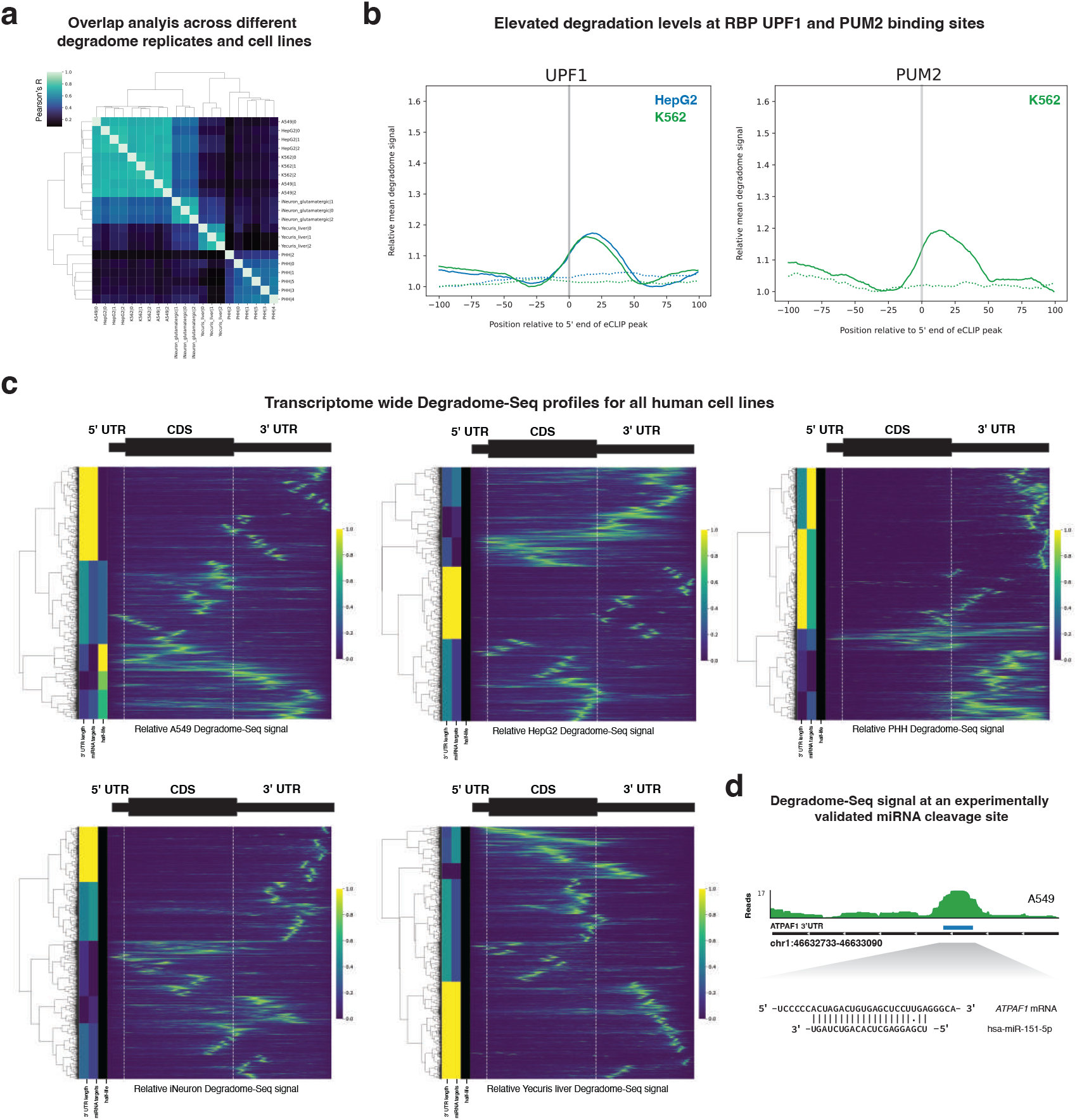
**(a):** Heatmap showing the correlation between the different degradome read coverages across human all cell lines and all the replicates for each corresponding cell line. **(b):** Relative degradome signal along protein-coding transcripts for all other human cell lines. Each row represents a transcript. Each column corresponds to a segment of the 5’ UTR, CDS, or 3’ UTR scaled so that all transcripts can be visualized together. **(c):** RMetaplots of average relative degradome signal at UPF1 and PUM2 RBP eCLIP peak loci. Dashed lines indicate degradome signal around control sites selected randomly from the same 3’ UTRs. Grey vertical bars represent the estimated location of the RBP binding site. To enable inter-cell line and target-control comparisons, degradome values for each dataset are scaled such that the minimum degradome value in the +/- 100 base window is 1.0. **(d):** Degradome-Seq read coverage at a hsa-miR-151-5p target in the ATPAF1 mRNA where transcript cleavage has been previously validated.

**Supp Fig. 4:**
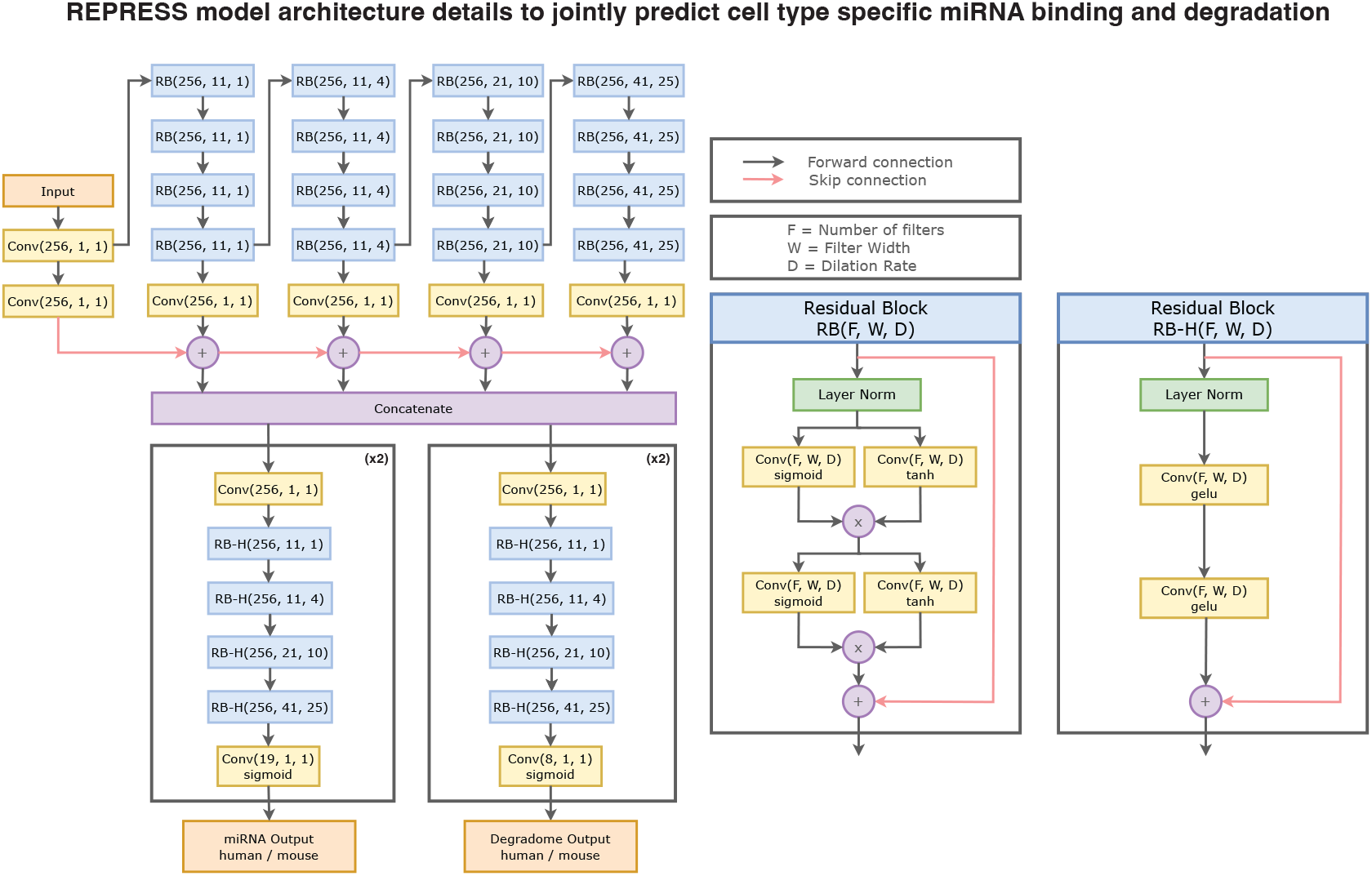
REPRESS model architecture including residual blocks and dilated convolution layers to jointly predict miRNA binding and degradation for a given input mRNA sequence. Each Residual Block / Convolution layer has the following parameters; F: Number of the convolution filters used, W: Filter width of each convolution filter and D: dilation rate of the convolution filters. This block diagram contains all the details to reproduce the REPRESS architecture.

**Supp Fig. 5:**
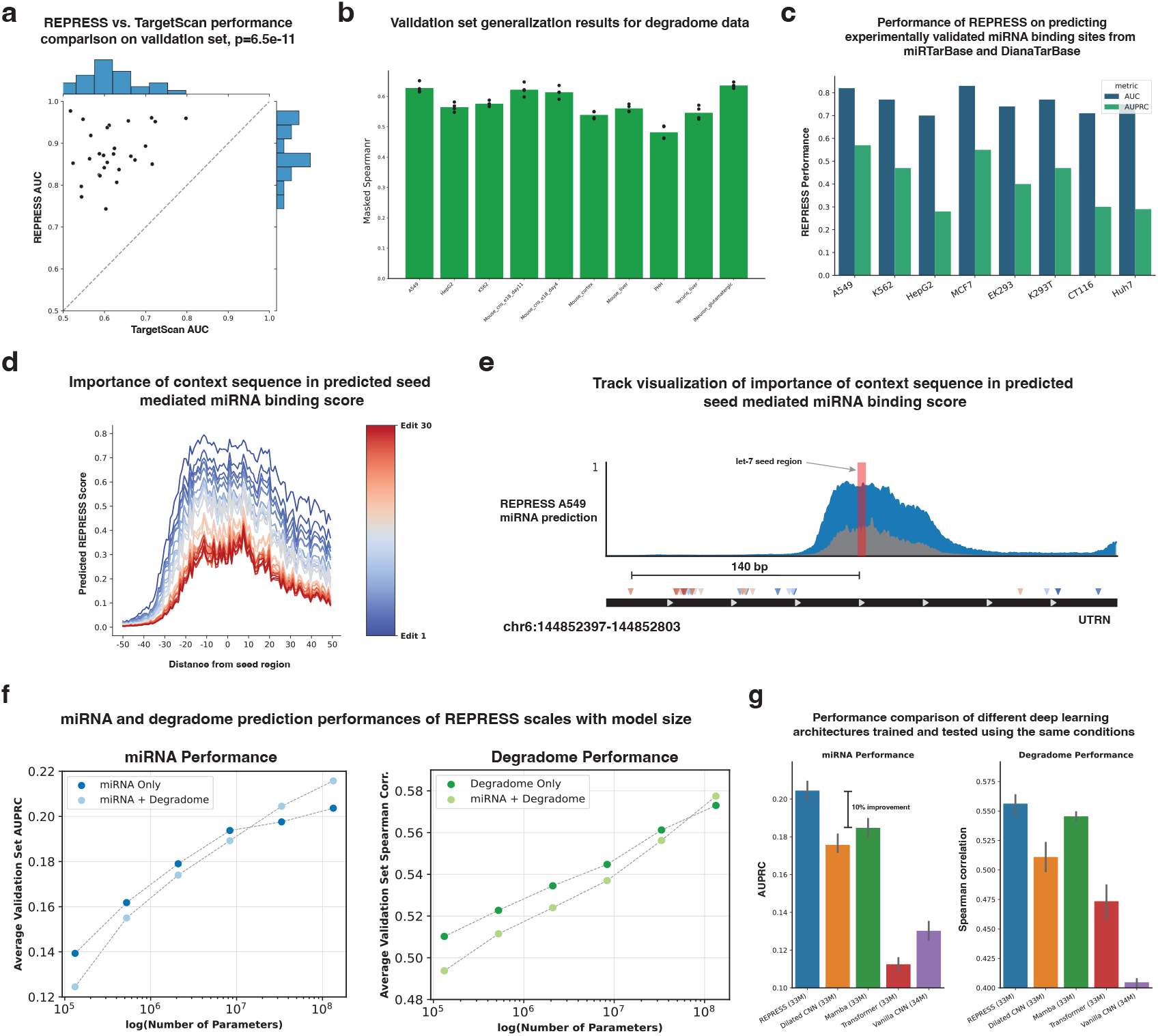
**(a):** Performance comparison of REPRESS and TargetScan in identifying miRNA binding sites in each of the cell lines from the validation set of our miRNA CLIP dataset. REPRESS outperforms TargetScan across all cell lines (shown as dots in scatter plot) with higher AUROCs. **(b):** Validation set performance of REPRESS on the Degradome-seq datasets. Each point corresponds to the performance on the validation set of each fold done for the 4-fold cross validation. **(c):** Performance of REPRESS in predicting miRNA binding sites validated in miRTarBase and DianaTarBase. This shows that the model is able to generalize beyond the CLIP dataset it was trained on. **(d):** Plot illustrating the importance of context sequence in predicted miRNA binding score for let-7 miRNA binding site in the 3’ UTR of *UTRN*. Each line indicates the predicted binding track after each incorporated non-seed edit. The edits were found using ISM within the context region of the let-7c binding site. After 30 non-seed edits the predicted miRNA binding score drops by *>*50%. **(e):** Track plot illustrating the REPRESS predicted miRNA binding track for the wild type sequence and the sequence after 30 non-seed edits, showing the location of the corresponding edits. The furthest edit is 140 bp away from the seed region of the miRNA binding site. **(f):** The performance of both miRNA binding and degradome tasks of REPRESS increase with the model size (number of parameters). This is consistent with scaling laws observed in deep learning architectures for sequence modeling [45, 46]. Additionally, jointly training on miRNA and degradome tasks increases generalization performance for both but only for the larger models. **(g):** Performance comparison of other popular ML architectures when trained on the REPRESS dataset. All the architectures were allowed to have ~ 34M parameters. The ConvNeXt inspired REPRESS architecture outperforms all existing popular ML architectures. The miRNA binding performance of REPRESS is 10% better than the second best Mamba State-Space architecture and ~ 17% better than a dilated CNN with similar number of parameters and the same 12.5 kb context window.

**Supp Fig. 6:**
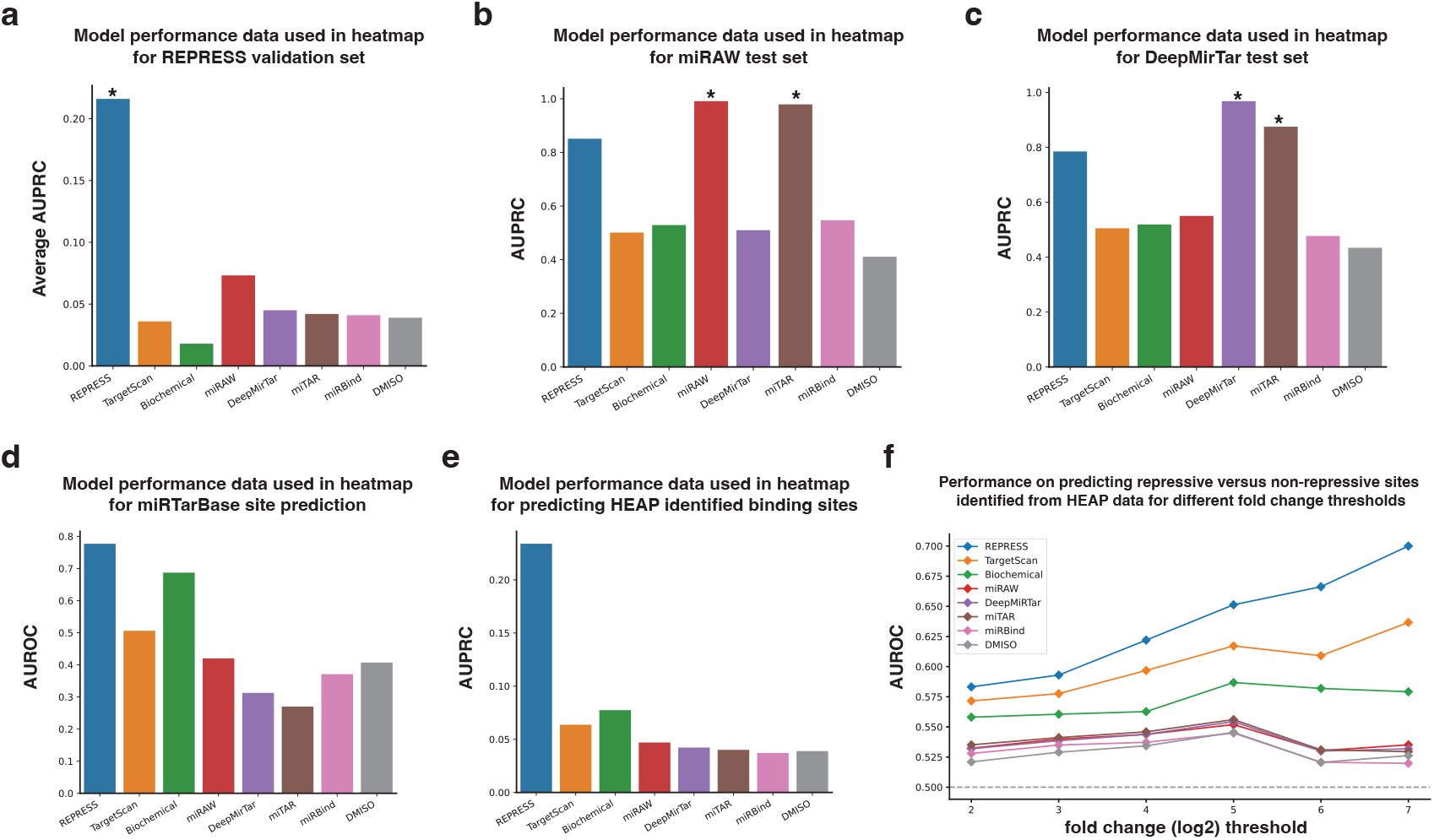
Model performance values used in the heatmap for each of the individual datasets / tasks used for validation (a-e). The * indicates that the dataset was used as the validation / test set for those corresponding models, hence, is from the same distribution as the data used to train that model. **(a):** Performance comparison on the REPRESS validation set using the average AUPRC across the 29 different cell lines. REPRESS significantly outperforms all baseline models. **(b):** Performance comparison on the test set from the miRAW model. The performance of REPRESS is close to miRAW and miTAR which were trained on data from an identical distribution. **(c):** Performance comparison on the test set from the DeepMirTar model. The performance of REPRESS is close to DeepMirTar and miTAR which were trained on data from an identical distribution. miRAW fails to generalize on the DeepMirTar test set and vice-versa. **(d):** Average AUPRC across 8 cell lines for predicting experimentally validated miRNA binding sites from miRTarBase and DianaTarBase. **(e):** REPRESS shows a significant performance improvement over all baseline models in predicting miRNA binding sites from non-binding sites identified from the HEAP assay. HEAP is an orthogonal assay different from the CLIP data used to train the model which REPRESS accurately generalizes to. **(f):** HEAP identifies miRNA-mRNA interactions and quantifies their repressive effect via log2 fold change in target expression. REPRESS outperforms existing models in identifying repressive HEAP identified sites from non-repressive HEAP identified sites for different log2 fold change thresholds.

**Supp Fig. 7:**
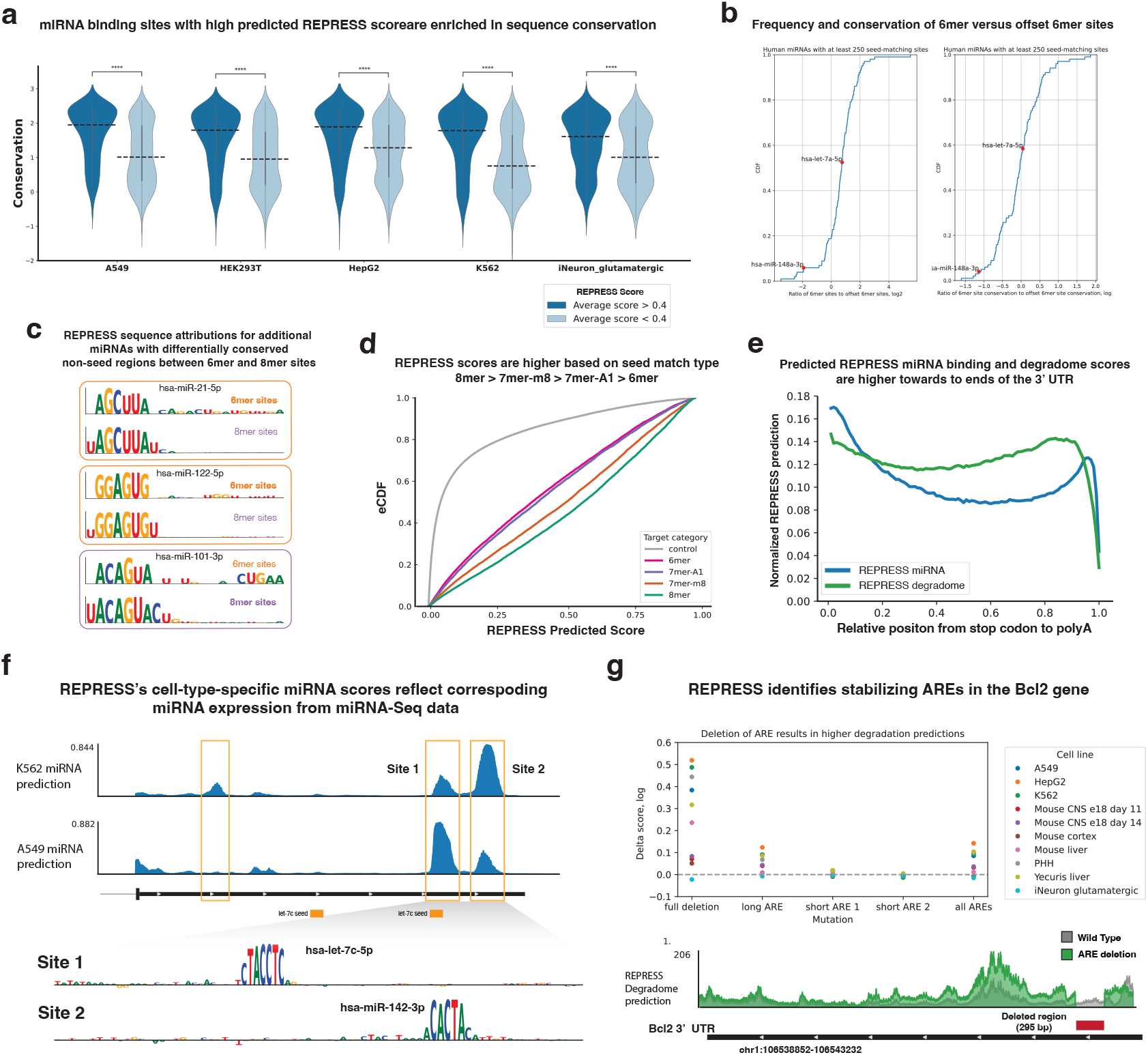
**(a):** miRNA binding sites with higher predicted REPRESS scores have a statistically significant enrichment in PhyloP conservation over predicted binding sites with lower scores across all the cell lines. **(b):** CDF plots for the prevalence of 6mer targets relative to offset 6mer targets (left), and the rate of conservation across vertebrates for 6mer targets relative to offset 6mer targets (right) for human miRNAs. Values for miRNAs hsa-let-7a-5p and hsa-miR-148a-3p profiled in Fig. 3a are highlighted in red. **(c):** REPRESS sequence attribution plots for highly expressed miRNAs with substantial differential conservation of non-seed regions between 6mer and 8mer targets not shown in Fig 3b. Base heights show the relative impact on miRNA binding that REPRESS assigned to each position (represented in terms of the corresponding miRNA position/base). **(d):** REPRESS scores for different seed match types follow the order of repressiveness established in the literature, 8mer *>* 7mer-m8 *>* 7mer-A1 *>* 6mer. **(e):** REPRESS’s miRNA and degradome predictions are higher towards the ends of the 3’ UTR closer to the stop codon and the poly(A) tail. Binding towards the termini of the 3’ UTR has been experimentally shown to be more repressive [57]. **(f):** Visualization of REPRESS’s miRNA predictions on A549 and K562 cell lines for the 3’ UTR of the gene *UTRN*. REPRESS scores site 2 higher in K562 and site 1 higher in A549. The predicted scores correspond to the miRNA expression of two miRNAs with hsa-let-7c-5p being higher expressed in A549 and hsa-miR-142-3p being higher expressed in K562. **(g):** REPRESS’s degradome predictions across different cell lines identify a validated stabilizing ARE [64], the deletion of which increases the overall degradation predictions of the transcript.

**Supp Fig. 8:**
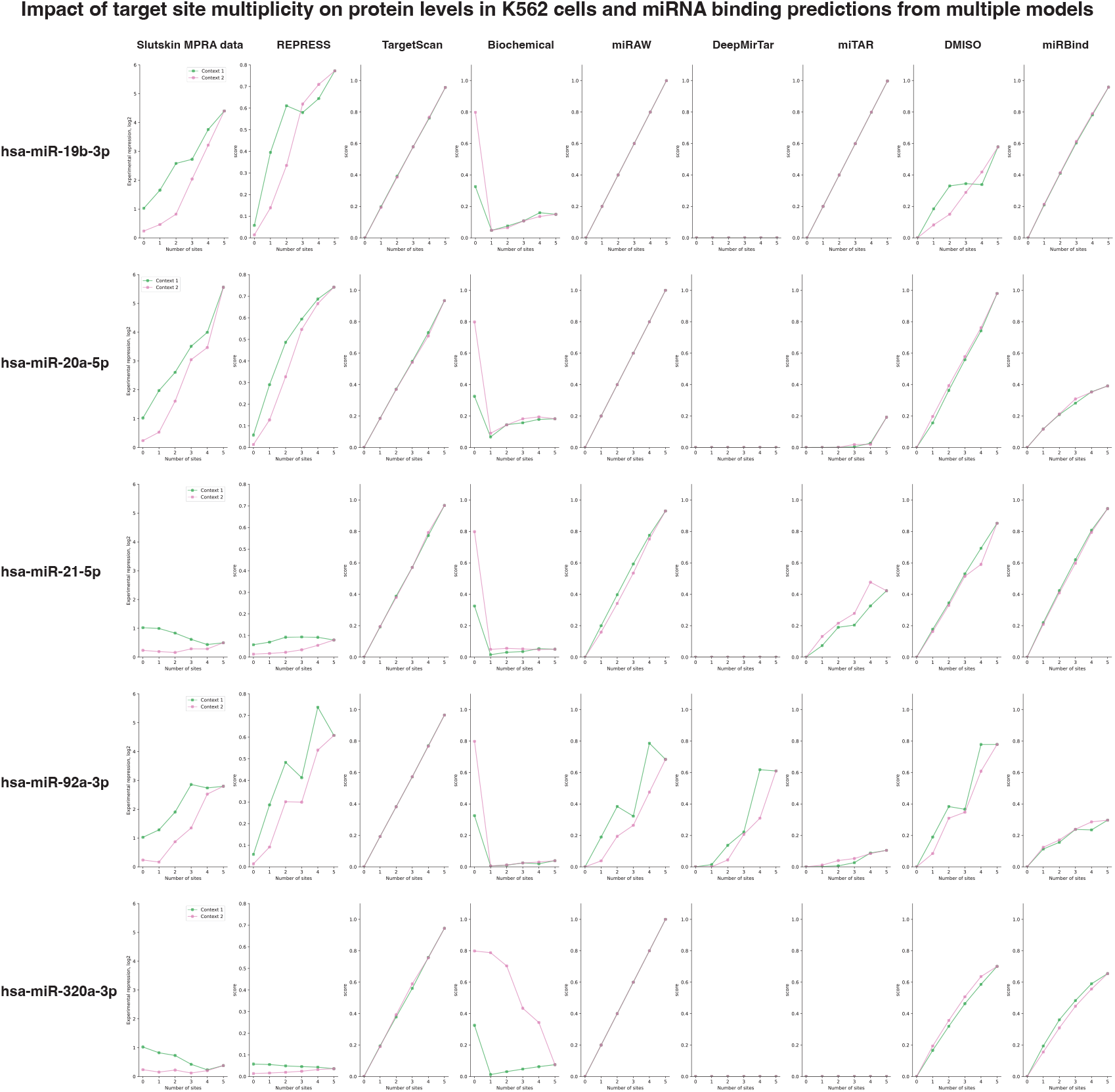
Matrix of experimental repression and model predictions for a subset of sequences designed to interrogate the impact of miRNA target multiplicity on repression and screened in an MPRA from [51]. For each plot, average values for all sequences with the same number of target sites and the same surrounding sequence context is shown. The first column shows the experimentally observed repression. The second column shows corresponding REPRESS predictions. Subsequent columns show predictions made by existing models for comparison. Biochemical model fold-change predictions were multiplied by −1 so as to correspond to repression. Each row corresponds to a different miRNA assessed. All plots in the same column share the same y-axis scale to allow comparisons across miRNAs. No additional normalization was performed for experimental repression and REPRESS prediction values. All other model predictions were scaled to the maximum value predicted by each model.

**Supp Fig. 9:**
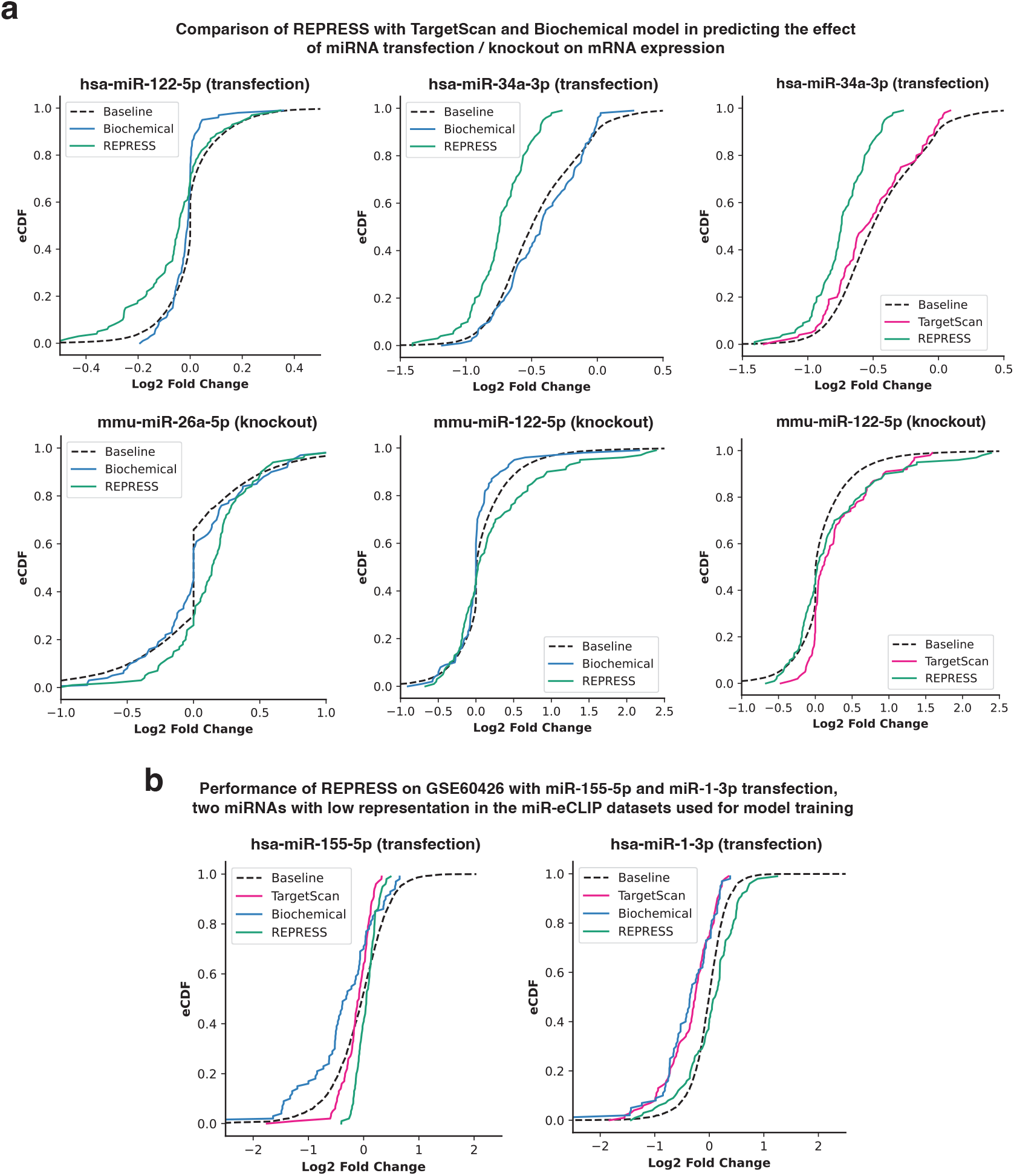
**(a):** Performance comparison of REPRESS, TargetScan and Biochemical on identifying transcripts up-regulated / repressed from miRNA knockout / transfection datasets. The figure illustrates eCDF plots of the fold changes for top predicted transcripts (N = 100) using REPRESS, TargetScan and Biochemical from several different miRNA knockout / transfection datasets. REPRESS outperforms both TargetScan and the Biochemical model on each of the 4 transfection / knockout datasets. **(b):** Performance of REPRESS, TargetScan and Biochemical model on GSE60426 miRNA transfection datasets of hsa-miR-1 and hsa-miR-155. Both these miRNAs are not well represented in our miR-eCLIP datasets with very low peak frequency. TargetScan and Biochemical model outperform REPRESS for miRNAs not well represented in our dataset.

**Supp Fig. 10:**
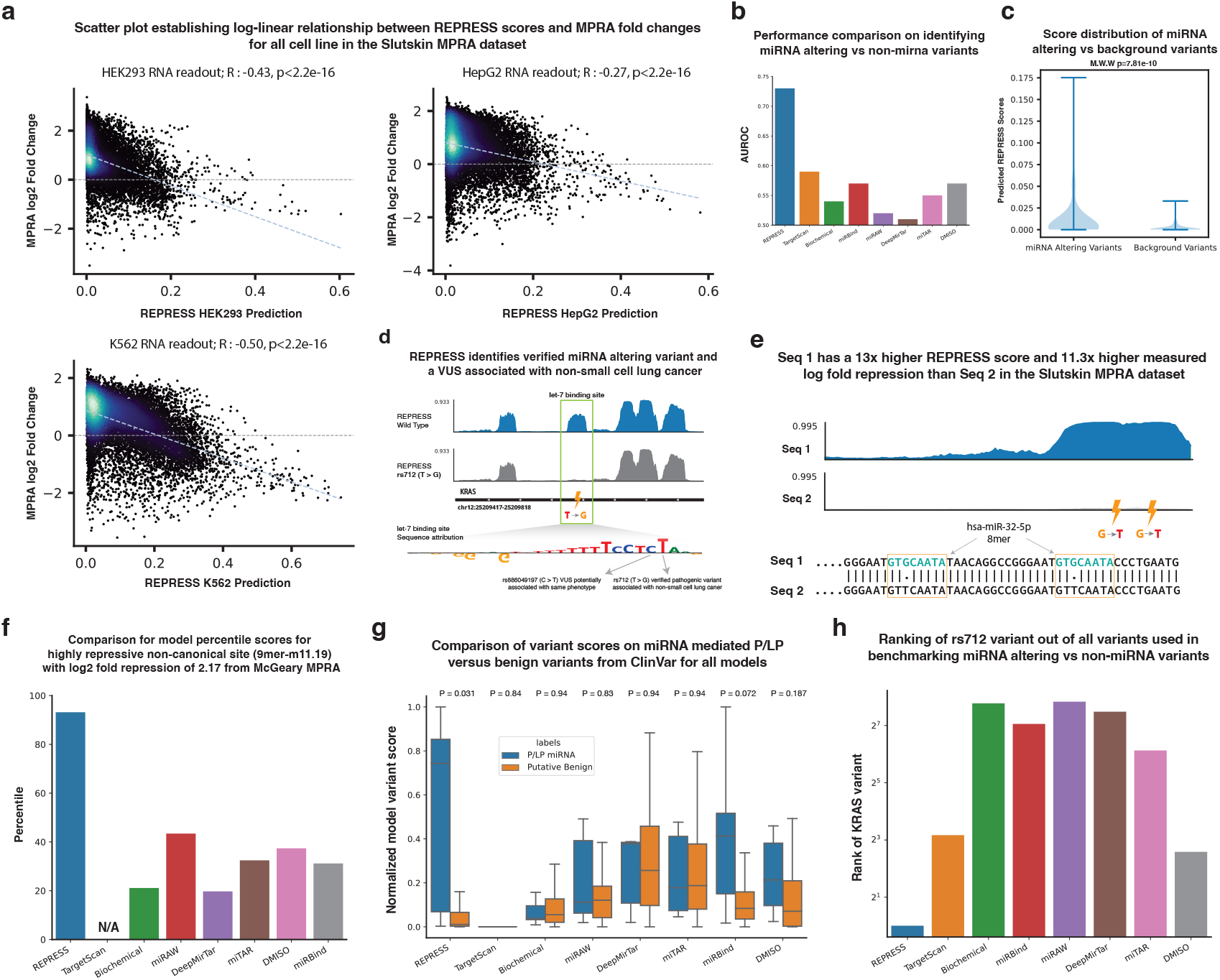
**(a):** Plot showing REPRESS predictions on MPRA sequences from the Slutskin MPRA and their corresponding fold changes. We establish a log-linear relationship between REPRESS predicted scores and corresponding fold change from the MPRA dataset. **(b):** REPRESS outperforms all baseline models in predicting validated miRNA altering from non-miRNA background variants. **(c):** Distribution of REPRESS scores for miRNA altering and background variants (*P* = 7.8 × 10^−10^). **(d):** Track visualization of REPRESS capturing the effect of a verified miRNA binding site disrupting variant (rs712). rs712 is associated with increased risk of non-small cell lung cancer due to the disruption of the let-7 miRNA binding site in the 3’ UTR of the KRAS gene. The plot also highlights the presence of a VUS (rs886049197) beside the verified variant that also disrupts let-7 binding and could potentially confer KRAS upregulation through a similar mechanism. **(e):** Track plot illustrating REPRESS’s miRNA predictions on two similar sequences from the Slutskin MPRA dataset. The two sequences only differ with G¿T mutations of two locations. Seq 1 has a 13x higher REPRESS prediction and 11.3x higher log fold change than Seq 2. This illustrates REPRESS’s sensitivity to small changes in the input sequence outside the distribution it was trained on. **(f):** Percentile comparison of all miRNA models on predicting a highly repressive non-canonical binding site (9mer.m11-19) with log2 fold repression of 2.17 from the McGeary MPRA dataset. **(g):** Box plot comparing normalized variant scores for all miRNA models on miRNA mediated P/LP variants from ClinVar versus benign variants. The Benjamini-Hochberg corrected P values for each model are shown on top for each model. **(h):** Comparison of the ranking of the rs712 pathogenic variant in KRAS for all models across all the variants (n=274) used to benchmark predicting validated miRNA altering versus background variants from GnomAD.

**Supp Fig. 11:**
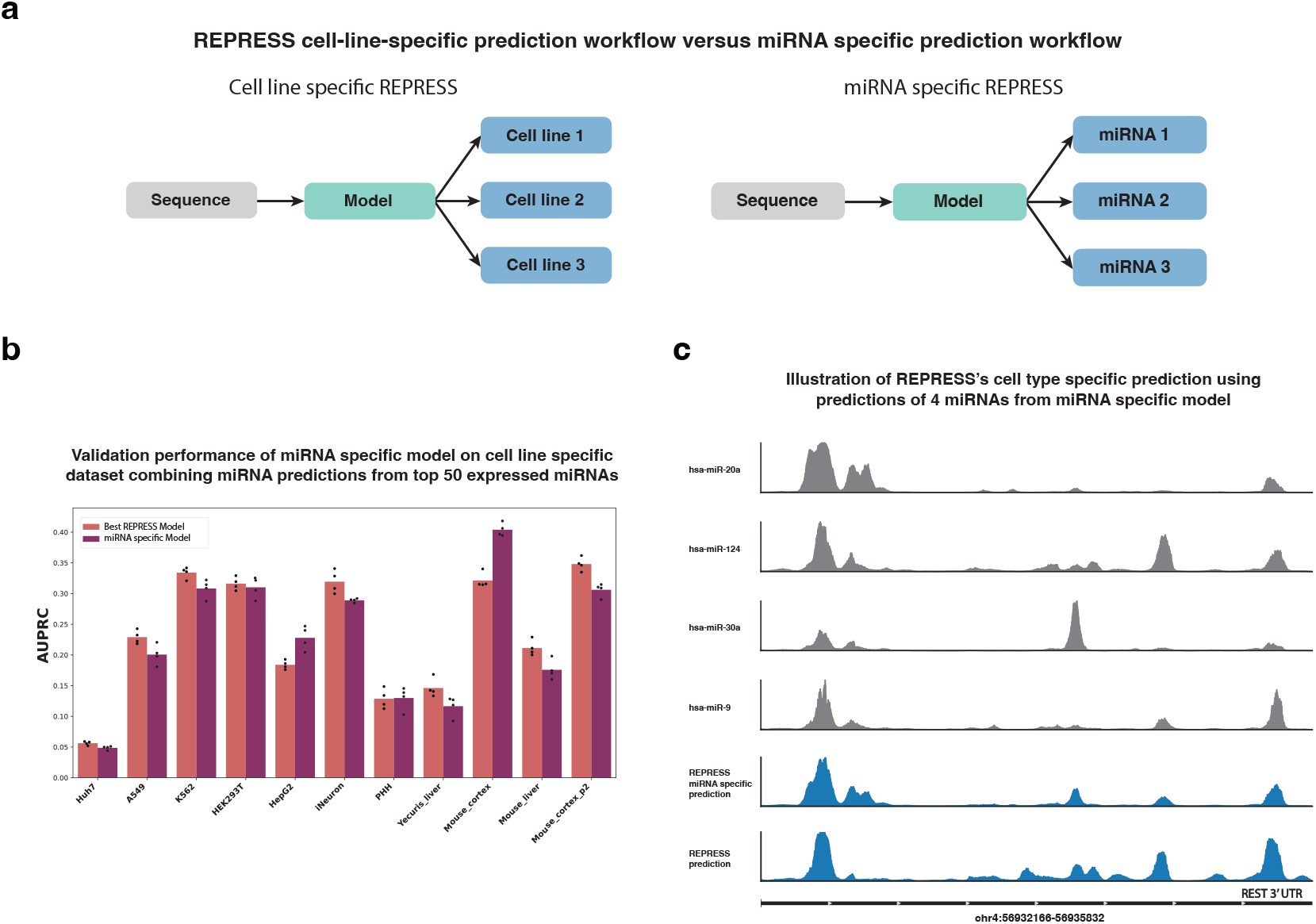
**(a):** a, Illustration of prediction workflows of cell-line-specific and miRNA-specific formulations of REPRESS. The miRNA specific formulation was trained to predict the binding of 339 individual pre-miRNAs (binding for both the 3p and 5p strands of the miRNA), each with at least 500 binding different binding sites across our miR-eCLIP datasets. **(b):** The performance of the miRNA-specific model on the cell-line-specific validation set by combining the predictions from the top 50 miRNAs expressed in that cell line. The miRNA expression weighted average was taken across the predictions of the top 50 miRNAs to get the prediction track for that corresponding cell line. The miRNA specific model maintains 97% of the average AURPC as that of the cell-line-specific version of REPRESS. **(c):** Track illustration of cell-line-specific REPRESS prediction and combining 4 different miRNA specific predictions for miRNAs expressed in iNeuron cells for the 3’ UTR of the REST gene.

## 7 Supplementary Tables

**Supp. Table 1:**
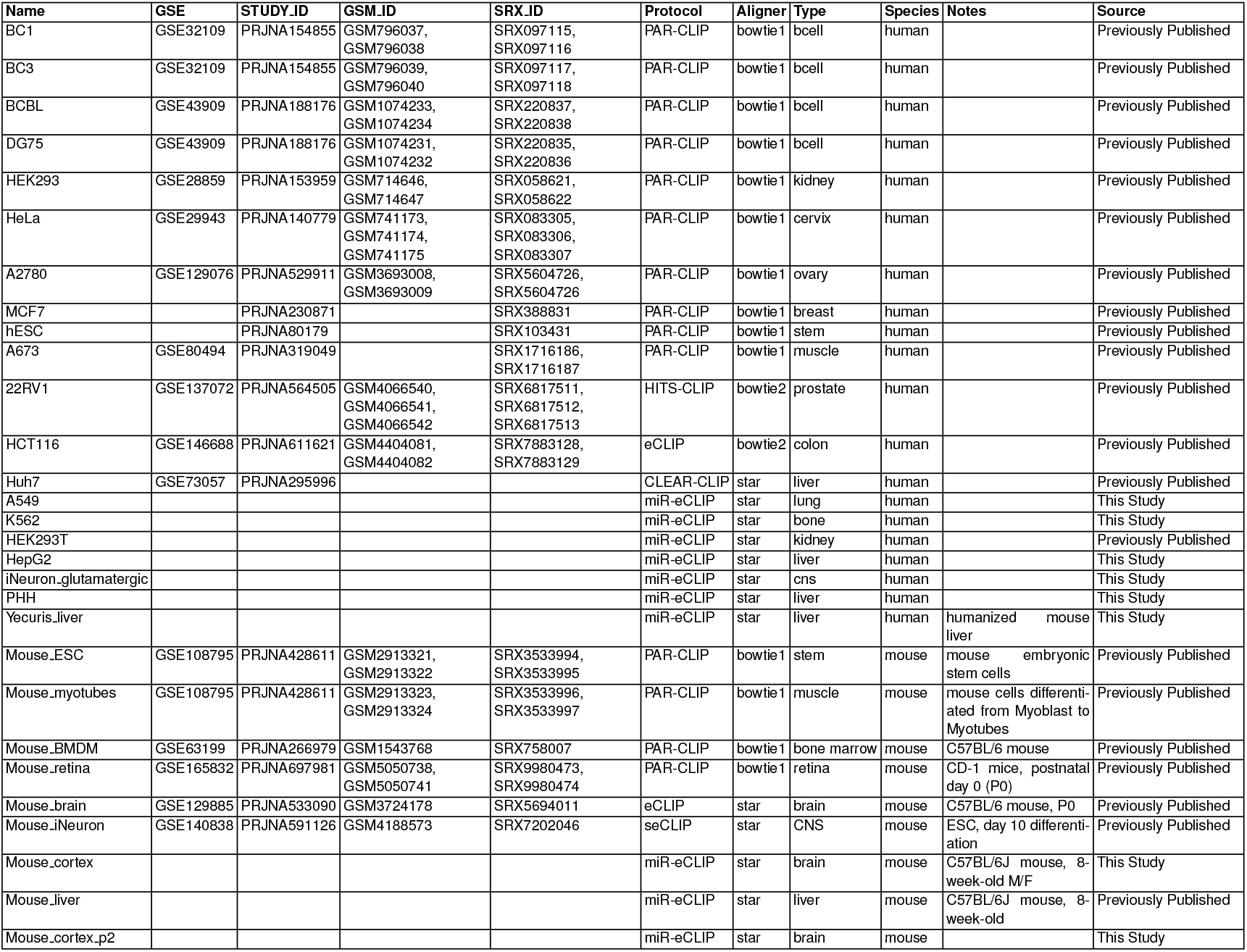
Published datasets and datasets presented in this study used to train the cell type-specific miRNA binding components of REPRESS.

**Supp. Table 2:**
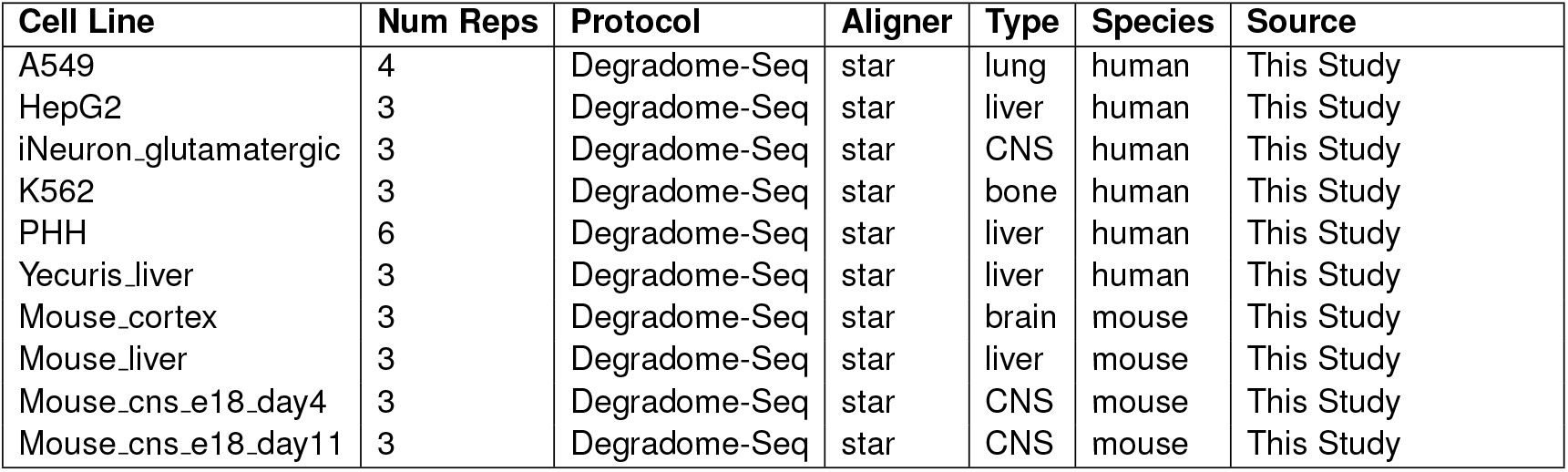
Published datasets and datasets presented in this study used to train the cell type-specific mRNA degradation components of REPRESS.

**Supp. Table 3:**
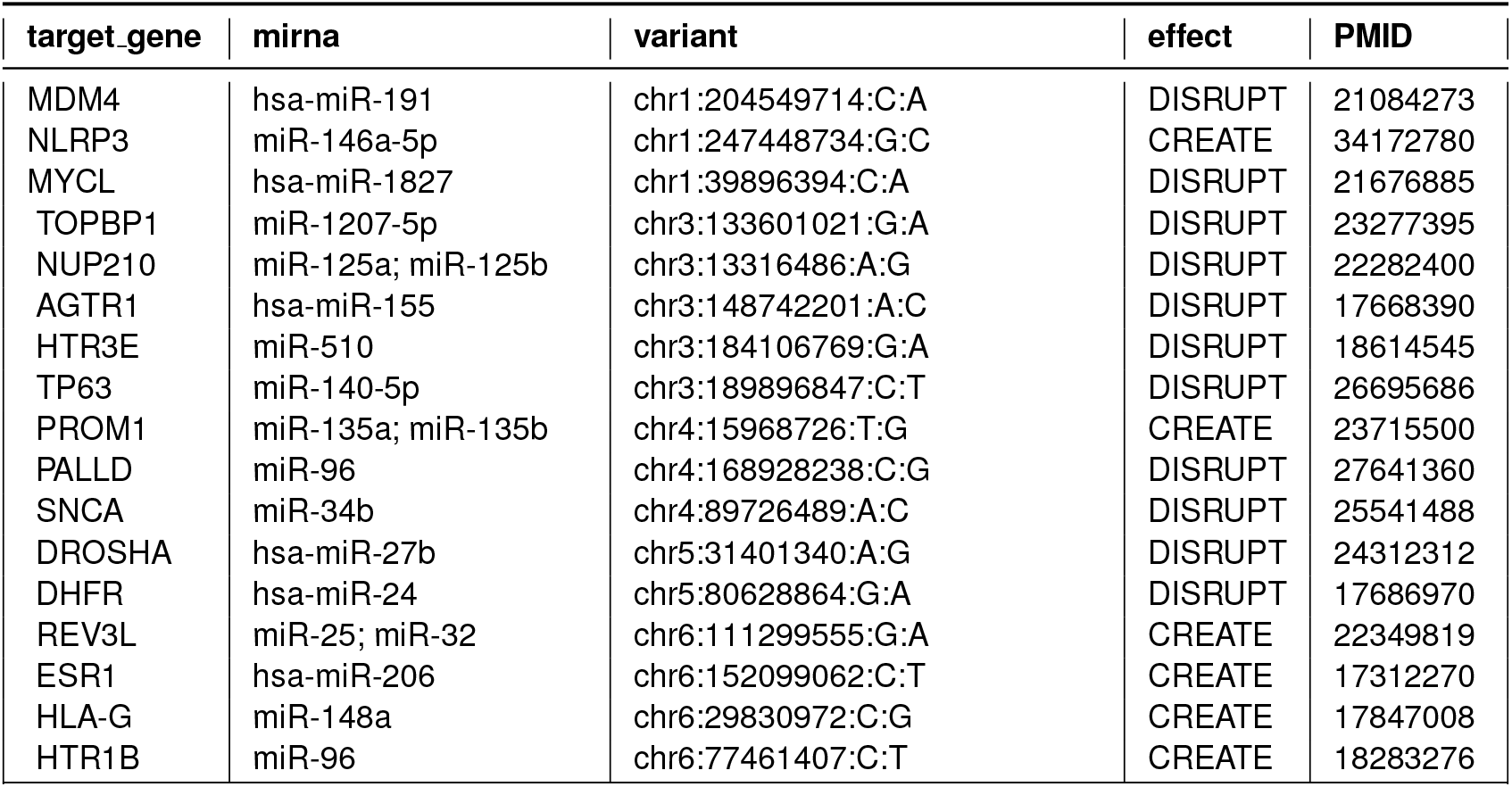
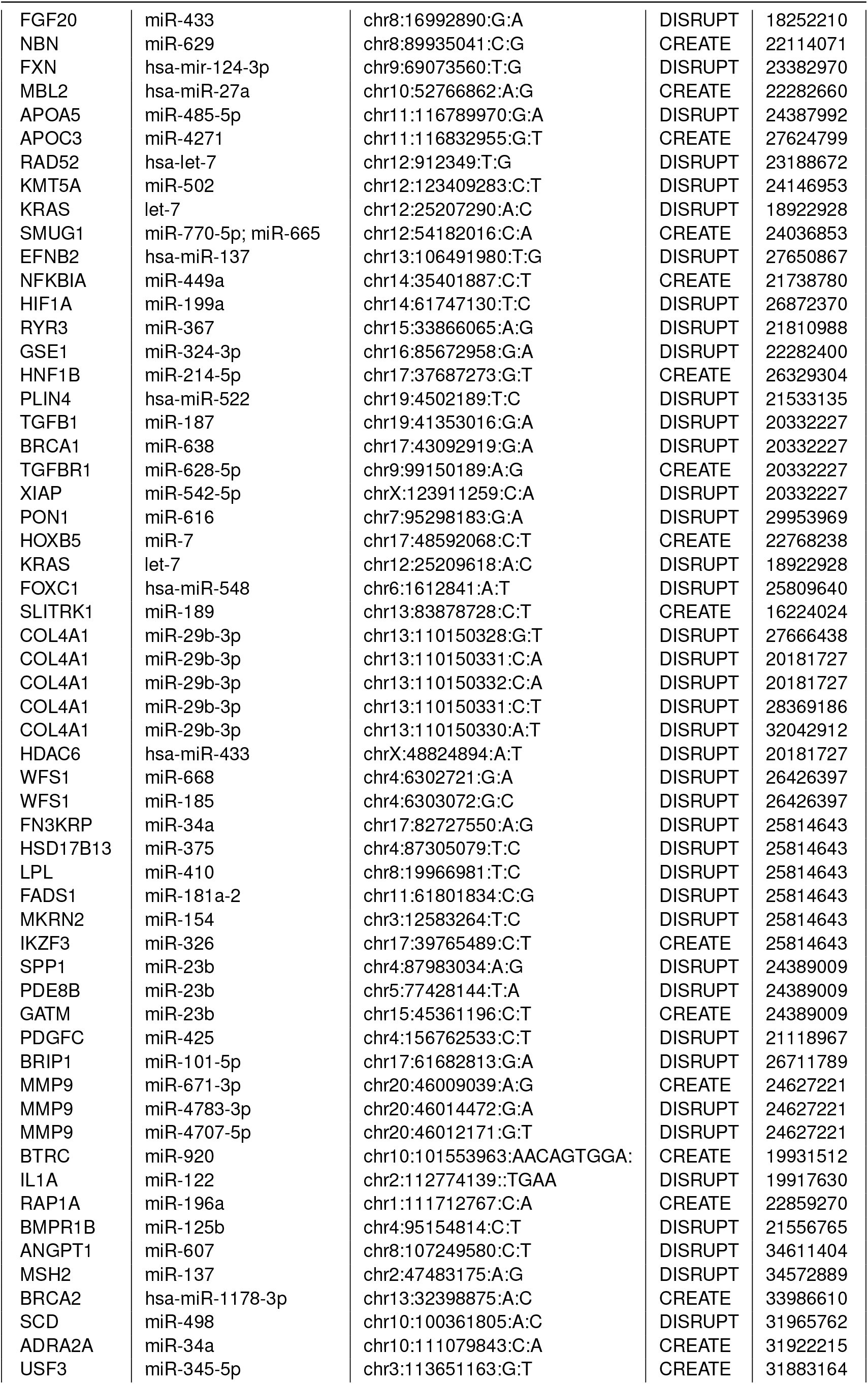
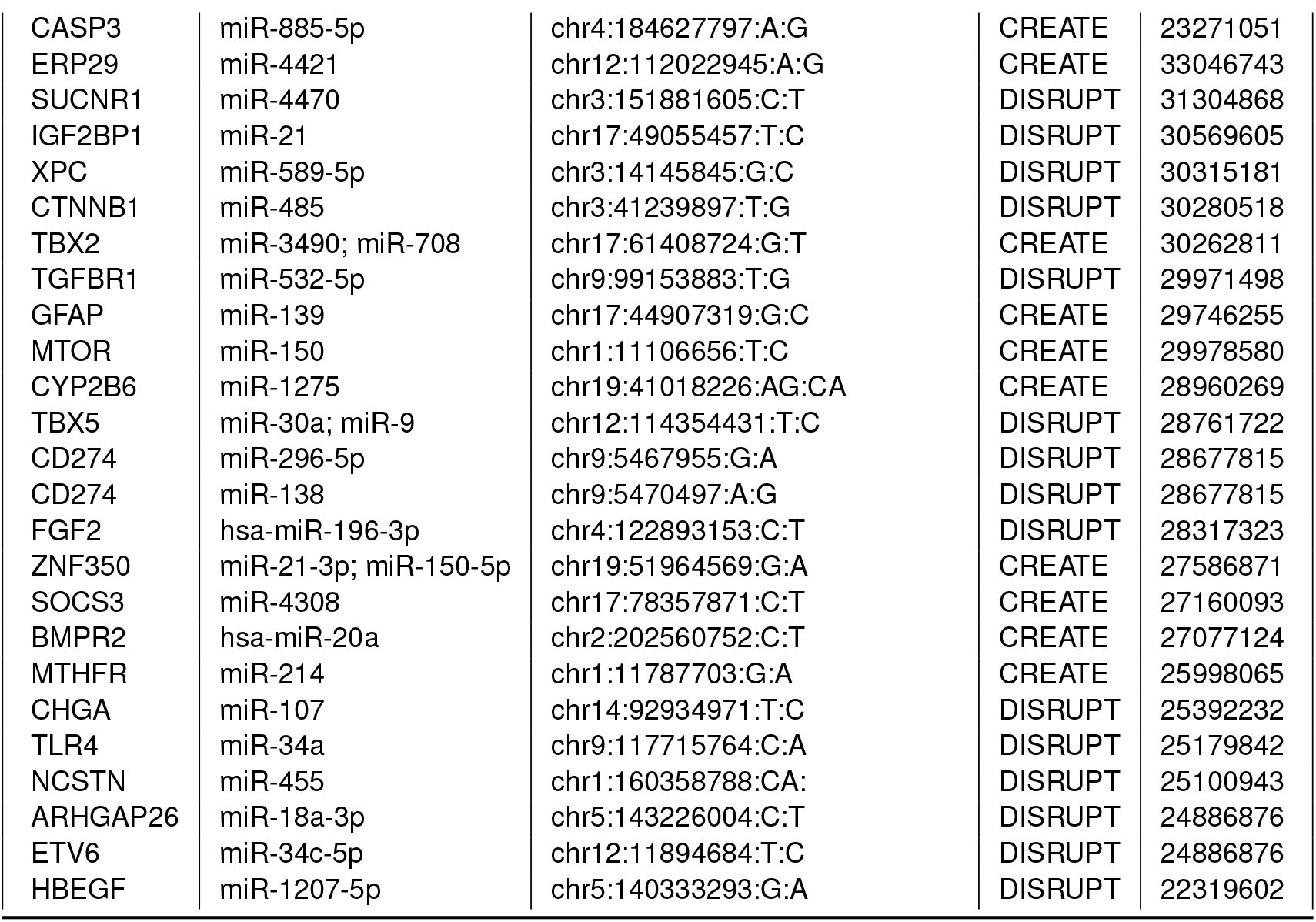
List of curated miRNA binding altering variants.

**Supp. Table 4:**
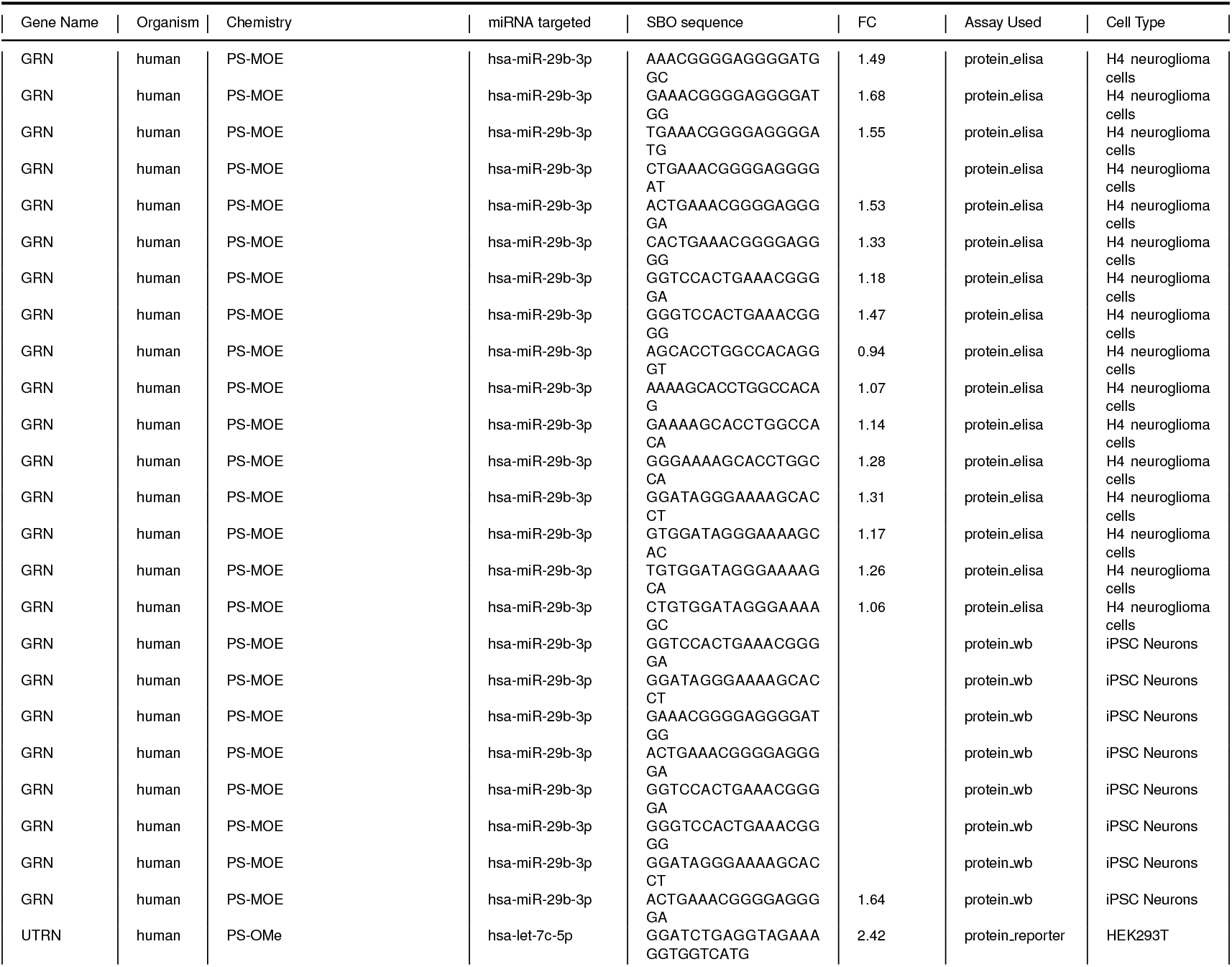
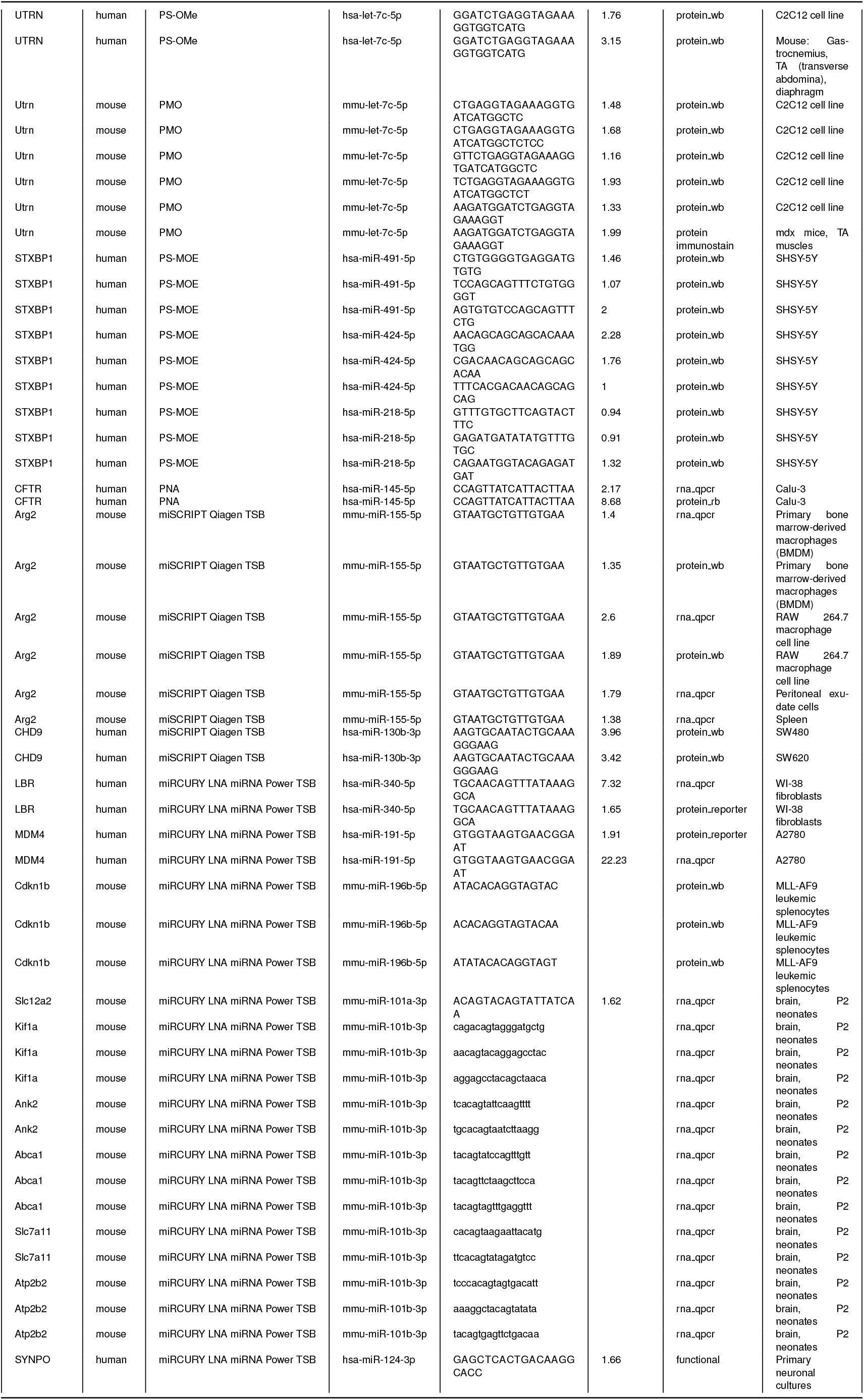
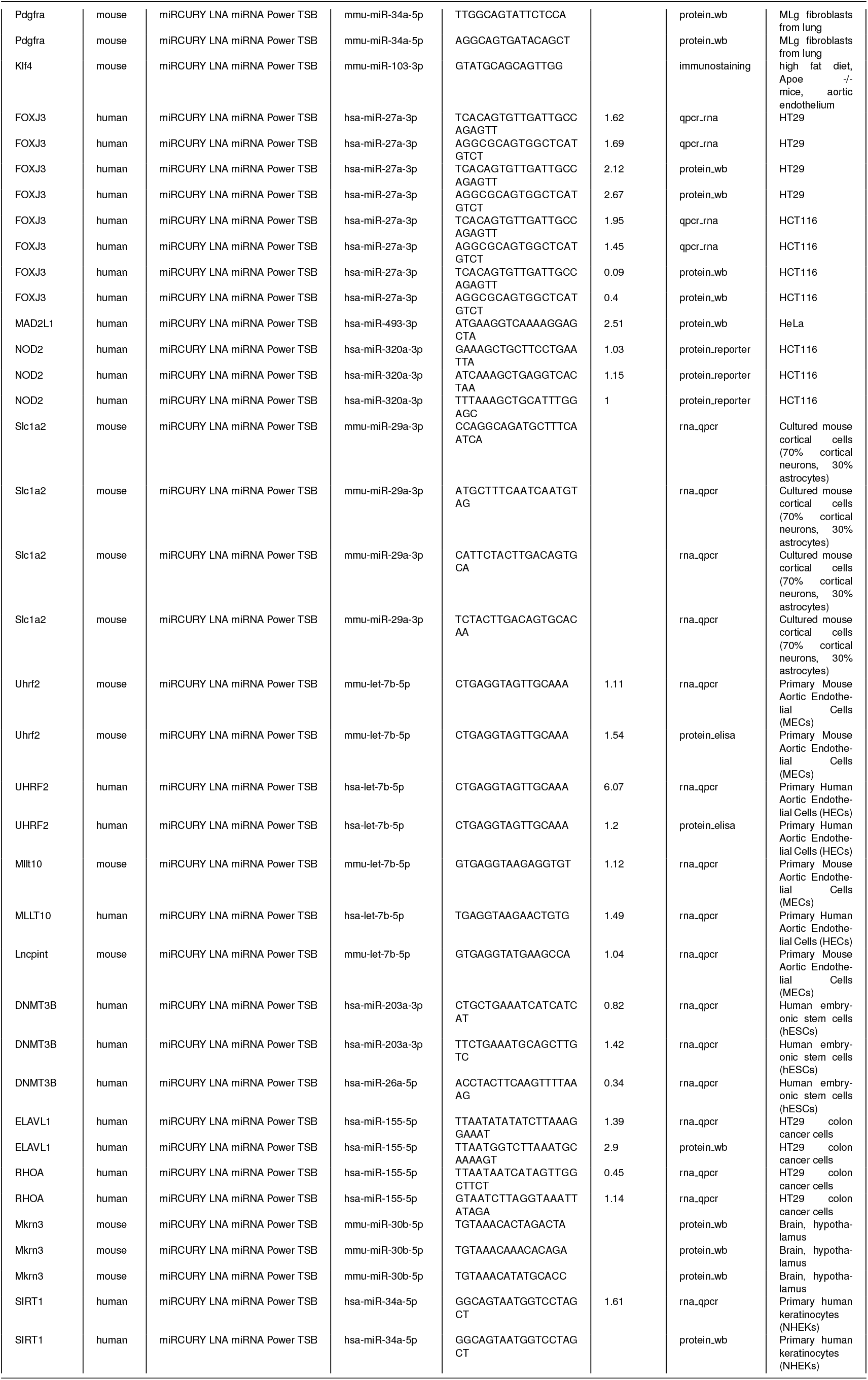
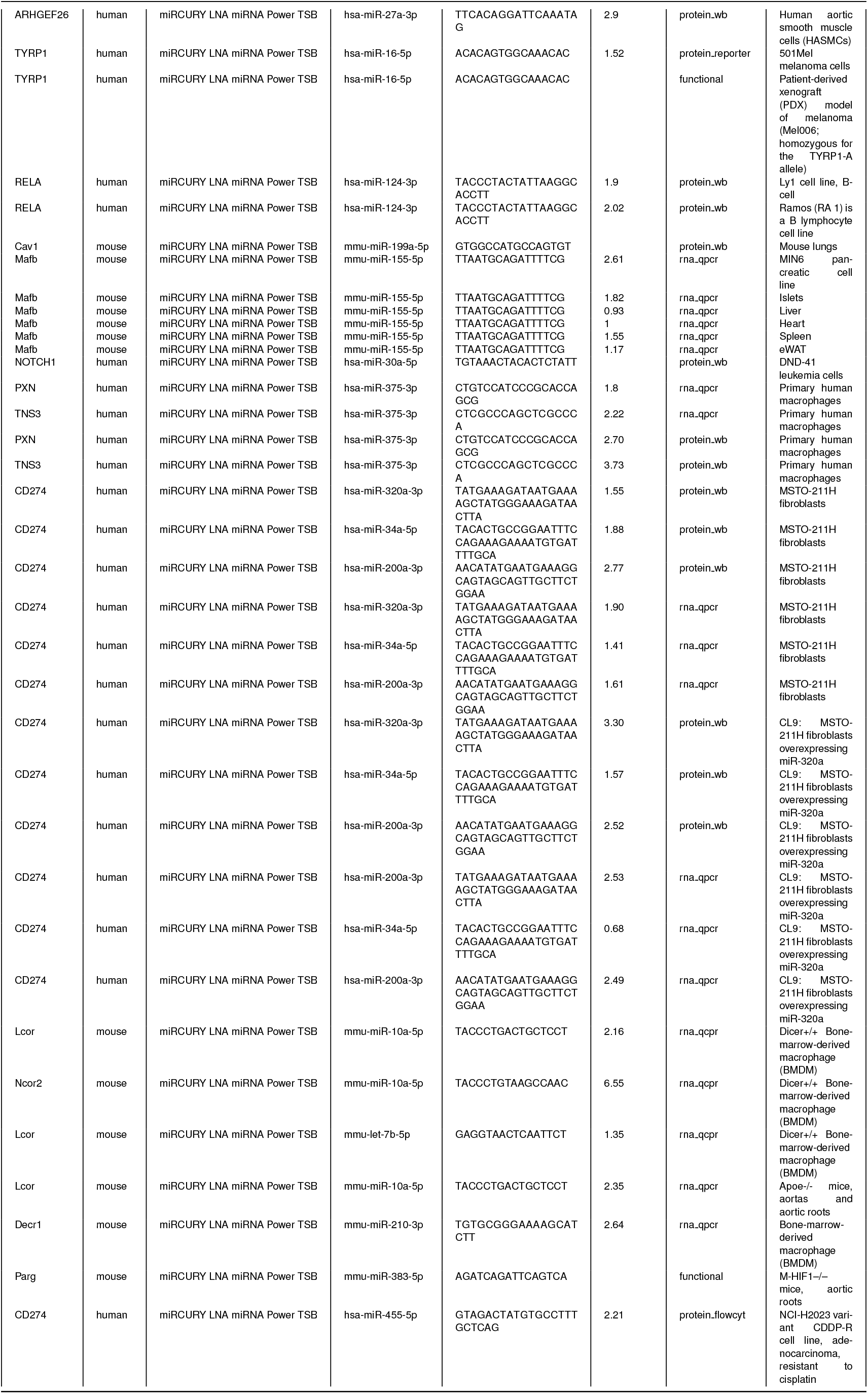
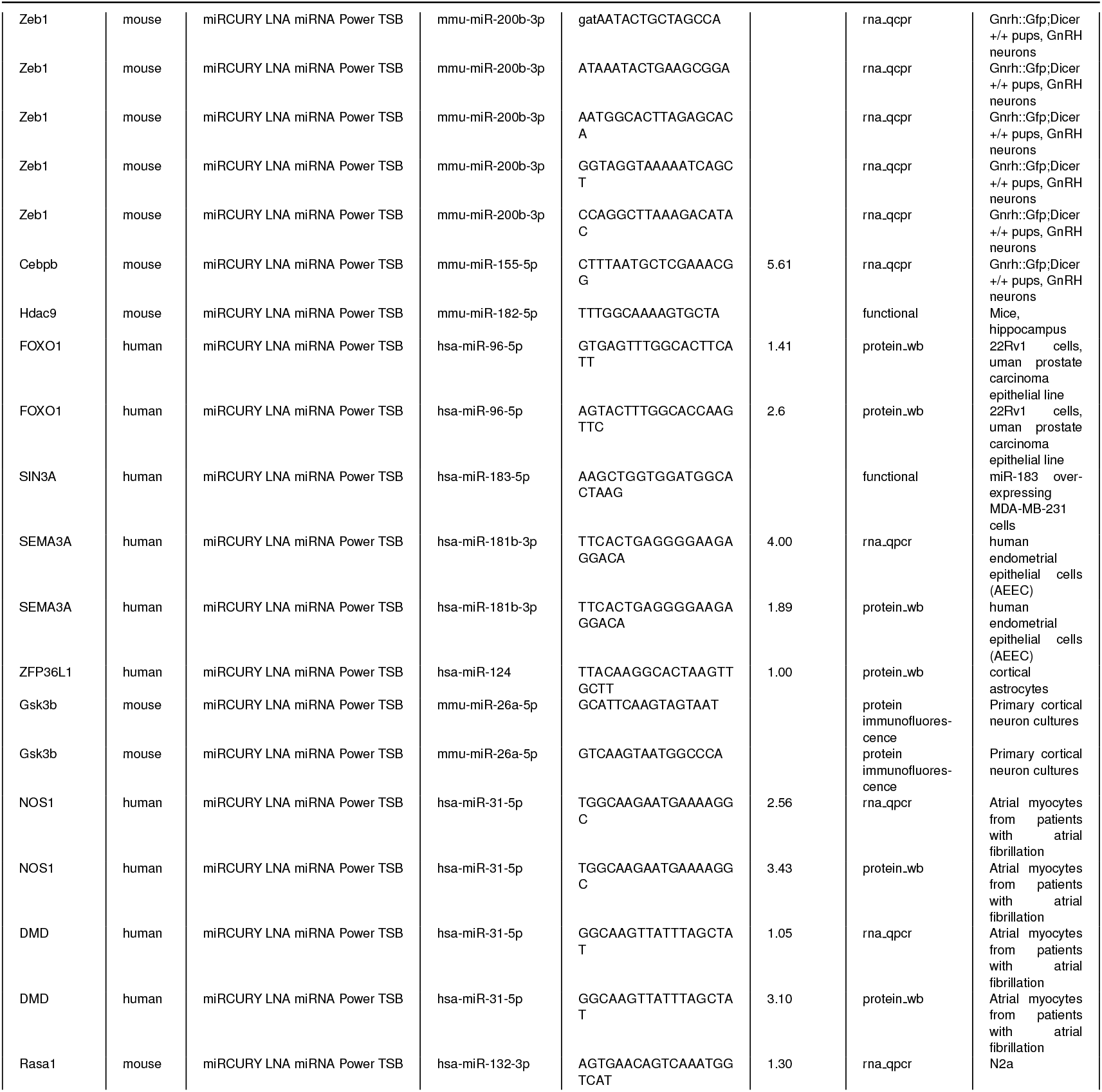
Curated literature SBO data detailing the published miRNA fold changes (FCs) across a variety of chemistries, organisms, and experimental protocols analyzed in this study. This comprehensive dataset includes information on gene names, miRNA targets, SBO sequences, assay types, and cell types corresponding to REPRESS head.

**Supp. Table 5:**
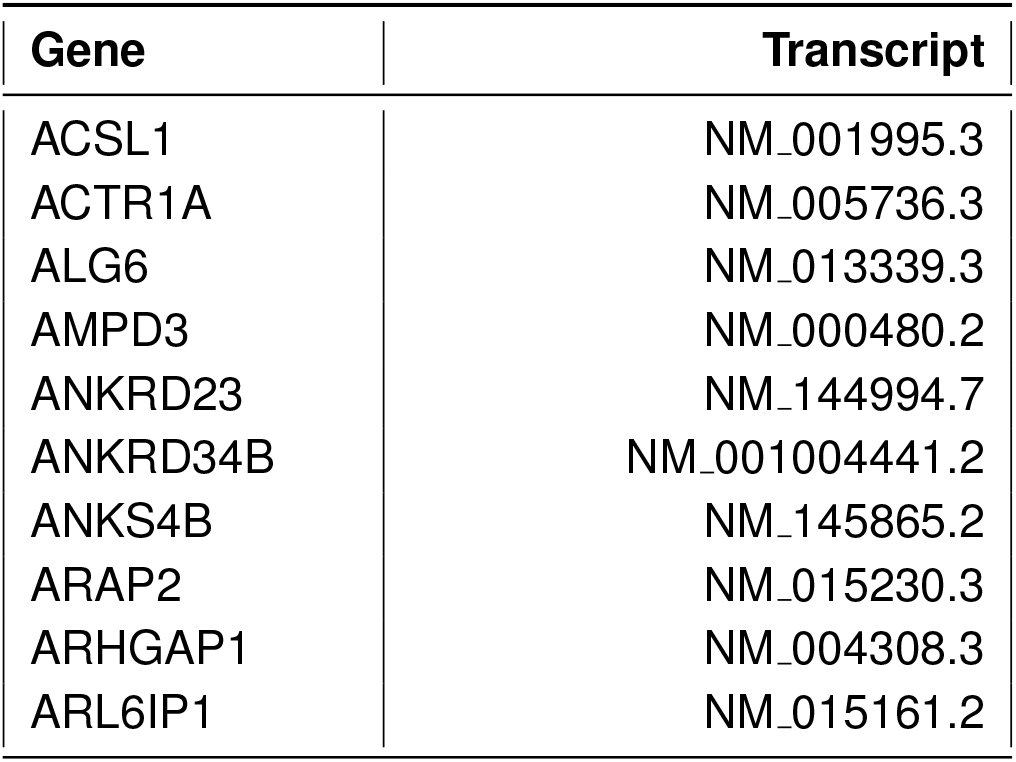
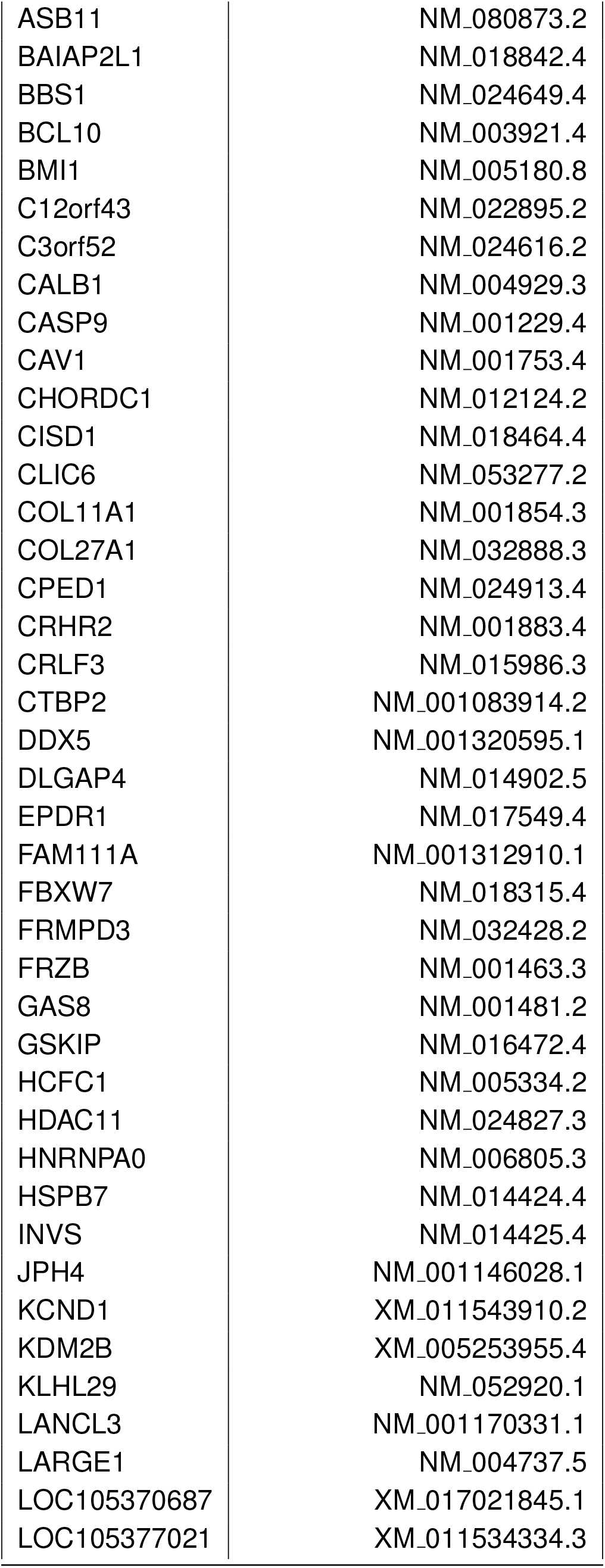
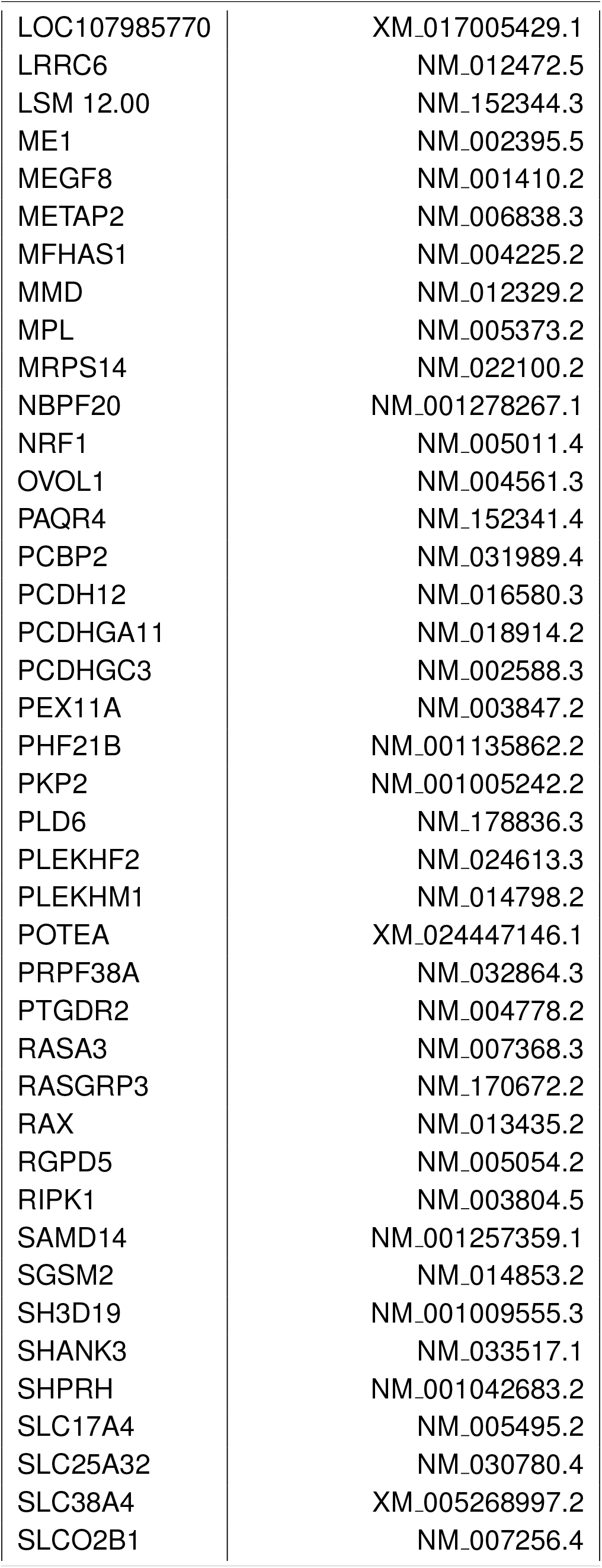
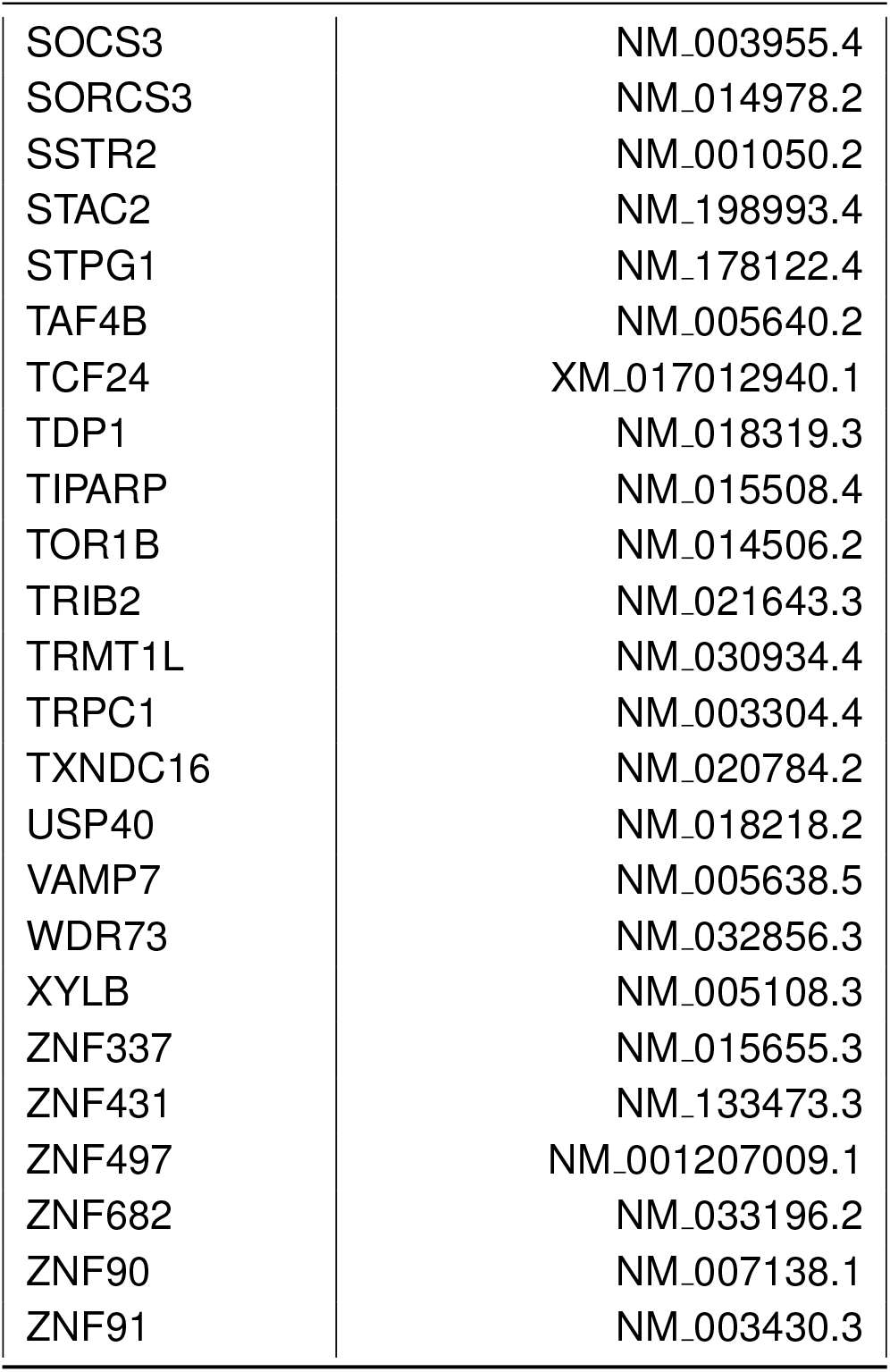
Transcripts used for the mRNA design analysis via in silico mutagenesis (ISM) and progressive edits with stability measured by Saluki. REPRESS was utilized to incorporate edits into native 3’ UTRs with ISM to minimize predicted probability of miRNA binding and degradation.

| Gene | Transcript |
| --- | --- |
| ACSL1 | NM_001995.3 |
| ACTR1A | NM_005736.3 |
| ALG6 | NM_013339.3 |
| AMPD3 | NM_000480.2 |
| ANKRD23 | NM_144994.7 |
| ANKRD34B | NM_001004441.2 |
| ANKS4B | NM_145865.2 |
| ARAP2 | NM_015230.3 |
| ARHGAP1 | NM_004308.3 |
| ARL6IP1 | NM_015161.2 |

| **Gene** | **Transcript** |
| --- | --- |
| ASB11 | NM_080873.2 |
| BAIAP2L1 | NM_018842.4 |
| BBS1 | NM_024649.4 |
| BCL10 | NM_003921.4 |
| BMI1 | NM_005180.8 |
| C12orf43 | NM_022895.2 |
| C3orf52 | NM_024616.2 |
| CALB1 | NM_004929.3 |
| CASP9 | NM_001229.4 |
| CAV1 | NM_001753.4 |
| CHORDC1 | NM_012124.2 |
| CISD1 | NM_018464.4 |
| CLIC6 | NM_053277.2 |
| COL11A1 | NM_001854.3 |
| COL27A1 | NM_032888.3 |
| CPED1 | NM_024913.4 |
| CRHR2 | NM_001883.4 |
| CRLF3 | NM_015986.3 |
| CTBP2 | NM_001083914.2 |
| DDX5 | NM_001320595.1 |
| DLGAP4 | NM_014902.5 |
| EPDR1 | NM_017549.4 |
| FAM111A | NM_001312910.1 |
| FBXW7 | NM_018315.4 |
| FRMPD3 | NM_032428.2 |
| FRZB | NM_001463.3 |
| GAS8 | NM_001481.2 |
| GSKIP | NM_016472.4 |
| HCFC1 | NM_005334.2 |
| HDAC11 | NM_024827.3 |
| HNRNPA0 | NM_006805.3 |
| HSPB7 | NM_014424.4 |
| INVS | NM_014425.4 |
| JPH4 | NM_001146028.1 |
| KCND1 | XM_011543910.2 |
| KDM2B | XM_005253955.4 |
| KLHL29 | NM_052920.1 |
| LANCL3 | NM_001170331.1 |
| LARGE1 | NM_004737.5 |
| LOC105370687 | XM_017021845.1 |
| LOC105377021 | XM_011534334.3 |

| **Gene** | **Transcript** |
| --- | --- |
| LOC107985770 | XM_017005429.1 |
| LRRC6 | NM_012472.5 |
| LSM 12.00 | NM_152344.3 |
| ME1 | NM_002395.5 |
| MEGF8 | NM_001410.2 |
| METAP2 | NM_006838.3 |
| MFHAS1 | NM_004225.2 |
| MMD | NM_012329.2 |
| MPL | NM_005373.2 |
| MRPS14 | NM_022100.2 |
| NBPF20 | NM_001278267.1 |
| NRF1 | NM_005011.4 |
| OVOL1 | NM_004561.3 |
| PAQR4 | NM_152341.4 |
| PCBP2 | NM_031989.4 |
| PCDH12 | NM_016580.3 |
| PCDHGA11 | NM_018914.2 |
| PCDHGC3 | NM_002588.3 |
| PEX11A | NM_003847.2 |
| PHF21B | NM_001135862.2 |
| PKP2 | NM_001005242.2 |
| PLD6 | NM_178836.3 |
| PLEKHF2 | NM_024613.3 |
| PLEKHM1 | NM_014798.2 |
| POTEA | XM_024447146.1 |
| PRPF38A | NM_032864.3 |
| PTGDR2 | NM_004778.2 |
| RASA3 | NM_007368.3 |
| RASGRP3 | NM_170672.2 |
| RAX | NM_013435.2 |
| RGPD5 | NM_005054.2 |
| RIPK1 | NM_003804.5 |
| SAMD14 | NM_001257359.1 |
| SGSM2 | NM_014853.2 |
| SH3D19 | NM_001009555.3 |
| SHANK3 | NM_033517.1 |
| SHPRH | NM_001042683.2 |
| SLC17A4 | NM_005495.2 |
| SLC25A32 | NM_030780.4 |
| SLC38A4 | XM_005268997.2 |
| SLCO2B1 | NM_007256.4 |
| SOCS3 | NM_003955.4 |
| SORCS3 | NM_014978.2 |
| SSTR2 | NM_001050.2 |
| STAC2 | NM_198993.4 |
| STPG1 | NM_178122.4 |
| TAF4B | NM_005640.2 |
| TCF24 | XM_017012940.1 |
| TDP1 | NM_018319.3 |
| TIPARP | NM_015508.4 |
| TOR1B | NM_014506.2 |
| TRIB2 | NM_021643.3 |
| TRMT1L | NM_030934.4 |
| TRPC1 | NM_003304.4 |
| TXNDC16 | NM_020784.2 |
| USP40 | NM_018218.2 |
| VAMP7 | NM_005638.5 |
| WDR73 | NM_032856.3 |
| XYLB | NM_005108.3 |
| ZNF337 | NM_015655.3 |
| ZNF431 | NM_133473.3 |
| ZNF497 | NM_001207009.1 |
| ZNF682 | NM_033196.2 |
| ZNF90 | NM_007138.1 |
| ZNF91 | NM_003430.3 |

## References

[1] Dassi, E. Handshakes and fights: the regulatory interplay of rna-binding proteins. Frontiers in molecular biosciences 4, 67 (2017).

[2] Halbeisen, R. E., Galgano, A., Scherrer, T. & Gerber, A. Post-transcriptional gene regulation: from genome-wide studies to principles. Cellular and molecular life sciences 65, 798–813 (2008).

[3] Lewis, B. P., Burge, C. B. & Bartel, D. P. Conserved seed pairing, often flanked by adenosines, indicates that thousands of human genes are microrna targets. cell 120, 15–20 (2005).

[4] Bartel, D. P. Micrornas: target recognition and regulatory functions. cell 136, 215–233 (2009).

[5] MacFarlane, L.-A. & R Murphy, P. Microrna: biogenesis, function and role in cancer. Current genomics 11, 537–561 (2010).

[6] Xu, K., Lin, J., Zandi, R., Roth, J. & Ji, L. Microrna-mediated target mrna cleavage and 3-uridylation in human cells. sci rep 6, 30242 (2016).

[7] Sheu-Gruttadauria, J., Xiao, Y., Gebert, L. F. & MacRae, I. J. Beyond the seed: structural basis for supplementary micro rna targeting by human argonaute2. The EMBO journal 38, e101153 (2019).

[8] Castello, A. et al. Insights into rna biology from an atlas of mammalian mrna-binding proteins. Cell 149, 1393–1406 (2012).

[9] Baltz, A. G. et al. The mrna-bound proteome and its global occupancy profile on protein-coding transcripts. Molecular cell 46, 674–690 (2012).

[10] Alles, J. et al. An estimate of the total number of true human mirnas. Nucleic acids research 47, 3353–3364 (2019).

[11] Mukherjee, N. et al. Deciphering human ribonucleoprotein regulatory networks. Nucleic acids research 47, 570–581 (2019).

[12] Gerstberger, S., Hafner, M. & Tuschl, T. A census of human rna-binding proteins. Nature Reviews Genetics 15, 829–845 (2014).

[13] Agarwal, V., Bell, G. W., Nam, J.-W. & Bartel, D. P. Predicting effective microrna target sites in mammalian mrnas. elife 4, e05005 (2015).

[14] McGeary, S. E. et al. The biochemical basis of microrna targeting efficacy. Science 366, eaav1741 (2019).

[15] Hogan, D. J., Riordan, D. P., Gerber, A. P., Herschlag, D. & Brown, P. O. Diverse rna-binding proteins interact with functionally related sets of rnas, suggesting an extensive regulatory system. PLoS biology 6, e255 (2008).

[16] Barreau, C., Paillard, L. & Osborne, H. B. Au-rich elements and associated factors: are there unifying principles? Nucleic acids research 33, 7138–7150 (2005).

[17] Wen, M., Cong, P., Zhang, Z., Lu, H. & Li, T. Deepmirtar: a deep-learning approach for predicting human mirna targets. Bioinformatics 34, 3781–3787 (2018).

[18] Gu, T., Zhao, X., Barbazuk, W. B. & Lee, J.-H. mitar: a hybrid deep learning-based approach for predicting mirna targets. BMC bioinformatics 22, 1–16 (2021).

[19] Klimentová, E. et al. mirbind: A deep learning method for mirna binding classification. Genes 13, 2323 (2022).

[20] Talukder, A., Zhang, W., Li, X. & Hu, H. A deep learning method for mirna/isomir target detection. Scientific Reports 12, 10618 (2022).

[21] Sood, P., Krek, A., Zavolan, M., Macino, G. & Rajewsky, N. Cell-type-specific signatures of micrornas on target mrna expression. Proceedings of the National Academy of Sciences 103, 2746–2751 (2006).

[22] Nam, J.-W. et al. Global analyses of the effect of different cellular contexts on microrna targeting. Molecular cell 53, 1031–1043 (2014).

[23] Nowakowski, T. J. et al. Regulation of cell-type-specific transcriptomes by microrna networks during human brain development. Nature neuroscience 21, 1784–1792 (2018).

[24] Addo-Quaye, C., Miller, W. & Axtell, M. J. Cleaveland: a pipeline for using degradome data to find cleaved small rna targets. Bioinformatics 25, 130–131 (2009).

[25] Folkes, L. et al. Paresnip: a tool for rapid genome-wide discovery of small rna/target interactions evidenced through degradome sequencing. Nucleic acids research 40, e103–e103 (2012).

[26] Thody, J. et al. Paresnip2: a tool for high-throughput prediction of small rna targets from degradome sequencing data using configurable targeting rules. Nucleic acids research 46, 8730–8739 (2018).

[27] Bracken, C. P. et al. Global analysis of the mammalian rna degradome reveals widespread mirna-dependent and mirna-independent endonucleolytic cleavage. Nucleic acids research 39, 5658–5668 (2011).

[28] Zhang, Y. et al. Identifying cleaved and noncleaved targets of small interfering rnas and micrornas in mammalian cells by spyclip. Molecular Therapy-Nucleic Acids 22, 900–909 (2020).

[29] Won, J.-I., Shin, J., Park, S. Y., Yoon, J. & Jeong, D.-H. Global analysis of the human rna degradome reveals widespread decapped and endonucleolytic cleaved transcripts. International Journal of Molecular Sciences 21, 6452 (2020).

[30] Zhou, F. et al. Identification of micrornas and their endonucleolytic cleavaged target mrnas in colorectal cancer. BMC cancer 20, 1–15 (2020).

[31] Schmidt, S. A. et al. Identification of smg6 cleavage sites and a preferred rna cleavage motif by global analysis of endogenous nmd targets in human cells. Nucleic acids research 43, 309–323 (2015).

[32] Helwak, A., Kudla, G., Dudnakova, T. & Tollervey, D. Mapping the human mirna interactome by clash reveals frequent noncanonical binding. Cell 153, 654–665 (2013).

[33] Hejret, V. et al. Analysis of chimeric reads characterises the diverse targetome of ago2-mediated regulation. Scientific Reports 13, 22895 (2023).

[34] Manakov, S. A. et al. Scalable and deep profiling of mrna targets for individual micrornas with chimeric eclip. BioRxiv 2022–02 (2022).

[35] Karginov, F. V. et al. Diverse endonucleolytic cleavage sites in the mammalian transcriptome depend upon micrornas, drosha, and additional nucleases. Molecular cell 38, 781–788 (2010).

[36] German, M. A. et al. Global identification of microrna–target rna pairs by parallel analysis of rna ends. Nature biotechnology 26, 941–946 (2008).

[37] Addo-Quaye, C., Eshoo, T. W., Bartel, D. P. & Axtell, M. J. Endogenous sirna and mirna targets identified by sequencing of the arabidopsis degradome. Current Biology 18, 758–762 (2008).

[38] Zhang, H. et al. Linearcofold and linearcopartition: linear-time algorithms for secondary structure prediction of interacting rna molecules. Nucleic Acids Research 51, e94–e94 (2023).

[39] Van Nostrand, E. L. et al. Principles of rna processing from analysis of enhanced clip maps for 150 rna binding proteins. Genome biology 21, 1–26 (2020).

[40] Haberman, N. et al. Abundant capped rnas are derived from mrna cleavage at 3’utr g-quadruplexes. bioRxiv (2023). URL https://www.biorxiv.org/content/early/2023/06/22/2023.04.27.538568.

[41] Yekta, S., Shih, I.-h. & Bartel, D. P. Microrna-directed cleavage of hoxb8 mrna. Science 304, 594–596 (2004).

[42] Liu, Z. et al. A convnet for the 2020s, 11976–11986 (2022).

[43] Huang, H.-Y. et al. mirtarbase update 2022: an informative resource for experimentally validated mirna–target interactions. Nucleic acids research 50, D222–D230 (2022).

[44] Karagkouni, D. et al. Diana-tarbase v8: a decade-long collection of experimentally supported mirna–gene interactions. Nucleic acids research 46, D239–D245 (2018).

[45] Kaplan, J. et al. Scaling laws for neural language models. arXiv preprint arXiv:2001.08361 (2020).

[46] Hesslow, D., Zanichelli, N., Notin, P., Poli, I. & Marks, D. Rita: a study on scaling up generative protein sequence models. arXiv preprint arXiv:2205.05789 (2022).

[47] Vaswani, A. et al. Attention is all you need. Advances in neural information processing systems 30 (2017).

[48] Gu, A. & Dao, T. Mamba: Linear-time sequence modeling with selective state spaces. arXiv preprint arXiv:2312.00752 (2023).

[49] Pla, A., Zhong, X. & Rayner, S. miraw: A deep learning-based approach to predict microrna targets by analyzing whole microrna transcripts. PLoS computational biology 14, e1006185 (2018).

[50] Li, X. et al. High-resolution in vivo identification of mirna targets by halo-enhanced ago2 pull-down. Molecular cell 79, 167–179 (2020).

[51] Vainberg Slutskin, I., Weingarten-Gabbay, S., Nir, R., Weinberger, A. & Segal, E. Unraveling the determinants of microrna mediated regulation using a massively parallel reporter assay. Nature Communications 9, 529 (2018).

[52] McGeary, S. E., Bisaria, N., Pham, T. M., Wang, P. Y. & Bartel, D. P. Microrna 3-compensatory pairing occurs through two binding modes, with affinity shaped by nucleotide identity and position. Elife 11, e69803 (2022).

[53] Xu, H. et al. Nfix circular rna promotes glioma progression by regulating mir-34a-5p via notch signaling pathway. Frontiers in molecular neuroscience 11, 225 (2018).

[54] Bakheet, T., Hitti, E. & Khabar, K. S. A. Ared-plus: an updated and expanded database of au-rich element-containing mrnas and pre-mrnas. Nucleic acids research 46, D218–D220 (2018).

[55] Friedman, R. C., Farh, K. K.-H., Burge, C. B. & Bartel, D. P. Most mammalian mrnas are conserved targets of micrornas. Genome research 19, 92–105 (2009).

[56] Wang, X. Composition of seed sequence is a major determinant of microrna targeting patterns. Bioinformatics 30, 1377–1383 (2014).

[57] Grimson, A. et al. Microrna targeting specificity in mammals: determinants beyond seed pairing. Molecular cell 27, 91–105 (2007).

[58] Sætrom, P. et al. Distance constraints between microrna target sites dictate efficacy and cooperativity. Nucleic acids research 35, 2333–2342 (2007).

[59] Li, P. et al. Differential inhibition of target gene expression by human micrornas. Cells 8, 791 (2019).

[60] Awan, H. M. et al. Comparing two approaches of mir-34a target identification, biotinylated-mirna pulldown vs mirna overexpression. RNA biology 15, 55–61 (2018).

[61] Luna, J. M. et al. Argonaute clip defines a deregulated mir-122-bound transcriptome that correlates with patient survival in human liver cancer. Molecular cell 67, 400–410 (2017).

[62] Fujiwara, Y. et al. mir-23a/b clusters are not essential for the pathogenesis of osteoarthritis in mouse aging and post-traumatic models. Frontiers in Cell and Developmental Biology 10, 1043259 (2023).

[63] Siegel, D. A., Le Tonqueze, O., Biton, A., Zaitlen, N. & Erle, D. J. Massively parallel analysis of human 3 utrs reveals that au-rich element length and registration predict mrna destabilization. G3 12, jkab404 (2022).

[64] Díaz-Muñoz, M. D., Bell, S. E. & Turner, M. Deletion of au-rich elements within the bcl2 3 utr reduces protein expression and b cell survival in vivo. PloS one 10, e0116899 (2015).

[65] Karczewski, K. J. et al. The mutational constraint spectrum quantified from variation in 141,456 humans. Nature 581, 434–443 (2020).

[66] Verdura, E. et al. Disruption of a mi r-29 binding site leading to col4a1 upregulation causes pontine autosomal dominant microangiopathy with leukoencephalopathy. Annals of neurology 80, 741–753 (2016).

[67] Tan, Z. et al. Allele-specific targeting of micrornas to hla-g and risk of asthma. The American Journal of Human Genetics 81, 829–834 (2007).

[68] Landrum, M. J. et al. Clinvar: public archive of relationships among sequence variation and human phenotype. Nucleic acids research 42, D980–D985 (2014).

[69] Landrum, M. J. et al. Clinvar: improving access to variant interpretations and supporting evidence. Nucleic acids research 46, D1062–D1067 (2018).

[70] Chin, L. J. et al. A snp in a let-7 microrna complementary site in the kras 3 untranslated region increases non–small cell lung cancer risk. Cancer research 68, 8535–8540 (2008).

[71] Li, Z.-H. et al. A let-7 binding site polymorphism rs712 in the kras 3 utr is associated with an increased risk of gastric cancer. Tumor Biology 34, 3159–3163 (2013).

[72] Hu, H., Zhang, L., Teng, G., Wu, Y. & Chen, Y. A variant in 3-untranslated region of kras compromises its interaction with hsa-let-7g and contributes to the development of lung cancer in patients with copd. International Journal of Chronic Obstructive Pulmonary Disease 1641–1649 (2015).

[73] Sengupta, K. et al. Genome editing-mediated utrophin upregulation in duchenne muscular dystrophy stem cells. Molecular Therapy-Nucleic Acids 22, 500–509 (2020).

[74] Song, D., Zhang, Q., Zhang, H., Zhan, L. & Sun, X. Mir-130b-3p promotes colorectal cancer progression by targeting chd9. Cell Cycle 21, 585–601 (2022).

[75] Idrees, M. et al. Decreased serum pon1 arylesterase activity in familial hypercholesterolemia patients with a mutated ldlr gene. Genetics and Molecular Biology 41, 570–577 (2018).

[76] Zhao, Y. et al. Association between pon1 activity and coronary heart disease risk: a meta-analysis based on 43 studies. Molecular genetics and metabolism 105, 141–148 (2012).

[77] Beaudet, L. et al. Alphalisa immunoassays: the no-wash alternative to elisas for research and drug discovery (2008).

[78] Bielefeld-Sevigny, M. Alphalisa immunoassay platform—the “no-wash” high-throughput alternative to elisa. Assay and drug development technologies 7, 90–92 (2009).

[79] Agarwal, V. & Kelley, D. R. The genetic and biochemical determinants of mrna degradation rates in mammals. Genome Biology 23, 245 (2022).

[80] Moore, M. J. et al. mirna–target chimeras reveal mirna 3-end pairing as a major determinant of argonaute target specificity. Nature communications 6, 8864 (2015).

[81] Zhou, S. et al. Degradome sequencing reveals an integrative mirna-mediated gene interaction network regulating rice seed vigor. BMC plant biology 22, 269 (2022).

[82] Chen, S., Zhou, Y., Chen, Y. & Gu, J. fastp: an ultra-fast all-in-one fastq preprocessor. Bioinformatics 34, i884–i890 (2018).

[83] Langmead, B., Trapnell, C., Pop, M. & Salzberg, S. L. Ultrafast and memory-efficient alignment of short dna sequences to the human genome. Genome biology 10, 1–10 (2009).

[84] Corcoran, D. L. et al. Paralyzer: definition of rna binding sites from par-clip short-read sequence data. Genome biology 12, 1–16 (2011).

[85] Langmead, B. & Salzberg, S. L. Fast gapped-read alignment with bowtie 2. Nature methods 9, 357–359 (2012).

[86] Lovci, M. T. et al. Rbfox proteins regulate alternative mrna splicing through evolutionarily conserved rna bridges. Nature structural & molecular biology 20, 1434–1442 (2013).

[87] Smith, T., Heger, A. & Sudbery, I. Umi-tools: modeling sequencing errors in unique molecular identifiers to improve quantification accuracy. Genome research 27, 491–499 (2017).

[88] Martin, M. Cutadapt removes adapter sequences from high-throughput sequencing reads. EMBnet. journal 17, 10–12 (2011).

[89] Kozomara, A., Birgaoanu, M. & Griffiths-Jones, S. mirbase: from microrna sequences to function. Nucleic acids research 47, D155–D162 (2019).

[90] Dobin, A. et al. Star: ultrafast universal rna-seq aligner. Bioinformatics 29, 15–21 (2013).

[91] Kim, D., Paggi, J. M., Park, C., Bennett, C. & Salzberg, S. L. Graph-based genome alignment and genotyping with hisat2 and hisat-genotype. Nature biotechnology 37, 907–915 (2019).

[92] Liao, Y., Smyth, G. K. & Shi, W. featurecounts: an efficient general purpose program for assigning sequence reads to genomic features. Bioinformatics 30, 923–930 (2014).

[93] Rodriguez, J. M. et al. Appris: selecting functionally important isoforms. Nucleic acids research 50, D54–D59 (2022).

[94] Cao, J., Zhou, W., Steemers, F., Trapnell, C. & Shendure, J. Sci-fate characterizes the dynamics of gene expression in single cells. Nature biotechnology 38, 980–988 (2020).

[95] Schofield, J. A., Duffy, E. E., Kiefer, L., Sullivan, M. C. & Simon, M. D. Timelapse-seq: adding a temporal dimension to rna sequencing through nucleoside recoding. Nature methods 15, 221–225 (2018).

[96] Bohn, E., Lau, T. T., Wagih, O., Masud, T. & Merico, D. A curated census of pathogenic and likely pathogenic utr variants and evaluation of deep learning models for variant effect prediction. Frontiers in Molecular Biosciences 10 (2023).

